# From Splicing to Survival: Understanding the Molecular Basis of Prognostic Differences in advanced Small Intestine Neuroendocrine Tumors

**DOI:** 10.64898/2026.10.22.760955

**Authors:** Víctor García-Vioque, Alejandro García-Álvarez, María Trinidad Moreno-Montilla, Jorge Hernando, Ricardo Blázquez-Encinas, Amparo García, Carlos López-López, Paula Jimenez-Fonseca, Rocio Garcia-Carbonero, Stefania Landolfi, Laia Blanco, Anna Casteras, Ana Vivancos, Alejandro Ibáñez-Costa, Sergio Pedraza-Arevalo, Jaume Capdevila, Justo P Castaño

**Affiliations:** Department of Cell Biology, Physiology and Immunology, University of Cordoba, Cordoba, 14014, Spain; Maimonides Biomedical Research Institute of Cordoba (IMIBIC), Cordoba, 14004, Spain; Reina Sofia University Hospital, Cordoba, 14004, Spain; Medical Oncology Department, Vall d’Hebron University Hospital. Vall d’Hebron Institute of Oncology (VHIO), Barcelona, 08035, Spain; Nuclear Medicine Department, Vall d’Hebron University Hospital, Barcelona, 08035, Spain; Department of Medical Oncology, Marqués de Valdecilla University Hospital, IDIVAL, UNICAN, Santander, 39008, Spain; Department of Medical Oncology, Hospital Universitario Central de Asturias, ISPA, Oviedo, 33011, Spain; Department of Medical Oncology, Hospital Universitario 12 de Octubre, Imas12 Research Institute, Medicine Faculty, Universidad Complutense de Madrid, Madrid, 28041, Spain; Pathology Department, Vall d’Hebron University Hospital, Barcelona, 08035, Spain; General Surgery Department, Vall d’Hebron University Hospital, Barcelona, 08035, Spain; Endocrinology Department, Vall d’Hebron University Hospital, Barcelona, 08035, Spain; CIBER Fisiopatología de la Obesidad y Nutrición (CIBERobn)

**Keywords:** metastasis, motifs, small intestine neuroendocrine tumors, splicing events, survival

## Abstract

Small intestine neuroendocrine tumors (SI-NETs) represent a challenging neoplasm due to their high rate of metastasis and the limited availability of effective therapies. Recent evidence has unveiled altered splicing as an influential player in tumor progression, linked to patient prognosis, which provides a powerful source of novel biomarkers and therapeutic targets in cancer. Here, we conducted a comprehensive analysis regarding the possible connection of the spliceosomic landscape of metastatic SI-NETs with patient survival. Initial analyses revealed a distinctive pattern of splicing events between the two conditions, short (< 5 years, *n* = 17 patients), vs. long survival (≥ 5 years, *n* = 26 patients), with strong discrimination capacity. Furthermore, numerous key splicing events were associated with relevant clinical parameters in SI-NETs. The observed impairments in spliceosomic parameters led us to interrogate the integrity of the splicing machinery. This revealed striking changes in the expression of a high proportion of core spliceosome components and splicing factors between short vs. longsurvival groups, suggesting a possible mechanistic link with the distinct profile of splicing events observed. Among them, CPSF2, a core component of the CPSF complex, stands out as its higher expression is associated with worse survival and vascular and perineural invasion. Interestingly, the shifts potentially generated by this factor supported a possible role in tumor plasticity oriented to higher tumor aggressiveness and treatment resistance. In sum, this study offers the first detailed snapshot of the status of the spliceosomic landscape in SI-NETs and its molecular linkage to patient survival, thus offering a new window towards identification of novel candidate biomarkers and/or therapeutic targets.

## Introduction

Neuroendocrine tumors (NETs) comprise a set of neoplasms that originate from cells of the diffuse neuroendocrine system.^1^ Regarding tumor site of origin, small intestine NETs (SI-NETs) represent the second type of NET by annual incidence (1.05 per 100,000 persons) and the third in terms of 20-year limited duration prevalence, according to data from the National Cancer Institute’s Surveillance, Epidemiology, and End Results (SEER) Program.^2^ Unfortunately, despite being regarded as slow-growing and indolent tumors, they exhibit a paradoxically high rate of metastasis at diagnosis (76% of the patients) or during follow-up,^3^ accounting for 27% of all NET with metastasis.^4^ This hinders the already limited therapeutic options available, where, as with other NETs, surgery remains the only curative treatment. When surgery is not feasible, clinical therapies are aimed at stabilizing tumor growth and alleviating tumor-related symptoms, such as systemic therapy with somatostatin analogs (SSAs), the mTOR inhibitor everolimus, as well as ablation and locoregional therapy.^5,6^ Moreover, peptide radionuclide receptor therapy (PRRT) reported, in the NETTER-1 study, a significant improvement in progression-free survival (PFS) and the response rate in comparison with the control group,^7^ although this improvement did not result significant in terms of overall survival (OS).^8^ More recently, the phase III CABINET trial demonstrated that cabozantinib significantly prolonged PFS in patients with previously treated advanced extrapancreatic and pancreatic neuroendocrine tumors, further expanding the therapeutic landscape for these patients.^9^ In view of this evidence, it is essential to identify novel biomarkers and targets to improve current therapies and develop new strategies to tackle these tumors.

One of the key challenges in NETs, particularly SI-NETs, lies in their marked heterogeneity.^10–12^ Although LOH of chromosome 18 is the only recurrent genetic alteration found in over 50% of SI-NETs,^13^ these tumors generally exhibit a low mutational burden,^14^ with *CDKN1B* being the most frequently mutated gene,^15^ albeit in only 8% of cases.^16^ This suggests that additional, less-explored molecular mechanisms, should contribute to their pathobiology.^17,18^ Multi-omics studies,^19^ especially those focusing on epigenetics, have revealed a distinctive DNA methylation landscape in SI-NETs,^20^ characterized by global hypomethylation and associated with tumor aggressiveness,^21^ finally leading to the classification of SI-NETs into three or four molecular subgroups with potential therapeutic implications.^22–25^ In contrast, transcriptomic data remain scarce, although downregulation of genes such as *RASSF1A* and *CDKN2A* has been linked to poor prognosis.^26^ In this scenario, there is now indisputable evidence for the pivotal role of RNA biology in cancer,^27^ which is also rising in NETs,^28^ and particularly involves disruptions in alternative splicing, as tumors frequently exhibit thousands of unique splicing events absent in normal tissue.^29^ This splicing dysregulation, often driven by aberrant splice variants or splicing factor mutations, has emerged as a new hallmark of cancer and a promising source of therapeutic tools.^30–32^ Nevertheless, in contrast to previous work by our group and others demonstrating the impact of splicing alterations in various cancers,^33–36^ including NETs,^37–40^ the specific role of alternative splicing in SI-NETs remains largely unexplored. Taken together, the high metastatic rate of SI-NETs, the limited availability of effective systemic therapies, and the lack of validated biomarkers underscore the need to explore additional molecular mechanisms. All this encouraged us in this study to assess the spliceosomic landscape in SI-NETs to shed light on this issue and thereby foster the discovery of new putative biomarkers to improve patient stratification and outcomes.

## Material and Methods

### Sample collection

The present study was developed using 43 formalin-fixed paraffin-embedded (FFPE) SI-NETs samples, which were collected by surgical resection. All specimens correspond to G1 or G2 tumors acquired from patients with metastases, whose clinicopathological data of the patients are listed in **Table S1**. The samples were obtained from different Spanish hospitals: Central de Asturias University Hospital (Oviedo), Vall d’Hebron University Hospital (Barcelona), Marqués de Valdecilla University Hospital (Santander) and 12 de Octubre University Hospital (Madrid). This work was approved by the Ethics Committees of those hospitals, in accordance with the Declaration of Helsinki and after the signature of the pertinent informed consents by patients. For this study, the samples were divided into two groups according to the survival time of the patients, establishing a longsurvival group (≥ 5 years, *n* = 26) and a short survival group (< 5 years, *n* = 17).

Additionally, expression data from Human Flp-In 293 cells transduced with a shRNA targeting *CPSF2* or with a non-silencing control shRNA were also employed. Samples were prepared using a TruSeq RNA Sample Preparation Kit (Illumina) according to manufacturer’s instructions. These data were publicly available from the Gene Expression Omnibus database under accession GSE60392.^41^

### RNA extraction

Total RNA was isolated from FFPE tissue sections. Manual macrodissection of the tumor area was performed using a scalpel. RNA was extracted using the Maxwell MDx 16 Instrument (Promega, Madrid, Spain) and the Maxwell 16 LEV RNA FFPE Kit (Promega, Madison, WI, USA), according to the manufacturer’s instructions. Briefly, sample slides were deparaffinized with xylene and lysed with proteinase K (ThermoFisher-Scientific, Madrid, Spain) while undergoing heat treatment. Subsequently, the supernatant was treated with DNase and total RNA was caught into a column and further eluted using RNase-free water. Finally, the RNA concentration and purity were assessed using a NanoDrop One Microvolume UV-Vis Spectrophotometer (Thermo Fisher Scientific), while RNA integrity was evaluated using the RNA Integrity Number (RIN) on a 2100 Bioanalyzer (Agilent Technologies, Santa Clara, CA, USA).

### RNA sequencing

Libraries were sequenced on a HiSeq2500 platform (Illumina) with a read length of 2 × 100 bp. On average, each sample yielded 66 million paired-end reads. Image analysis, base calling, and quality scoring were performed using the manufacturer’s Real-Time Analysis software (RTA v1.18.64), and FASTQ files were generated using CASAVA (Illumina’s Consensus Assessment of Sequence and Variation software).

### RNA-seq processing and alignment

Raw reads were processed using *Trim Galore* v0.6.10 and underwent quality analysis to ensure good read quality using *FastQC* v0.12.1. Subsequently, the alignment of these reads to the reference genome (GRCh38) was carried out using *STAR* aligner v2.7.9a^42^ due to its high performance as an aligner and its suitability for subsequent splicing analysis. Specific alignment parameters were defined for further splicing analysis.

### Gene expression quantification

The BAM files generated through alignment were used with the Python tool *htseq*-*count* v2.0.2^43^ to quantify gene expression. For this purpose, the overlap of reads with the annotated gene reference of the GRCh38 genome was assessed. Then, the *DESeq2* v1.38.3 tool was used to import the data and, after removal of the batch effect using the *ComBat-seq* v6.1^44^ tool, they were normalized using variance stabilizing transformation (VST) method, following the procedure provided by the developers.

### Differential expression analysis

Differential expression analysis was used to ascertain which genes were differentially expressed between both groups of samples. To this end, we employed *DESeq2*, as previously described.^36,38^ Genes that fulfilled two criteria were considered differentially expressed: fold-change value > 2 and a *p*-value < 0.05, without applying adjustment due to cohort size and sample similarity.

### RNA splicing analysis

The BAM files generated in the previous steps were analyzed using the *rMATS*-*turbo*^45^ v4.1.2 tool to establish the alternative splicing events that occur differentially between the two groups. The *-novelSS* flag was used to define novel splice sites and we set a significance threshold for those events with a *p*-value < 0.05 and a dPSI (differential Percent Spliced In) < -0.1 or > 0.1. The *rMATS*-derived output, consisting of annotations of splicing events and their quantitative changes across groups, was subjected to further analysis to characterize this process, as detailed below.

### Unsupervised clustering

We selected the VST counts corresponding to the expression of the 208 differentially expressed genes found. Unsupervised hierarchical clustering was applied to these data through the R package *pheatmap*, precisely by Ward clustering method and Euclidean clustering distance. Likewise, we carried out the clustering of the differential splicing events detected and regarded as significant, using the PSI for each event of each sample.

### Enrichment analysis

We employed the lists of differentially expressed genes as well as differentially spliced genes in gene enrichment analyses by obtaining terms from the database of Gene Ontology: Biological Process, that was acquired from Molecular Signatures Databases (*MSigDB*).^46^ The same approach was adopted for the genes outlined in the analysis of genes whose expression or splicing seems to change depending on the *CPSF2* gene expression. To achieve this, we chose the R package *clusterProfiler* v4.6.2.^47^

### Splice events characterization

The files generated by *rMATS* were subjected to several analyses by means of the various tools: *SpliceTools*^48^ to evaluate different splicing event characteristics such as the splice site score of splicing events, the size of the exons or introns affected by differential splicing and the fate of each isoforms regarding nonsense-mediated decay (NMD); *ExonOntology*^49^ to infer certain protein characteristics from the exon genomic coordinates, such as exon function or possible protein post-translational modification (PTM); and *rMAPS*^50^ that is specialized in detecting binding sites of RNA-binding proteins around the splicing events. For the latter, we chose the motifs of our proteins of interest available in the database in addition to the CPSF2 binding motif (AGGUAG) described by Misra *et al*.^41^ Furthermore, we utilized the *rmats2sashimiplot* v3.0.0 tool^45^ to represent sashimi plots of differential splicing events deemed to be relevant.

### Search for RNA-binding motifs

The scan for the presence of selected RNA motifs was performed using *MEME Suite* v5.5.7 software.^51^ Specifically, the *FIMO* tool^52^ was employed to calculate the occurrences of the CPSF2-associated RNA motif within the matched sequences between both cohorts susceptible to be spliced by CPSF2. The cutoff for the *p*-value was set below 10^-4^, following the recommendations of the tool developers.

### Statistical analysis

All statistical analyses were performed using *GraphPad Prism* v8.0.1 (GraphPad, La Jolla, CA, USA) or *R* v4.2.2. For all comparisons, potential outliers were identified and excluded using the ROUT method (Q = 1%), and data distribution was assessed with the Shapiro-Wilk test and, according to the results, Student’s t test or Mann-Whitney U tests were performed. Similarly, correlation analyses were performed with Pearson or Spearman tests according to the normality distribution of the samples. Data are shown as mean ± SD or median ± IQR, as appropriate. Likewise, PLS-DA (Partial Least Squares-Discriminant Analysis) and VIP (Variable Importance in Prediction) Score analyses were conducted to determine the most discriminant molecules within the clinical groups defined. These analyses were performed using *Metaboanalyst* 6.0 (McGill University, Montreal, QC, Canada).^53^

Survival analyses were carried out by running the *survival* v3.4-0 and *survminer* v0.4.9 packages of R. Heatmaps were created with the *pheatmap* 1.0.12 package. All other plots were created using GraphPad Prism 8 or the R package *ggplot2* v3.5.1.

Due to limitations in sample size, which constrain the statistical power of the comparisons, we considered all results with a *p*-value below 0.05 to be significant, except when otherwise specified.

## Results

### Differences in their global transcriptomic profile do not inform sufficiently to understand the dissimilar survival of the two groups of SI-NET patients

Overall analysis of transcriptomic data from SI-NET samples derived from patients categorized as long-versus short-survival (threshold set at 5 years) revealed 208 differentially expressed between both survival groups (**Fig. 1A**, **Table S2**). Of these, 85 were downregulated in the long-survival group and 123 were upregulated. Unsupervised hierarchical clustering based on this gene set showed limited separation between the two survival groups, with samples from both groups intermingled within the clusters (**Fig. 1B**). On the other hand, enrichment analyses of the differentially expressed genes showed associations with distinct biological processes. Thus, genes overexpressed in patients with long survival were linked to immune response processes, such as complement activation, B-cell activation, and phagocytosis (**Fig. 1C**), whereas genes with lower expression in these tumors were associated with processes such as synapses, ion homeostasis or cell division (**Fig. 1D**). Nevertheless, although some potentially informative changes in gene expression could be observed between tumors from patients with short and long survival, no prominent findings or insightful evidence emerged pointing toward the possible causes of differential survival between these two groups of patients.

**Figure 1.**
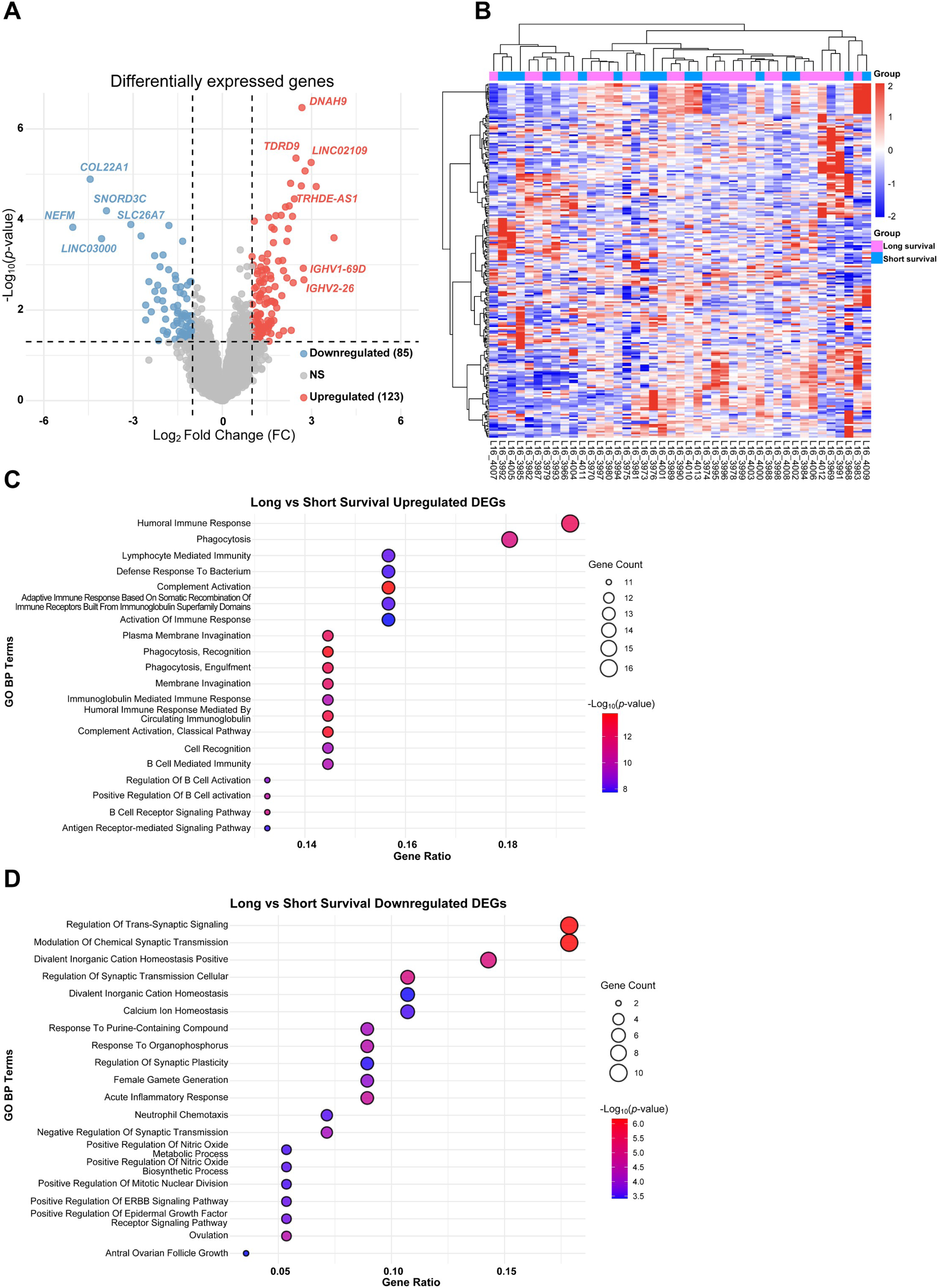
The transcriptomic landscape of the SI-NETs does not clearly discrimiate long and short survival groups. **(A)** Volcano plot of differentially expressed genes when comparing the long vs. short survival group. Upregulated genes are colored in red and downregulated genes are colored in blue. Moreover, genes of each set with the highest fold change and the lowest *p*-value are highlighted with their names. The threshold was set at 0.05 for *p*-value and |2| for fold change. **(B)** Unsupervised hierarchical heatmap of the 208 differentially expressed genes between the two survival groups. Long-survival patient samples are shown in pink and short-survival patient samples are shown in blue. **(C, D)** Enrichment analysis using the Gene Ontology Biological Process database for upregulated **(C)** and downregulated **(D)** genes in the subsets of samples. The size of the dots represents the number of genes matching each term and the color of the dots represents the *p*-value.

### Spliceosomics may help distinguish between short- and long-survival patients

The lack of major illustrative findings derived from global transcriptomic analyses led us to seek for alternative strategies to identify molecular differences between the distinct survival groups. In this context, the growing evidence supporting a major role of alternative splicing dysregulation in cancer prompted us to characterize the splicing landscape in this SI-NETs cohort. A dedicated alternative splicing analysis identified 478 differential splicing events between the short- and long-survival groups, with unsupervised clustering showing clear separation of most samples according to survival group (**Fig. 2A**, **Table S3**). Interestingly, the three splicing events that contributed most to this separation according to the VIP score (*ULK3*, *GUK1* and *ANKRD54*; **Fig. 2B**) corresponded to genes involved in key signaling pathways, such as mTOR or Sonic hedgehog. Specifically, when alternative splicing events (ASEs) were classified according to their class frequency, the most abundant were skipped exon (SE, 154), followed by mutually exclusive exons (MXE, 139) and retained intron (RI, 111), in contrast with the relatively low number of alternative 5’ splice site (A5SS, 40) and alternative 3’ splice site (A3SS, 34) events identified (**Fig. 2C, Fig. S1A**).

**Figure 2.**
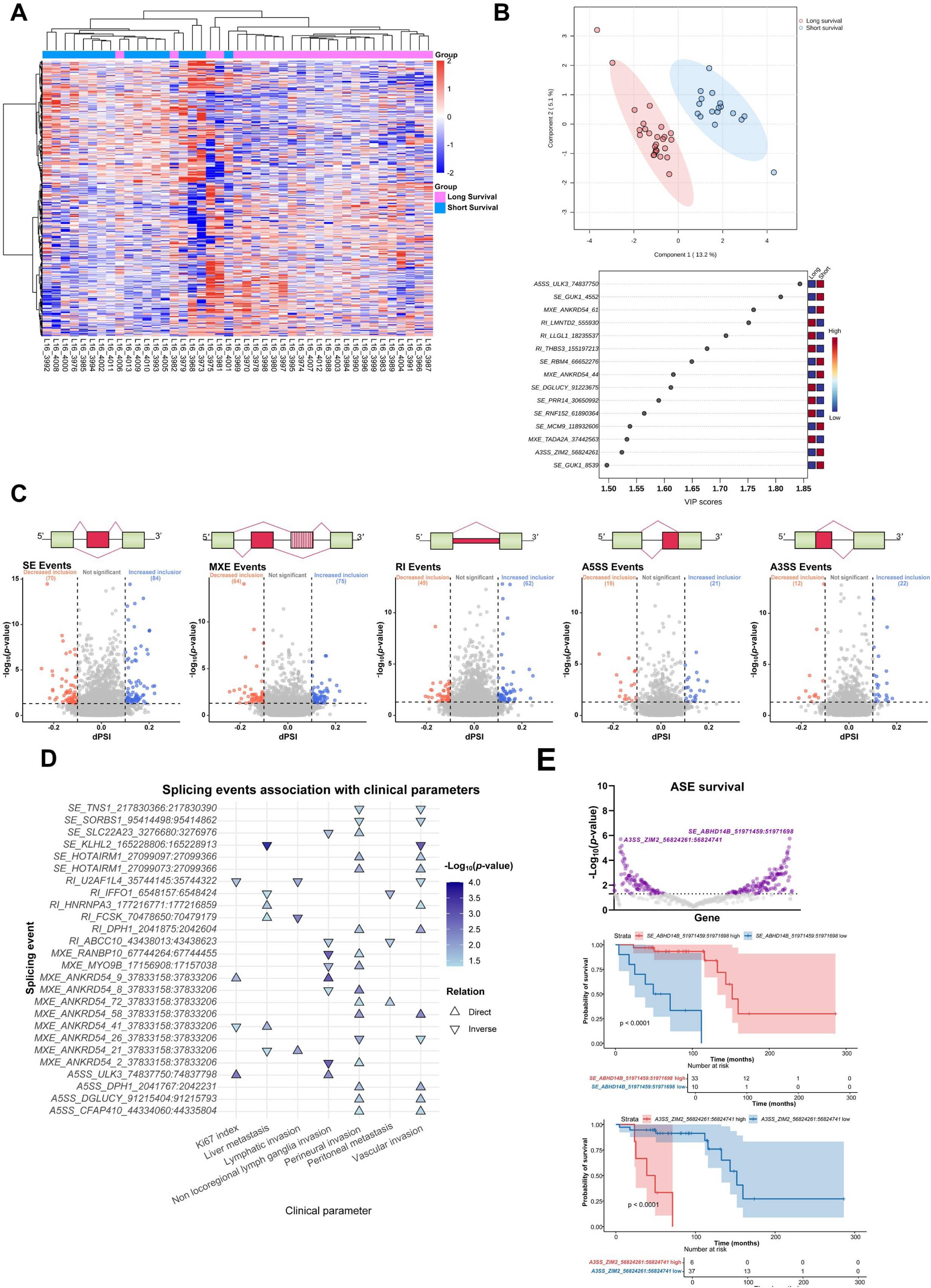
The spliceosomic profile showed strong separation between short- and long-survival groups. **(A)** Unsupervised hierarchical heatmap of the 478 differential spicing events when comparing the long-survival vs. short-survival group. Long-survival patient samples are shown in pink and short-survival patient samples are shown in blue. **(B)** PLS-DA analysis and corresponding VIP scores showing the 15 splicing events that contribute most to the distinction between the two groups. **(C)** Volcano plots for each type of splicing event emphasizing those that were differentially significant between the two sets. Events with minor inclusion appear in red, events with major inclusion in blue, and non-significant events in gray. Graphical representations of each one of the events are presented. The threshold was set at 0.05 for *p*-value and |0.1| for dPSI. **(D)** Association of differential splicing events with selected relevant clinical parameters. Events associated with at least two clinical parameters are represented. The triangles represent the association, being direct if the triangle points upwards and inverse if it points downwards. The *p*-value of these relationships is represented by the color of the triangles. **(E)** Survival analysis of the 478 differential splicing events between the two groups. All significant relationships are shown in purple. The two events with the lowest *p*-value (calculated by log-rank test) are emphasized showing their Kaplan-Meier models as an example. Samples with high PSI are presented in red pink and samples with low PSI for each event are shown in blue. The statistical significance cut-off was established according to *p*-value < 0.05.

Of note, further analysis of the changes detected in ASEs revealed their association with relevant clinical parameters such as Ki67 index, organ invasion, or metastasis at diagnosis. Furthermore, in this regard, we identified 26 differentially spliced events that were significantly linked to at least two of the clinical variables evaluated (**Fig. 2D**). As an example, the A5SS of *ULK3* mentioned above was more frequent in samples from patients with short survival and was correlated with a higher Ki67 index and nonregional lymph node metastasis, highlighting a link between this splicing event and higher tumor aggressiveness. In the same line, and equally relevant, we found numerous associations between splicing events and patient OS, with *ABHD14B* and *ZIM2* standing out among more than 220 events, representing the two most significant extremes, which are presented separately (**Fig. 2E**, **Table S4**).

### Survival differences in SI-NETs patients are characterized by a global spliceosomic perturbation

In the present study, we focused specifically on SE and RI events, as these are among the most frequently studied alternative splicing events in cancer, and are well supported by currently available analytical tools are designed to analyze these types of splicing events. Further, we aimed at understanding the potential consequences of the emerging ASEs, particularly in relation to their predicted degradation by the NMD system. Using this approach, we first identified a predicted shift in the fate of 67 transcripts, driven by changes in NMD-targeting motifs located within alternatively excluded exons. In samples from patients with long-survival, 31 transcripts were predicted to lose NMD susceptibility and 3 to acquire it, whereas in samples from patients with short-survival, 31 transcripts were predicted to lose NMD susceptibility and 2 to acquire it (**Fig. 3A**). In line with this, we next investigated the potential functional impact of the differentially spliced exons on the resulting proteins. Most of these exons have a structural function (25%), followed by those involved in PTMs (23.53%), binding (9.93%), localization (8.82%) and, to a lesser extent, catalytic activity (4.04%). Of the analyzed exons, 16.18% were not present in the reference database, and 12.50% could not be assigned to any of the aforementioned functional categories (**Fig. 3B**). Considering the outcome of the event, i.e. whether the exon was retained or skipped, it was observed that, for exons related to structure or PTMs (the most represented categories), approximately 70% were retained in the final transcripts of samples from long-survival patients, compared to those with short-survival (**Fig. S1B**). Within the PTM-related exons, 173 PTMs were identified, the majority of which were phosphorylations (38.15%), followed by, in a much smaller proportion, methylations (5.20%), acetylations (4.62%), and ubiquitinylations (4.62%) (**Fig. S1C**). Consistent with the findings on exon function, most exons that encode phosphorylation sites in the final protein were more preserved in the long-survival group (**Fig. S1D**). The SE event in *HNRNPA1* or the RI event in *PTPN6* serve to exemplify this differential splicing between the two groups (**Fig. S1E**). These results highlight the importance of splicing in regulating RNA expression and protein function, an aspect that has not been previously explored in SI-NETs.

**Figure 3.**
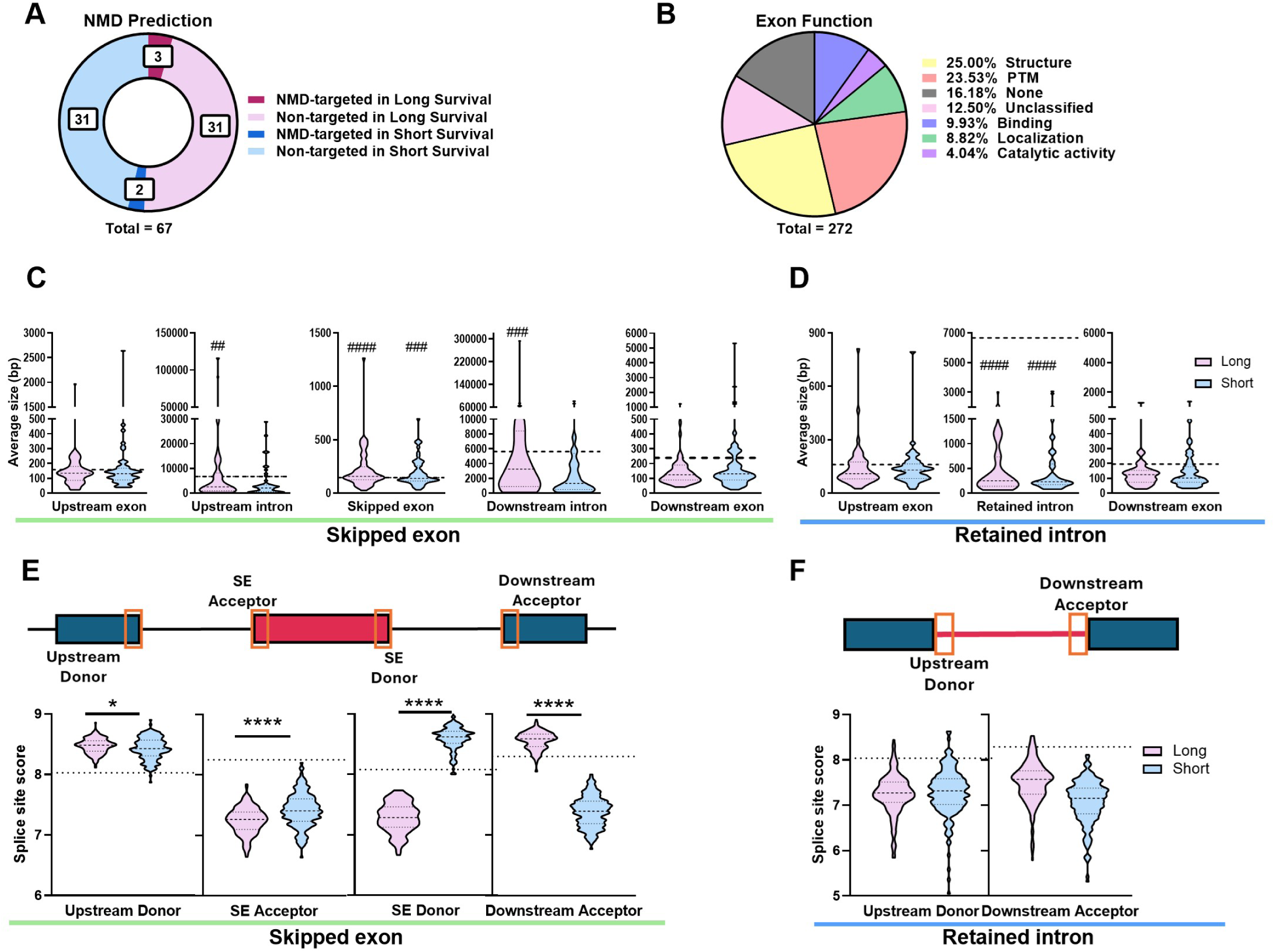
Differential splicing events show predicted functional consequences and altered splice-site features. **(A)** Prediction of transcripts susceptible to NMD following a fate switch between the two groups. Transcripts with a fate change in the long survival samples are shown in pink and those detected in the short survival samples are shown in blue. Dark colors represent transcripts that became NMD-targeted and previously were not, while light colors represent transcripts that avoided NMD. **(B)** Functional characterization of exons that were differentially affected by exon skipping events between both groups of samples. **(C)** Average size of the crucial regions in the SE event. **(D)** Average size of the crucial regions in the RI event. The long survival group is represented in pink and the short survival group in blue. Data are represented by median ± interquartile range. The dashed line represents the average size of all annotated events. The symbols indicate significant differences when comparing each group with the mean of the annotated events, such that ## < 0.01, ### < 0.001 and #### < 0.0001. **(E)** Splice site score of each of the critical positions in the SE event. **(F)** Splice site score of each of the critical positions in the RI event. The long-survival group is shown in pink and the short-survival group in blue. Data are represented by median ± interquartile range. The dashed line represents the average score of all annotated events. This includes a representation of the specific location of each of these regions in the events analyzed. Symbols depict significant differences when comparing the groups to each other, such that * *p* < 0.05 and **** *p* < 0.0001.

These differential splicing events may reflect alterations in the processing of precursor RNA molecules. For this reason, we studied the characteristics of the key regions of the transcript that participate in different splicing events. Interestingly, we found that, in the case of SE events, both flanking introns in long-survival group were significantly shorter than the average of the annotated ones, whereas they tended to be longer than those in the short-survival one (**Fig. 3C**). A similar pattern was observed for RI events, in which the analyzed introns were smaller than the average size of annotated RI events in both survival groups and showed comparable sizes groups between groups (**Fig. 3D**). In line with this, the splice site score analysis of SE events revealed substantial differences between our samples and the general annotation. More importantly, long- and short-survival samples exhibited marked differences, with long-survival samples showing stronger upstream donor and downstream acceptor sites, but weaker SE acceptor and donor sites (**Fig. 3E**). In contrast, no significant changes were detected in splice site score for RI events. (**Fig. 3F**). These results indicate that splicing in SI-NETs may be altered due to the differential splice site selection, resulting in splicing elements with lengths that deviate from the average, which, in turn, could generate changes in splicing potentially associated with patient survival.

### Alterations in the splicing machinery as a link between splicing changes and survival in SI-NETs patients

Based on the findings regarding splicing changes associated with patientś survival, we hypothesized that these alterations could result from an upstream disturbance in the splicing machinery, the ultimate effector of the splicing process. To test this notion, we first analyzed the expression of the 315 genes related to the splicing process, which revealed alterations in 47 of them, a 15% of the total (**Fig. 4A**). Moreover, the analysis of their possible correlation to OS showed that 23 of the 47 dysregulated factors (48.9%) were significantly associated with patient survival (**Fig. 4B**, **S2-4)**. Additionally, four of these splicing factors were associated with selected clinical parameters: *PSPC1* with Ki-67 index, *DHX32* and *SRSF12* with metastasis, and *CPSF2* with perineural and vascular invasion (**Fig. 4C**).

**Figure 4.**
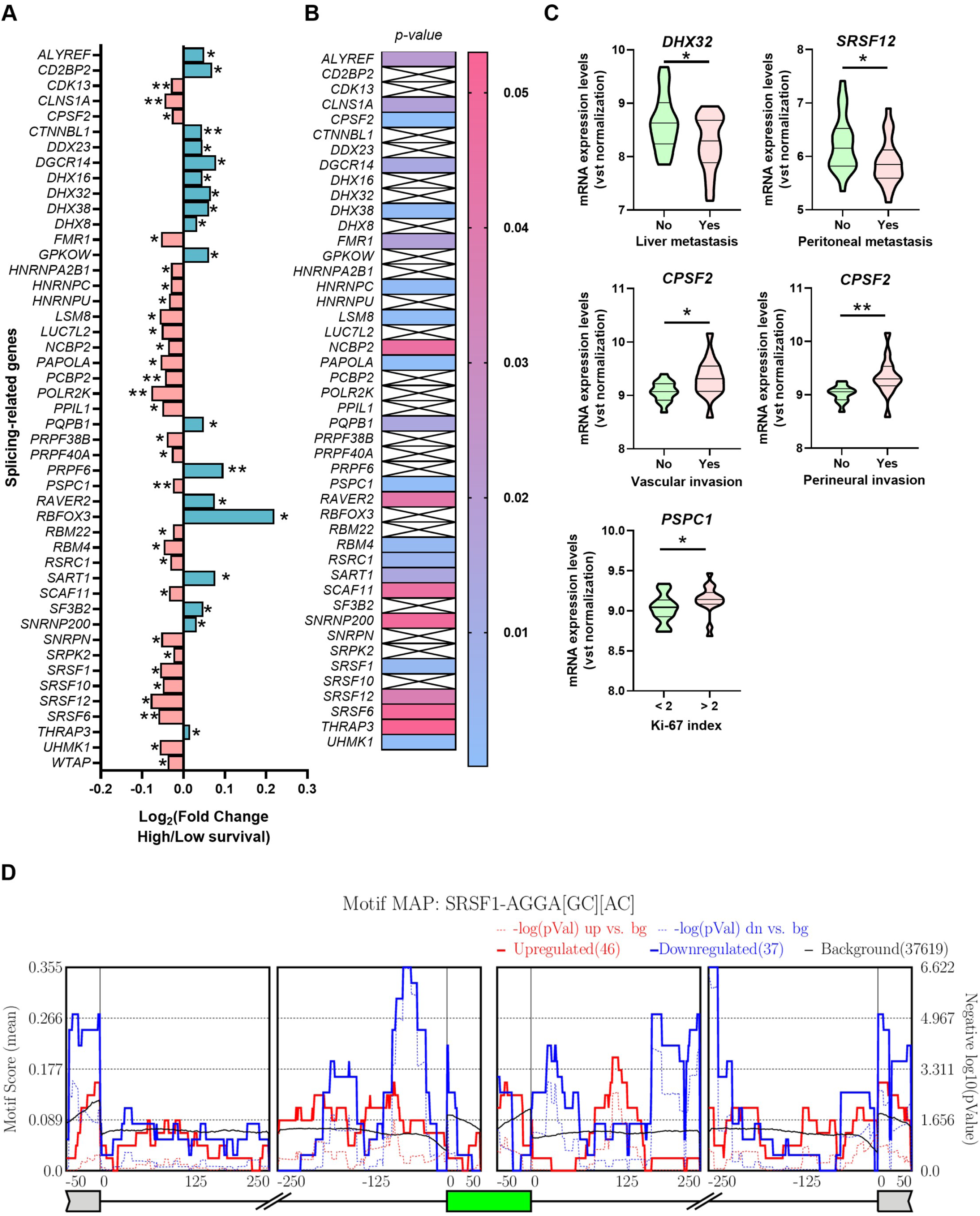
Different expression profile of splicing machinery components in SI-NETs samples from patients with long and short survival. **(A)** Fold change of mRNA levels expressions of spliceosome components and splicing factors that are altered when comparing SI-NETs from patients with long survival with SI-NETs from patients with short survival. Data are represented by fold change mRNA levels mean ± SD. **(B)** Survival analysis of the splicing machinery components that are different between both sets. Genes with significant association are colored according to their *p*-value. *P*-value threshold (calculated by log-rank test) set at *p* < 0.05. **(C)** Association of selected machinery genes with relevant clinical parameters. Data are represented by median ± interquartile range. Symbols depict significant differences when comparing the groups to each other, such that * *p* < 0.05 and ** *p* < 0.01. **(D)** Mean of the motif score for the SRSF1 splicing factor in the detected SE events. The red line represents the exons that are more included in the long survival group, the blue line represents the exons that are less included, and the black line represents the mean of the SE events scored. The solid lines symbolize the mean motif score, while the dashed lines represent the *p*-value. *P*-value threshold set at *p* < 0.05.

Therefore, once we established a subset of factors of interest (altered between groups and associated with clinical parameters), we sought to ascertain whether they could predictably drive the differential splicing events detected in our samples. For this purpose, we analyzed the existence of their binding/recognition motifs (motif score) in our data using *rMAPS* and obtained significant variations among both experimental groups, as illustrated for SRSF1. This factor is more expressed in short survival group, where high density peaks of the SRSF1 motif are found upstream and downstream of the exons included in these samples. (**Fig. 4D**).

### *CPSF2* might be essential in the aggressiveness and worse prognosis of SI-NETs

Among the numerous altered splicing-related genes, the most prominent was *CPSF2*, which showed higher expression in individuals with worse prognosis and was associated with reduced survival, as well as with perineural and vascular invasion. For this reason, we conducted a more exhaustive analysis. When evaluating the potential influence of this factor on the differential splicing events identified, we discovered a significant enrichment of the CPSF2 binding motif (AGGUAG) within the target exons involved in the SE events, as well as in the downstream introns of this gene element (**Fig. 5A**).

**Figure 5.**
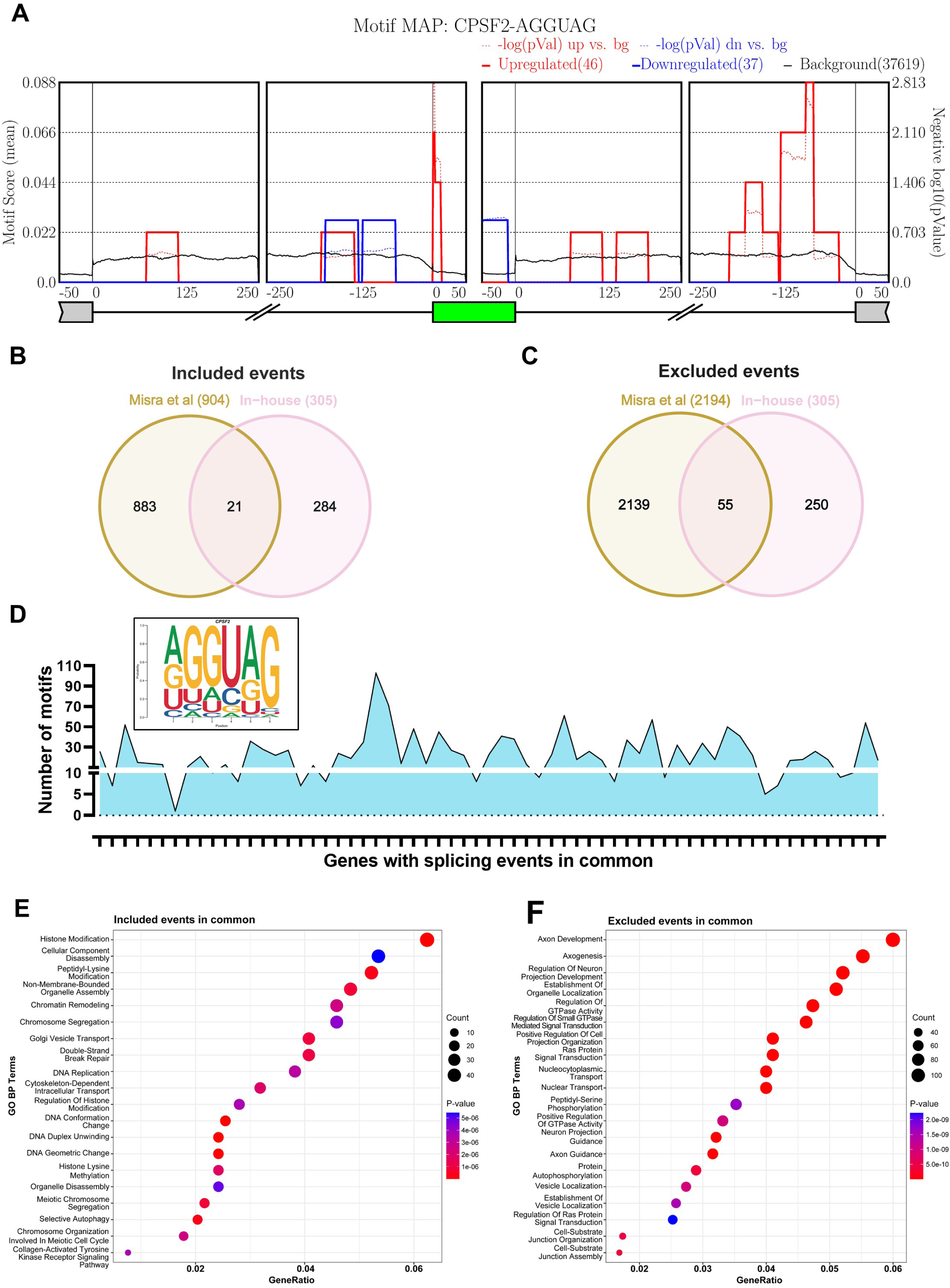
CPSF2 emerges as a candidate regulator potentially associated with poor prognosis in SI-NET. **(A)** Mean of the motif score for the CPSF2 factor in the detected SE events. The red line represents the exons that are more included in the long survival group, the blue line represents the exons that are less included, and the black line represents the mean of the SE events scored. The solid lines symbolize the mean motif score, while the dashed lines represent the *p*-value. *P*-value threshold set at *p* < 0.05. **(B)** Venn diagram of overlapping genes affected by splicing events between the in-house cohort and the inclusion events of the Misra *et al.* cohort. **(C)** Venn diagram of overlapping genes affected by splicing events between the in-house cohort and the exclusion events of the Misra *et al.* cohort. **(D)** Identification of occurrences of the specific CPSF2 RNA-binding motif (shown at the top) in the RNA sequences of the overlapping genes between both cohorts to be susceptible to undergoing splicing by CPSF2. **(E, F)** Enrichment analysis using the Gene Ontology Biological Process database for included **(E)** and excluded **(F)** events. The size of the dots represents the number of genes matching each term and the color of the dots represents the *p*-value.

To further investigate the impact of *CPSF2* in SI-NETs, we obtained available sequencing data following *CPSF2* silencing in Flp-In-293 cells, along with the list of genes whose splicing is mediated by *CPSF2*, according to the iCLIP-seq results reported by Misra *et al*.^41^ Upon examining the number of overlapping genes that were positively correlated with *CPSF2* expression in our cohort and those whose expression was reduced by silencing this gene in the Misra *et al.* dataset, we identified 21 overlapping genes. This number was very close to the 23 overlapping genes found when comparing those that increased their expression upon silencing *CPSF2* and that were negatively correlated in our cohort (**Fig. S5A-D**). Moreover, when characterizing the genes potentially influenced by *CPSF2* expression, we observed distinct enrichment patterns in both cases. Positively correlated genes were associated with hormone regulation and transport, as well as with cell signaling (**Fig. S5E**), whereas negatively correlated genes were linked to cell motility and response to hypoxia (**Fig. S5F**).

Concerning splicing, comparison of the genes affected by events detected in our cohort with those reported by Misra *et al.* identified 21 overlapping genes in inclusion events and 55 in exclusion events (**Fig. 5B-C**, **Table S5**). To further support this analysis, we searched for the CPSF2 binding motif and detected at least one ocurrence of this motif in the mRNA sequence of each overlapping target genes (**Fig. 5D**). Finally, enrichment analysis of both gene groups demonstrated once more two clearly distinct functional profiles: inclusion events were associated with DNA regulation and modification, as well as cellular transport, whereas exclusion events were associated with nerve development, cell signaling, and cellular organization (**Fig. 5E-F**).

## Discussion

SI-NETs are indolent, slow-growing tumors with a high rate of metastasis.^3^ In the largest single-center prognostic study of patients with SI-NETs to date, about two-thirds of patients presented metastases at the time of diagnosis.^54^ Unfortunately, radical surgery appears to be the only potentially curative treatment for both local and metastatic disease.^5^ At the systemic level, SSAs are mostly used in first-line therapy, based on the two main clinical trials that demonstrated benefits in terms of time to tumor progression, which, however, did not observe a positive impact on OS.^55,56^ Thus, although PRRT has recently been recommended as a second-line treatment, no curative therapies remain available beyond surgery.^6^ For all these reasons, a better molecular characterization of the disease is needed to more accurately classify patients and offer them more effective, personalized treatment. In this study, we offer primary evidence that the splicing process is altered in relation to patient survival in SI-NETs, providing a novel angle for approaching this disease.

Global transcriptomic analysis of a cohort of SI-NET patients revealed that, despite changes in expression of numerous genes between samples from patients with short-(< 5 years) or long-survival (≥ 5 years), this parameter was unable to accurately cluster the samples according to survival. This may be due to the high heterogeneity of NETs in general, or to the specific characteristics of this cohort, in which the tumors presented very particular clinical features. However, in keeping with our observations, previous reports also indicated that comparisons between primary SI-NETs of differing aggressiveness did not reveal significantly discriminant RNA expression changes.^57^ Since survival differences between patients did not seem to be reflected (and perhaps lie in) overt transcriptomic dissimilarities, we posited that the change could originate at a point subsequent to gene expression. In this scenario, RNA splicing has arisen in recent years as a process whose imbalance plays a fundamental role in numerous pathologies,^58^ including cancer^32^ and more specifically, NETs.^28,40^ Indeed, splicing dysregulation has not only been proposed as an additional hallmark of cancer, but has been demonstrated to pervade every and each of the other hallmarks proposed by Hanahan and Weinberg.^30,59,60^ Accordingly, we decided to investigate whether this regulatory mechanism could shed a new molecular light of the survival-related differences in SI-NETs, a previously unexplored avenue in these tumors.

The “spliceosomic” approach unveiled a radically contrasting picture, providing a splicing event profile that almost completely distinguished between both patient groups. This nearly perfect linkage of splicing event patterns to patient prognosis was not totally unexpected, as parallel discoveries have previously been reported in colorectal cancer,^61^ hepatocellular carcinoma^62^ or pancreatic cancer^63^, highlighting the tight bonds between splicing and cancer. Thus, our results on SI-NETs further document the plausible importance of the association between ASEs and clinical parameters, in this case, patient survival, strengthening the cumulative evidence reported in other cancers.^29,64,65^ The distinct, survival-related alterations found in splicing events led us to investigate the status of the upstream machinery (i.e., the spliceosome core components and related splicing factors), which is ultimately responsible for generating transcript variants. Interestingly, profile analysis of the splicing machinery revealed that a significant proportion of its elements undergo substantial changes in expression, which are, moreover, associated with patient survival and relevant parameters of tumor aggressiveness. It could be conceivable that these changes are unrelated to those observed previously in splicing events and could merely be due to random variation. However, our results argue against this notion, in that some of the key dysregulated splicing factors, including *SRSF1* or *CPSF2*, are known to bind near the affected splicing events, and may therefore contribute to the observed changes. Furthermore, we detected differences in splice-site selection and in the length of genomic regions involved in SE and RI events, suggesting alterations in the activity of the splicing machinery that may affect the resulting RNA product. Hence, these findings support a link between altered expression of splicing machinery components and survival-associated splicing changes in SI-NETs.

The underpinnings of the plausible functional significance of splicing changes observed, and on how these are related (or might even contribute) to increased or decreased survival in SI-NET patients remain to be elucidated. This notwithstanding, our present findings already provide suggestive evidence that such significance is real. Thus, in-depth analyses of the altered splicing events reveal a clear impact of the expected fate of the affected transcripts, including fate switches related to NMD, or the loss or gain of exons involved in catalytic activity, binding, or regulation through PTMs of the final protein. In fact, these observations are clearly in line with a growing body of evidence indicating that shifts in gene splicing derive in more aggressive cancer phenotypes, either by preventing the processing of new, aberrant transcripts by the NMD machinery^66,67^ or by generating distinct isoforms with characteristics that enhance tumor aggressiveness and/or malignancy.^68–70^

To further explore the relationship between changes in the splicing machinery and the dysregulation of alternative splicing, we selected *CPSF2,* since this gene, scarcely studied in cancer to date, was overexpressed in short-survival samples and associated with perineural and vascular invasion. Of note, our results are in keeping with the limited, previous reports on CPSF2 in hepatocellular carcinoma, adrenocortical carcinoma, uveal melanoma,^71^ and colorectal cancer, where high mRNA levels of this gene were associated with poor prognosis in terms of disease-specific, disease-free, and recurrence-free survival, although CPSF2 has shown context-dependent prognostic associations across cancer types.^72^ CPSF2 is a core component of the CPSF complex and is involved in 3’-end processing of pre-mRNAs, where it plays a crucial role in linking transcription, splicing, and polyadenylation through interactions with factors such as CPSF73 and mCF/mPSF.^73–76^ Beyond its canonical processing function, CPSF2 has emerged as a key player in alternative splicing regulation, acting both independently and in cooperation with core spliceosome components, such as U1 and U2 or the U2AF2 factor,^77–79^ and splicing factors like RBFOX2, NOVA2, and HNRNPA1.^41^ Notably, only one-third of RBFOX2-regulated exons were independent of CPSF2, while a majority of CPSF2-regulated exons functioned independently of RBFOX2, highlighting its broader influence. It is worth noting the importance of RBFOX2 in tumor progression and metastasis.^63^ In our study, motif enrichment analyses showed CPSF2 binding within or near exons affected by SE, aligning with its known role in modulating splicing outcomes.^41^ Moreover, predicted CPSF2-mediated events were enriched in genes involved in cell migration, cytoskeletal remodeling, and key oncogenic signaling pathways, as well as in processes linked to tumor aggressiveness and treatment resistance, including chromatin remodeling, histone modification, DNA repair, and selective autophagy, that has been linked to resistance to treatment.^80^ Consistent with these findings, *CPSF2* expression has been correlated with markers of genomic instability such as microsatellite instability and tumor mutational burden,^71^ supporting its potential involvement in tumor progression and malignancy.^81,82^

The main limitations of this study are its sample size, the single-cohort design, the absence of an independent validation cohort, and the exploratory nature of high-dimensional analyses performed using nominal *p*-value thresholds. In addition, potential imbalances between the short- and long-survival groups in relevant clinicopathological variables, treatments received, sample type, and follow-up duration could not be robustly assessed. Therefore, survival-defined groups may capture differences related to unmeasured clinical confounders, and residual confounding cannot be excluded. However, the development of suitable SI-NET models remains highly challenging, as both preclinical animal models and bona fide SI-NET cell lines are notably limited for this particular type of neoplasm. ^83^

## Conclusions

Overall, these findings provide primary evidence on the potential association between splicing dysregulation and survival SI-NET patients, a relationship that was not apparent from global transcriptomic analysis alone. Furthermore, we identify a plausible link between survival-related alterations in components of the splicing machinery and the differential splicing events they may regulate. Finally, given the growing therapeutic relevance of RNA splicing, our data identify previously unexplored candidates within the splicing machinery that warrant further investigation as potential biomarkers and as a basis for future splicing-targeted strategies in SI-NETs.

## Ethics declaration

All procedures performed in studies involving human participants were in accordance with the ethical standards of the institutional and/or national research committee and with the 1964 Helsinki Declaration and its later amendments or comparable ethical standards. The study was approved by the Bioethics Committee of the Medical University of Cordoba/IMIBIC (protocol code PI19/01603, v1 – 13/03/2019) and Ethics Committees from all the Hospitals involved in the study. Informed consent from each patient was obtained through the pertinent Biobanks.

## Authors contribution

All authors contributed to the study conceptualization and design. Samples and clinical data were collected by A.G.A., J.H., A.G., C.L.L., P.J.F., R.G.C., S.F., L.B., A.C., A.V., and J.C. Material preparation and data curation were performed by V.G.V., M.T.M.M., A.G.A, R.B.E., S.P.A., A.I.C., and J.P.C. Investigation, methodology, software and formal analysis were performed by V.G.V., M.T.M.M., and R.B.E. The funding was obtained by S.P.A., A.I.C., J.C., and J.P.C. and all of them took charge of project administration and supervision. The first draft of the manuscript was written by V.G.V. and S.P.A. All authors reviewed and approved the final manuscript.

## Acknowledgements

We sincerely thank all the patients and their families for generously donating the samples and clinical data for research purposes. We also acknowledge the technical help of the personnel of core services of the IMIBIC (UCAIB) and University of Cordoba (SCAI).

## Appendix A. Supplementary data

All data supporting the findings of this study are available within the paper and its Supplementary Information.

## Figure legends

**Supplementary Figure S1. (A)** Distribution of differential exclusion and inclusion splicing events between the two conditions according to event type. **(B)** Distribution of differential exclusion and inclusion SE events among both conditions according to exon function. **(C)** Representation of PTMs occurring in exons undergoing differential exon skipping between the two conditions. **(D)** Distribution of differential exclusion and inclusion exon skipping events among both sets according to the type of PTMs that the exon undergoes. **(E)** Sashimi plots of one SE event (left) and one RI event (right) representative of the differences between the two tumor groups. In purple are shown the RPKMs by coordinates in the long-survival group and in red, those in the short-survival group. There is a schematic of the two splicing variants for each of the events depicted.

**Supplementary Figure S2.** Kaplan-Meier significant analyses of the components of the splicing machinery differentially expressed between the long-survival and short-survival samples. The respective curves at high (red) and low (blue) levels of each factor are shown, as well as the *p*-value calculated by log-rank test. Threshold *p*-value set at *p* < 0.05.

**Supplementary Figure S3.** Kaplan-Meier significant analyses of the components of the splicing machinery differentially expressed between the long-survival and short-survival samples. The respective curves at high (red) and low (blue) levels of each factor are shown, as well as the *p*-value calculated by log-rank test. Threshold *p*-value set at *p* < 0.05.

**Supplementary Figure S4.** Kaplan-Meier significant analyses of the components of the splicing machinery differentially expressed between the long-survival and short-survival samples. The respective curves at high (red) and low (blue) levels of each factor are shown, as well as the *p*-value calculated by log-rank test. Threshold *p*-value set at *p* < 0.05.

**Supplementary Figure S5. (A)** Venn diagram of overlapping positively correlated-genes between the in-house cohort and the Misra *et al.* cohort. **(B)** Venn diagram of overlapping negatively correlated-genes between the in-house cohort and the Misra *et al.* cohort. **(C)** Correlations of *CPSF2* with the overlapping positively correlated-genes between the in-house cohort and the Misra *et al.* cohort. **(D)** Correlations of *CPSF2* with the overlapping negatively correlated-genes between the in-house cohort and the Misra *et al.* cohort. The bars display the correlation coefficient between *CPSF2* with each gene, while the color of the bars represents the *p*-value of each correlation. **(E, F)** Enrichment analysis using the Gene Ontology Biological Process database for positively **(E)** and negatively **(F)** correlated-genes overlapping between the in-house cohort and the Misra *et al.* cohort. The size of the dots represents the number of genes matching each term and the color of the dots represents the *p*-value.

## Abbreviations

A3SS: alternative 3’ splice site
A5SS: alternative 5’ splice site
ASEs: alternative splicing events
BAM: Binary Alignment Map
CASAVA: Consensus Assessment of Sequence and Variation
CPSF: cleavage and polyadenylation specificity factor
dPSI: differential Percent Spliced In
FFPE: formalin-fixed paraffin-embedded
iCLIP-seq: individual-nucleotide resolution UV crosslinking and immunoprecipitation followed by sequencing
LOH: loss of heterozygosity
mCF: mammalian cleavage factor
mPSF: mammalian polyadenylation specificity factor
MSigDB: Molecular Signatures Databases
mTOR: mechanistic target of rapamycin
MXE: mutually exclusive exons
NETs: neuroendocrine tumors
NMD: nonsense-mediated decay
OS: overall survival
PFS: progression-free survival
PLS-DA: Partial Least Squares-Discriminant Analysis
PSI: Percent Spliced In
pre-mRNA: precursor messenger RNA
PRRT: Peptide Receptor Radionuclide Therapy
PTM: post-translational modification
RI: retained intron
RIN: RNA Integrity Number
RNA-seq: RNA sequencing
RTA: Real-Time Analysis
SE: skipped exon
SEER: Surveillance, Epidemiology, and End Results
SI-NETs: small intestine neuroendocrine tumors
snRNP: small nuclear ribonucleoproteins
SSAs: somatostatin analogs
VIP: Variable Importance in Projection
VST: variance stabilizing transformation.

