## Supplementary Material for "From Splicing to Survival: Understanding the Molecular Basis of Prognostic Differences in advanced Small Intestine Neuroendocrine Tumors"

Fig. S1

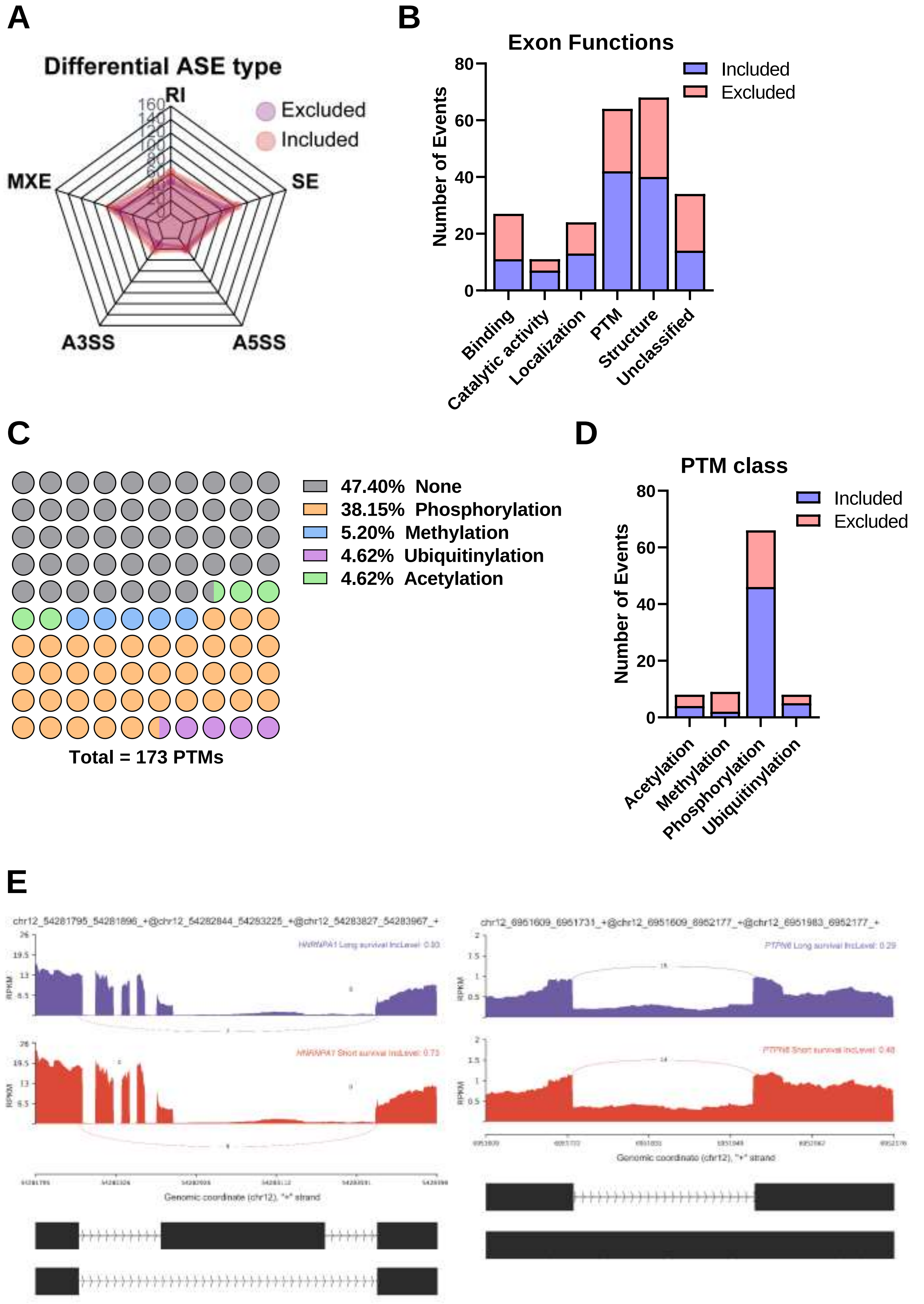
A
Differential ASE type
RI
SE
A5SS
A3SS
MXE
Excluded
Included
B
Exon Functions
Included
Excluded
Number of Events
Binding
Catalytic activity
Localization
PTM
Structure
Unclassified
C
47.40% None
38.15% Phosphorylation
5.20% Methylation
4.62% Ubiquitinylation
4.62% Acetylation
Total = 173 PTMs
D
PTM class
Included
Excluded
Number of Events
Acetylation
Methylation
Phosphorylation
Ubiquitinylation
E
chr12_54281795_54281896_+@chr12_54282844_54283225_+@chr12_54283827_54283967_+
chr12_6951609_6951731_+@chr12_6951609_6952177_+@chr12_6951983_6952177_+
RPKM
Genomic coordinate (chr12), "+" strand

### Fig. S2

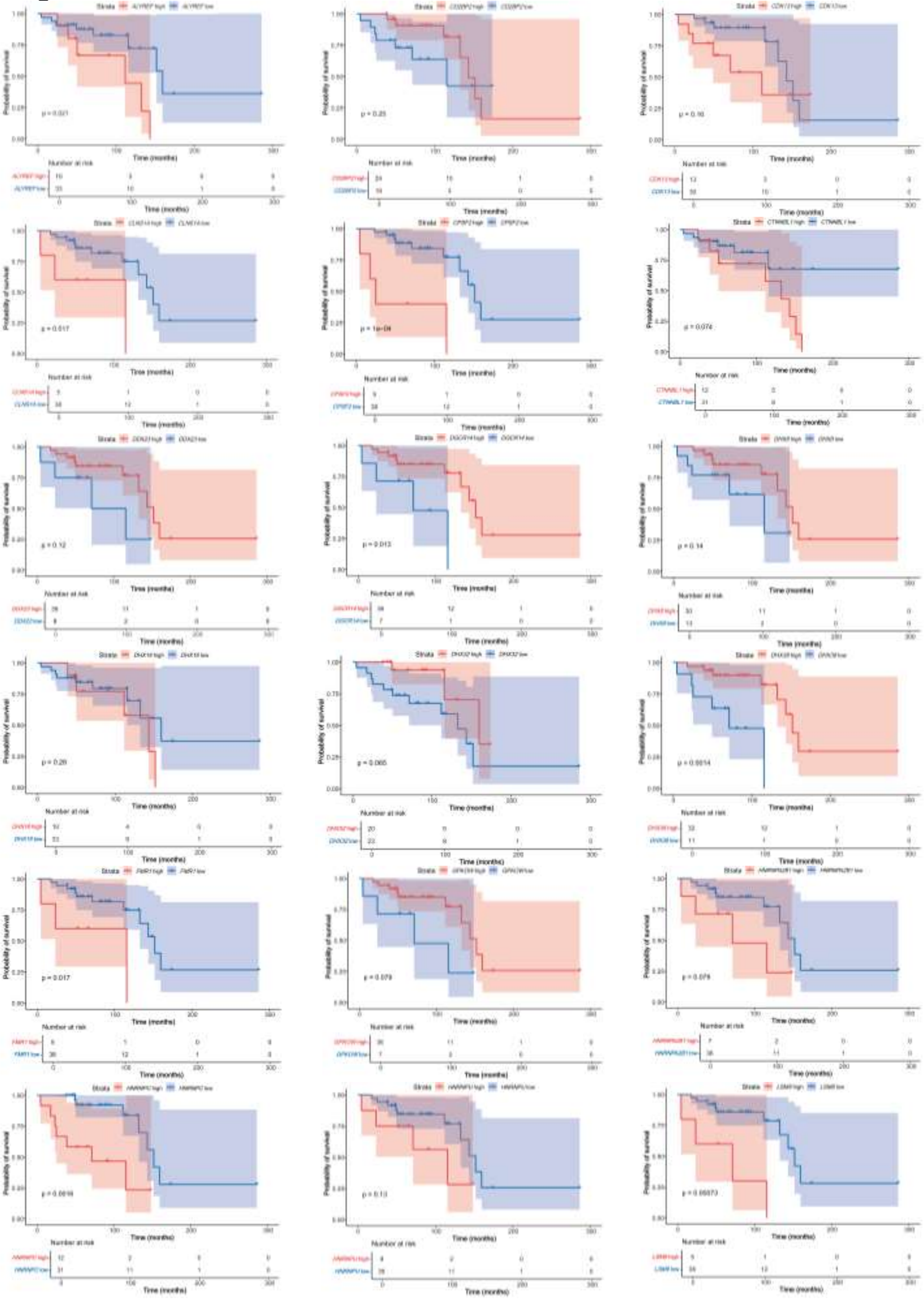

**Fig. S3**

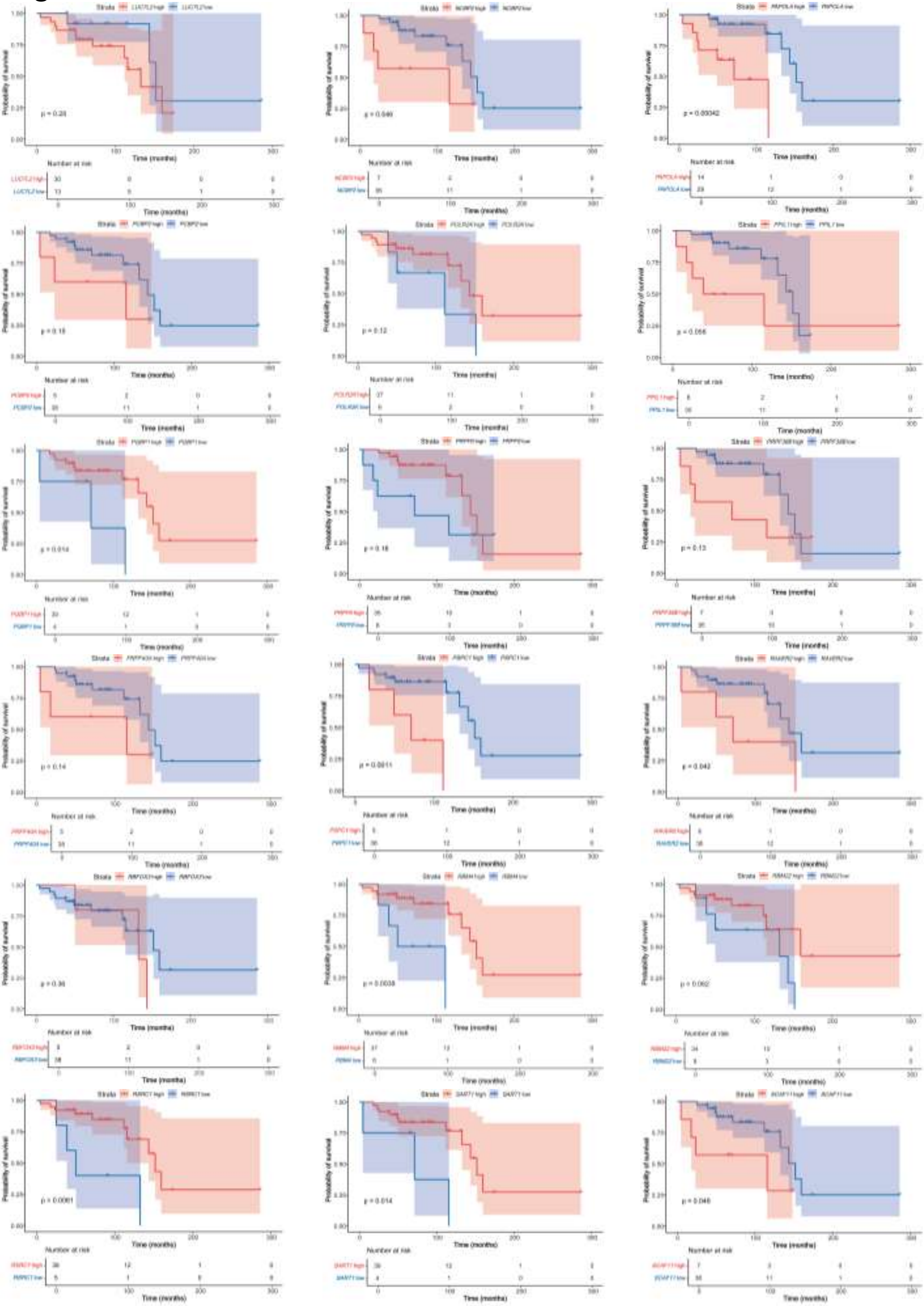

### Fig. S4

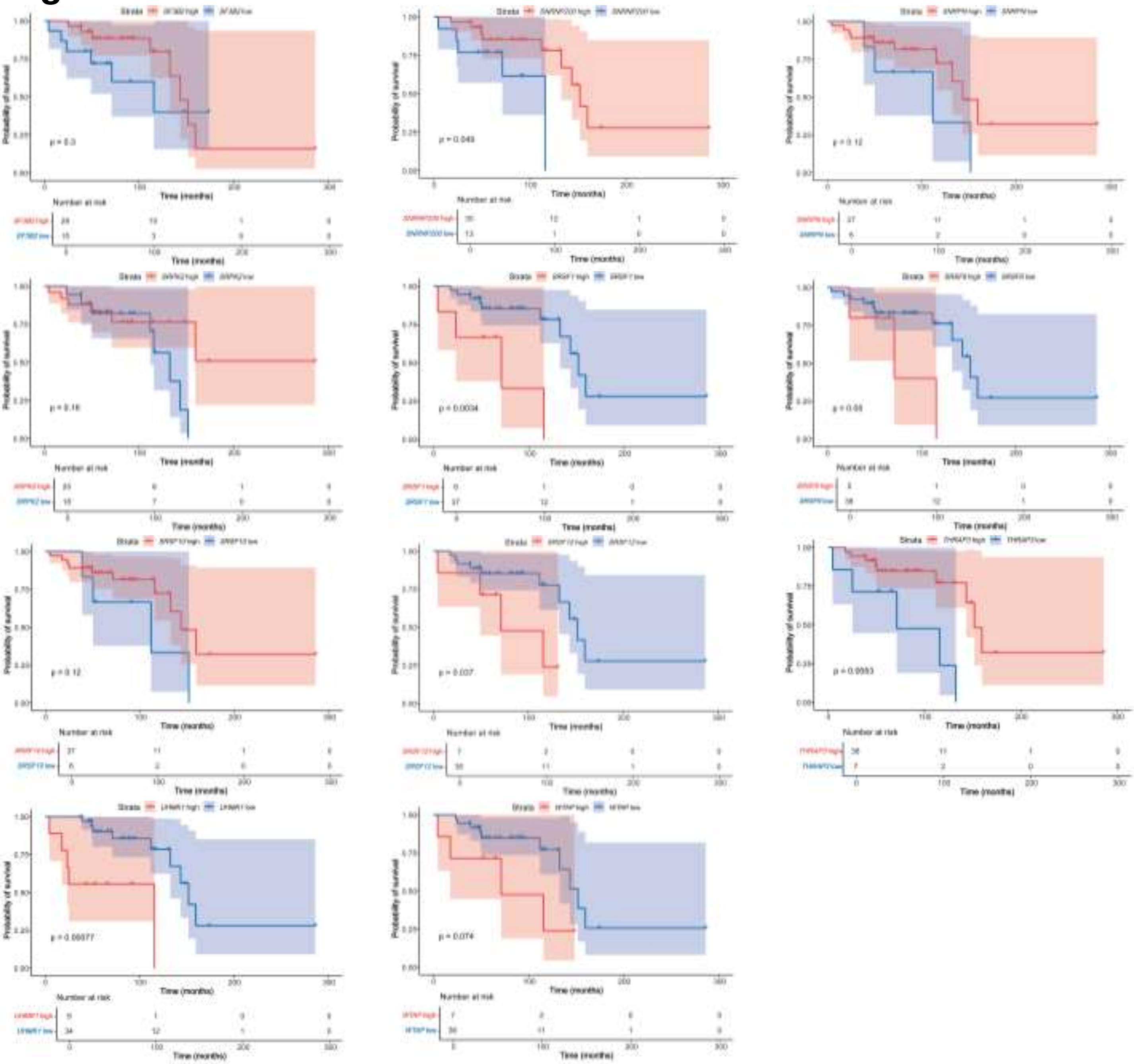

Fig. S5

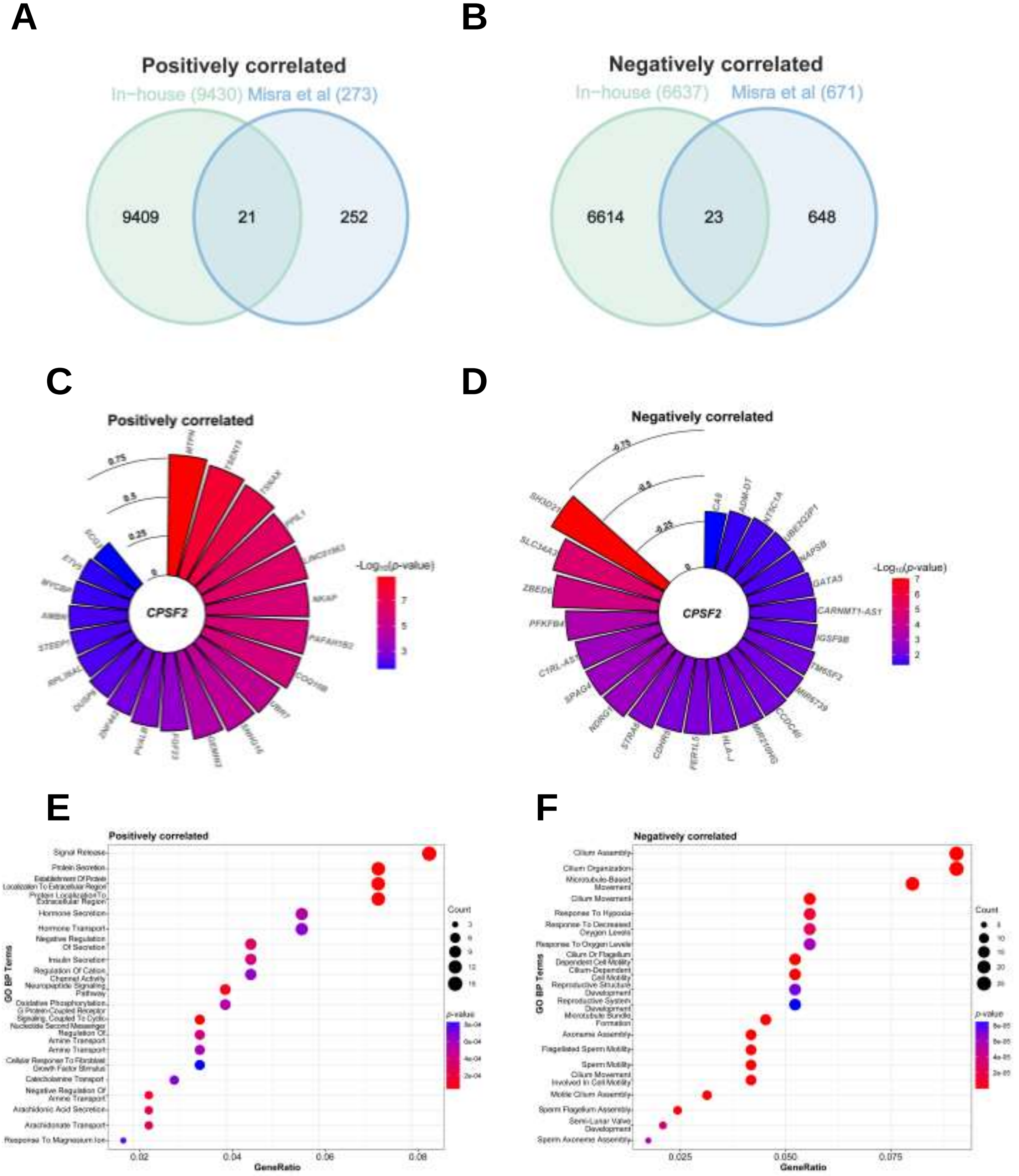
A
Positively correlated
In−house (9430)
Misra et al (273)
9409
21
252
B
Negatively correlated
In−house (6637)
Misra et al (671)
6614
23
648
C
Positively correlated
CPSF2
-Log10(p-value)
D
Negatively correlated
CPSF2
-Log10(p-value)
E
Positively correlated
GO BP Terms
GeneRatio
Count
p.value
F
Negatively correlated
GO BP Terms
GeneRatio
Count
p.value

**Table S1. Summary of the clinical parameters of the SI-NETs patients.**

| Parameter | High survival | Low survival | Total |
|---|---|---|---|
| **Number of samples** | 26 | 17 | 43 |
| **Age (mean ± SD, years)** | 64.5 ± 11.3 | 64.5 ± 12.8 | 64.5 ± 11.8 |
| **Survival (mean ± SD, months)** | 113.7 ± 46.9 | 41.4 ± 15 | 85.1 ± 51.7 |
| **Female (%)** | 30.8 | 52.9 | 39.5 |
| **Male (%)** | 69.2 | 47.1 | 60.5 |
| **Ki-67 (< 2, %)** | 50 | 52.9 | 51.2 |
| **Liver metastasis (%)** | 50 | 64.7 | 55.8 |
| **Lymphatic invasion (%)** | 34.6 | 47.1 | 39.5 |
| **Non-locoregional lymph ganglia invasion (%)** | 19.2 | 35.3 | 25.6 |
| **Perineural invasion (%)** | 30.8 | 35.3 | 32.6 |
| **Peritoneal metastasis (%)** | 50 | 52.9 | 51.2 |
| **Vascular invasion (%)** | 34.6 | 35.3 | 34.9 |

**Table S2. Differentially expressed genes among high and low survival groups.**

**Table S3. Differential splicing events between the two groups in the in-house cohort.**

**Table S4. Association of differential events according to survival with overall survival in the in-house cohort.**

| GeneID | Gene Symbol | logFC | logCPM | P Value |
|---|---|---|---|---|
| ENSG00000007174 | DNAH9 | 2,67586183 | 2,90924416 | 3,38E-07 |
| ENSG00000156414 | TDRD9 | 2,4754724 | 3,07693949 | 4,42E-06 |
| ENSG00000259786 | LINC02109 | 2,98546389 | 1,81942408 | 5,52E-06 |
| ENSG00000279668 | None | 2,78088488 | 2,23607927 | 8,49E-06 |
| ENSG00000169436 | COL22A1 | -4,4423792 | 6,73208855 | 1,29E-05 |
| ENSG00000144485 | HES6 | 2,29416825 | 3,41370498 | 1,59E-05 |
| ENSG00000228933 | None | 2,65612742 | 2,25623964 | 1,81E-05 |
| ENSG00000237268 | None | 3,15504408 | 2,29942693 | 1,87E-05 |
| ENSG00000236333 | TRHDE-AS1 | 2,4222462 | 3,62392526 | 3,50E-05 |
| ENSG00000169245 | CXCL10 | 2,24820281 | 2,40980302 | 5,00E-05 |
| ENSG00000104044 | OCA2 | 2,12286937 | 1,92243031 | 5,30E-05 |
| ENSG00000264940 | SNORD3C | -3,8941822 | 4,47383301 | 6,41E-05 |
| ENSG00000234840 | LINC01239 | 1,98052003 | 1,62241947 | 8,06E-05 |
| ENSG00000197168 | NEK5 | 1,79761826 | 1,28301986 | 8,21E-05 |
| ENSG00000250337 | PURPL | 2,35822589 | 2,80504445 | 8,52E-05 |
| ENSG00000124743 | KLHL31 | 1,55442251 | 1,8724985 | 8,99E-05 |
| ENSG00000197935 | ZNF311 | 1,07610436 | 0,79188561 | 0,00010867 |
| ENSG00000147606 | SLC26A7 | -3,0759399 | 3,38671869 | 0,00012881 |
| ENSG00000262533 | None | -1,7960249 | 2,06231897 | 0,00013528 |
| ENSG00000104722 | NEFM | -5,027181 | 6,26194368 | 0,00014854 |
| ENSG00000196218 | RYR1 | 1,66211302 | 4,52803144 | 0,0001493 |
| ENSG00000198797 | BRINP2 | 2,21740234 | 1,75330401 | 0,00015923 |
| ENSG00000140090 | SLC24A4 | 1,73897716 | 3,16894381 | 0,00016398 |
| ENSG00000124882 | EREG | -2,7309854 | 2,14884282 | 0,00023067 |
| ENSG00000277693 | None | 3,75353127 | 3,32156523 | 0,00025311 |
| ENSG00000241956 | LINC03000 | -4,0590919 | 3,69797547 | 0,0002655 |
| ENSG00000073756 | PTGS2 | -1,3333274 | 4,10222422 | 0,00030191 |
| ENSG00000166819 | PLIN1 | 2,16923887 | 5,42929776 | 0,000303 |
| ENSG00000148344 | PTGES | 1,71728074 | 2,37279405 | 0,00032517 |
| ENSG00000172508 | CARNS1 | 1,52572866 | 2,61770841 | 0,00057636 |
| ENSG00000185499 | MUC1 | -1,9933409 | 4,55639245 | 0,00060621 |
| ENSG00000007402 | CACNA2D2 | -2,2750862 | 3,4154004 | 0,00063012 |
| ENSG00000176222 | ZNF404 | 1,00084818 | 1,7902824 | 0,00066046 |
| ENSG00000281641 | SAMD12-AS1 | 1,33648147 | 0,85242815 | 0,00071673 |
| ENSG00000237928 | NFIA-AS2 | 2,02782985 | 1,07246434 | 0,00080363 |
| ENSG00000196604 | POTEF | 1,88773274 | 1,58976612 | 0,00082601 |
| ENSG00000152766 | ANKRD22 | 1,49477084 | 1,18180435 | 0,00084359 |
| ENSG00000187987 | ZSCAN23 | 1,04080125 | 2,09541941 | 0,001014 |
| ENSG00000162595 | DIRAS3 | -2,093394 | 5,33241996 | 0,00110946 |
| ENSG00000250696 | None | 1,28955125 | 2,70289872 | 0,00111211 |
| ENSG00000280411 | IGHV1-69D | 2,72199329 | 3,41459269 | 0,00119425 |
| ENSG00000185818 | NAT8L | 1,45896335 | 2,43165275 | 0,00123494 |
| ENSG00000263050 | DPH1-AS1 | -1,8044855 | 2,72515097 | 0,00128637 |
| ENSG00000180730 | SHISA2 | 1,49093337 | 2,67902241 | 0,00131622 |
| ENSG00000131044 | TTLL9 | 1,32886355 | 1,17582825 | 0,00133379 |
| ENSG00000211651 | IGLV1-44 | 2,05548152 | 3,55988086 | 0,00135075 |
| ENSG00000260423 | LINC02367 | 1,19022109 | 1,08461541 | 0,00137129 |
| ENSG00000105409 | ATP1A3 | 1,17539121 | 1,33477599 | 0,00158002 |
| ENSG00000124203 | ZNF831 | 1,61843156 | 5,57836234 | 0,00171831 |
| ENSG00000135218 | CD36 | 1,35341326 | 6,28234471 | 0,00172125 |
| ENSG00000175749 | EIF3KP1 | -1,5823112 | 1,8222704 | 0,00172218 |

| | | | | |
|---|---|---|---|---|
| ENSG00000170323 | FABP4 | 2,17074373 | 4,9223572 | 0,00195291 |
| ENSG00000198010 | DLGAP2 | 1,63236601 | 1,54335581 | 0,00202914 |
| ENSG00000117115 | PADI2 | 1,44401865 | 5,21467951 | 0,00203622 |
| ENSG00000151322 | NPAS3 | -1,5702923 | 3,29693705 | 0,00208352 |
| ENSG00000211951 | IGHV2-26 | 2,74289477 | 2,44764278 | 0,00215084 |
| ENSG00000162444 | RBP7 | 1,24496513 | 1,69340093 | 0,00227747 |
| ENSG00000237781 | ADAMTSL4-AS2 | -1,0510825 | 3,75472289 | 0,00234106 |
| ENSG00000255823 | None | -2,4681954 | 4,16148683 | 0,00236482 |
| ENSG00000250098 | None | -1,0844722 | 0,81349868 | 0,00239485 |
| ENSG00000162722 | TRIM58 | 1,37650069 | 1,67667086 | 0,0024608 |
| ENSG00000186806 | VSIG10L | 1,29073802 | 2,13431847 | 0,0024709 |
| ENSG00000227674 | LINC00355 | 2,38280036 | 1,92215471 | 0,00249715 |
| ENSG00000223039 | RN7SKP268 | -1,1749712 | 1,27133477 | 0,00281991 |
| ENSG00000100314 | CABP7 | -2,1690543 | 1,87537918 | 0,00282542 |
| ENSG00000172901 | LVRN | 1,77190753 | 1,76687958 | 0,00303451 |
| ENSG00000169031 | COL4A3 | -1,4014123 | 4,40511726 | 0,00306669 |
| ENSG00000173208 | ABCD2 | 1,14346258 | 2,07235567 | 0,00309014 |
| ENSG00000227681 | None | -1,5528935 | 1,93734098 | 0,00315317 |
| ENSG00000163749 | CCDC158 | 1,08457704 | 5,07208089 | 0,00330664 |
| ENSG00000102924 | CBLN1 | 1,55942829 | 1,43754798 | 0,0035277 |
| ENSG00000181126 | HLA-V | 1,36977399 | 1,88490369 | 0,00365334 |
| ENSG00000110148 | CCKBR | -1,9818669 | 1,52034911 | 0,00374438 |
| ENSG00000108231 | LGI1 | -2,2661703 | 2,57085 | 0,00387025 |
| ENSG00000203727 | SAMD5 | -1,4186892 | 5,53129653 | 0,00412517 |
| ENSG00000132854 | KANK4 | 1,28375226 | 2,36903881 | 0,00433515 |
| ENSG00000250350 | None | -1,2994336 | 1,92939038 | 0,00437937 |
| ENSG00000105696 | TMEM59L | 1,28687533 | 2,64551391 | 0,00461883 |
| ENSG00000211662 | IGLV3-21 | 1,99963831 | 4,99628953 | 0,00480281 |
| ENSG00000188596 | CFAP54 | 1,14070751 | 3,33476812 | 0,00484499 |
| ENSG00000250155 | SLC1A3-AS1 | 1,24095158 | 1,11862405 | 0,0051623 |
| ENSG00000251095 | None | 1,48282115 | 3,0941756 | 0,00554103 |
| ENSG00000162772 | ATF3 | -1,1928964 | 5,81162749 | 0,00559726 |
| ENSG00000173805 | HAP1 | -2,1832807 | 3,86312394 | 0,00584645 |
| ENSG00000226674 | TEX41 | 1,24563714 | 3,95556704 | 0,00620858 |
| ENSG00000139292 | LGR5 | -2,0091953 | 2,44249515 | 0,00627592 |
| ENSG00000270550 | IGHV3-30 | 1,64100421 | 3,78097812 | 0,00636696 |
| ENSG00000185985 | SLITRK2 | 1,67884976 | 1,78696538 | 0,0063767 |
| ENSG00000224373 | IGHV4-59 | 1,75323641 | 3,66862005 | 0,00670845 |
| ENSG00000138755 | CXCL9 | 1,5100478 | 4,0432123 | 0,00694903 |
| ENSG00000211673 | IGLV3-1 | 1,76550057 | 3,85872111 | 0,00696517 |
| ENSG00000092758 | COL9A3 | 1,43672874 | 2,48139147 | 0,00740992 |
| ENSG00000164604 | GPR85 | 1,01714171 | 1,87793911 | 0,00750055 |
| ENSG00000236699 | ARHGEF38 | -2,5760775 | 3,62780646 | 0,00776146 |
| ENSG00000280281 | None | -1,5012133 | 1,60331362 | 0,0078631 |
| ENSG00000115112 | TFCP2L1 | -1,636158 | 3,92092 | 0,00852822 |
| ENSG00000164266 | SPINK1 | 1,8285605 | 3,98226397 | 0,00891365 |
| ENSG00000280660 | None | -1,9453771 | 2,09296792 | 0,00953192 |
| ENSG00000125740 | FOSB | -1,490275 | 8,47085497 | 0,01055323 |
| ENSG00000116183 | PAPPA2 | -2,3885323 | 5,79647577 | 0,01093026 |
| ENSG00000157423 | HYDIN | 1,14068643 | 2,51028767 | 0,01119396 |
| ENSG00000164619 | BMPER | -1,5085123 | 3,99993393 | 0,01160733 |
| ENSG00000279637 | None | -1,2643859 | 2,90347853 | 0,01184471 |

| | | | | |
|---|---|---|---|---|
| ENSG00000175445 | LPL | 1,28028758 | 5,15209748 | 0,01206753 |
| ENSG00000211898 | IGHD | 1,65924579 | 2,34958066 | 0,0121341 |
| ENSG00000166573 | GALR1 | 1,27821365 | 1,82966638 | 0,01240096 |
| ENSG00000197766 | CFD | 1,17176594 | 4,26499583 | 0,01295976 |
| ENSG00000109321 | AREG | -1,2856897 | 2,10383092 | 0,01310148 |
| ENSG00000183092 | BEGAIN | -1,0360202 | 3,62212849 | 0,01318937 |
| ENSG00000172602 | RND1 | -1,1271155 | 3,56875411 | 0,01330908 |
| ENSG00000276107 | THBS1-IT1 | -1,2479051 | 5,30435238 | 0,01366534 |
| ENSG00000170379 | TCAF2 | -1,3699259 | 3,08894431 | 0,01371611 |
| ENSG00000231561 | CEACAMP5 | 1,20783152 | 1,16667895 | 0,0138565 |
| ENSG00000119973 | PRLHR | -1,3559801 | 5,0044236 | 0,0149263 |
| ENSG00000273604 | EPOP | 1,03002153 | 2,08042602 | 0,01523461 |
| ENSG00000109846 | CRYAB | 1,13816623 | 5,3577552 | 0,015299 |
| ENSG00000105835 | NAMPT | -1,164903 | 9,24769204 | 0,01539563 |
| ENSG00000256654 | None | -1,9062244 | 2,73981255 | 0,01585151 |
| ENSG00000278530 | CHMP1B2P | 1,11262832 | 2,0900915 | 0,01626039 |
| ENSG00000277586 | NEFL | -2,5769516 | 4,72737991 | 0,01654386 |
| ENSG00000273328 | None | 1,15135261 | 2,12626612 | 0,01674805 |
| ENSG00000153822 | KCNJ16 | 1,52670161 | 2,59731125 | 0,01697114 |
| ENSG00000211640 | IGLV6-57 | 1,73229841 | 3,03704641 | 0,01703892 |
| ENSG00000100146 | SOX10 | 1,63486126 | 3,09284645 | 0,01742715 |
| ENSG00000140795 | MYLK3 | -1,5309695 | 2,20527651 | 0,01743659 |
| ENSG00000153303 | FRMD1 | 1,2846197 | 2,06396465 | 0,01817272 |
| ENSG00000173825 | TIGD3 | 1,03437019 | 1,20302705 | 0,01885712 |
| ENSG00000287553 | None | -1,0790153 | 1,78075334 | 0,01914311 |
| ENSG00000279091 | None | -1,5139976 | 2,64550067 | 0,01931362 |
| ENSG00000137265 | IRF4 | 1,20292567 | 4,93221085 | 0,01955329 |
| ENSG00000240086 | None | 1,55350046 | 2,64115988 | 0,01959129 |
| ENSG00000234389 | None | -1,0403499 | 1,3859524 | 0,01959962 |
| ENSG00000211649 | IGLV7-46 | 1,64211227 | 1,75832675 | 0,01989422 |
| ENSG00000155511 | GRIA1 | -1,3697089 | 2,86145477 | 0,02028108 |
| ENSG00000243449 | C4orf48 | 1,21774596 | 0,92719147 | 0,02080322 |
| ENSG00000211964 | IGHV3-48 | 1,57583355 | 3,22588827 | 0,02131068 |
| ENSG00000139304 | PTPRQ | 1,03288706 | 1,22462377 | 0,02131984 |
| ENSG00000099194 | SCD | 1,0172986 | 7,4971346 | 0,02139219 |
| ENSG00000169676 | DRD5 | -1,633475 | 4,42201126 | 0,02182834 |
| ENSG00000196208 | GREB1 | -1,0565869 | 4,23571698 | 0,02204434 |
| ENSG00000241351 | IGKV3-11 | 1,47190955 | 4,57931226 | 0,02232855 |
| ENSG00000244040 | IL12A-AS1 | -1,2032655 | 2,94649574 | 0,02272054 |
| ENSG00000162998 | FRZB | -1,1429058 | 5,3543511 | 0,0228349 |
| ENSG00000211896 | IGHG1 | 1,56648957 | 9,78391707 | 0,02300194 |
| ENSG00000286058 | None | 1,25355315 | 2,09348358 | 0,02323668 |
| ENSG00000007314 | SCN4A | 1,06234724 | 1,85737917 | 0,02327431 |
| ENSG00000233928 | None | 1,39506925 | 1,06278631 | 0,02382446 |
| ENSG00000165023 | DIRAS2 | 1,24000028 | 1,32295333 | 0,02392869 |
| ENSG00000145536 | ADAMTS16 | 1,40851705 | 1,50796511 | 0,02479468 |
| ENSG00000254101 | LINC02055 | 1,54111147 | 2,35529964 | 0,0255965 |
| ENSG00000243466 | IGKV1-5 | 1,27592254 | 4,97577206 | 0,02660538 |
| ENSG00000234604 | None | -1,0611582 | 1,4685325 | 0,02714417 |
| ENSG00000273149 | None | 1,22877301 | 1,44767604 | 0,02796572 |
| ENSG00000181092 | ADIPOQ | 2,3162366 | 5,30181804 | 0,02806948 |
| ENSG00000125538 | IL1B | -1,0358449 | 2,83329157 | 0,02838116 |

| | | | | |
|---|---|---|---|---|
| ENSG00000189430 | NCR1 | -1,9839281 | 4,47769159 | 0,02838237 |
| ENSG00000186188 | FFAR4 | -1,2861437 | 1,30666115 | 0,02861128 |
| ENSG00000233393 | None | 1,1359308 | 1,22557468 | 0,02877938 |
| ENSG00000184811 | TRARG1 | 2,05773441 | 2,70335803 | 0,02883048 |
| ENSG00000186301 | MST1P2 | 1,2969308 | 4,44396774 | 0,02931123 |
| ENSG00000176884 | GRIN1 | -1,4485594 | 0,94524229 | 0,02944585 |
| ENSG00000173599 | PC | 1,17334328 | 5,8240964 | 0,03012739 |
| ENSG00000224769 | MUC20P1 | -1,0540162 | 3,14070009 | 0,0305173 |
| ENSG00000232216 | IGHV3-43 | 1,5076636 | 1,7314914 | 0,03053452 |
| ENSG00000111262 | KCNA1 | -1,842192 | 3,52791474 | 0,03069864 |
| ENSG00000135324 | MRAP2 | -1,3588996 | 2,26254589 | 0,03116062 |
| ENSG00000239951 | IGKV3-20 | 1,3502403 | 5,63673141 | 0,03185431 |
| ENSG00000104267 | CA2 | -1,0893068 | 4,57899731 | 0,03195017 |
| ENSG00000188107 | EYS | 1,01996507 | 2,36309421 | 0,03201085 |
| ENSG00000131409 | LRRC4B | 1,026247 | 3,79031652 | 0,032962 |
| ENSG00000169174 | PCSK9 | 1,63371956 | 3,3071309 | 0,03300786 |
| ENSG00000211965 | IGHV3-49 | 1,3324731 | 2,01686296 | 0,03328547 |
| ENSG00000079931 | MOXD1 | -1,0356657 | 4,6636894 | 0,03347928 |
| ENSG00000172020 | GAP43 | -1,268813 | 2,12531634 | 0,0337522 |
| ENSG00000183196 | CHST6 | -1,0734925 | 4,32598636 | 0,03423728 |
| ENSG00000254042 | SLIT3-AS2 | -1,8664438 | 1,1984341 | 0,0344368 |
| ENSG00000124915 | MYRF-AS1 | -1,0491548 | 1,27157108 | 0,03455303 |
| ENSG00000224223 | VSTM2A-OT1 | 1,27758499 | 2,99678018 | 0,0346488 |
| ENSG00000124780 | KCNK17 | -1,249769 | 1,97670743 | 0,03564553 |
| ENSG00000174697 | LEP | 1,90550223 | 2,83124796 | 0,03590695 |
| ENSG00000239921 | LINC01471 | 1,10398996 | 1,90015839 | 0,03615434 |
| ENSG00000143546 | S100A8 | -1,4082754 | 2,46745352 | 0,03704372 |
| ENSG00000007908 | SELE | -1,0383425 | 2,97408429 | 0,03747784 |
| ENSG00000163661 | PTX3 | -1,0694215 | 1,06540276 | 0,03783034 |
| ENSG00000124731 | TREM1 | -1,226168 | 3,03249875 | 0,03899873 |
| ENSG00000165646 | SLC18A2 | -1,1639005 | 5,77622855 | 0,0405458 |
| ENSG00000086619 | ERO1B | -1,0551159 | 5,84865127 | 0,04104039 |
| ENSG00000115602 | IL1RL1 | -1,2080017 | 4,27106821 | 0,04151003 |
| ENSG00000244731 | C4A | 1,22872951 | 5,60311628 | 0,04228824 |
| ENSG00000187957 | DNER | 1,24149059 | 2,53886594 | 0,04277138 |
| ENSG00000087237 | CETP | 1,26180796 | 1,89765606 | 0,04294777 |
| ENSG00000133454 | MYO18B | 1,18712571 | 2,66565577 | 0,0431686 |
| ENSG00000215478 | CES5AP1 | 1,15962476 | 2,00715399 | 0,04353003 |
| ENSG00000274979 | None | 1,06117057 | 3,52508125 | 0,04366045 |
| ENSG00000257743 | MGAM2 | -1,191081 | 4,69792265 | 0,04374452 |
| ENSG00000260075 | NSFP1 | -1,4471652 | 0,56338122 | 0,04387564 |
| ENSG00000100678 | SLC8A3 | -1,5890509 | 2,02113745 | 0,04406025 |
| ENSG00000123342 | MMP19 | -1,1045893 | 5,87654873 | 0,04430985 |
| ENSG00000165841 | CYP2C19 | -1,1035498 | 6,06079986 | 0,04548602 |
| ENSG00000108342 | CSF3 | -1,4392929 | 1,09516474 | 0,04550954 |
| ENSG00000270933 | None | 1,20512545 | 1,14204162 | 0,04631724 |
| ENSG00000124205 | EDN3 | -2,1429584 | 3,76480901 | 0,04713892 |
| ENSG00000223403 | MEG9 | -1,1065027 | 2,28205977 | 0,04773882 |
| ENSG00000162747 | FCGR3B | -1,1207397 | 1,91561676 | 0,04825278 |
| ENSG00000181143 | MUC16 | 1,08000302 | 1,95051661 | 0,04857915 |
| ENSG00000211950 | IGHV1-24 | 1,57115654 | 3,44177737 | 0,04874069 |
| ENSG00000211941 | IGHV3-11 | 1,30756665 | 2,7615384 | 0,04971335 |

| | | | | |
|---|---|---|---|---|
| ENSG00000205362 | MT1A | 1,22019844 | 1,75170217 | 0,04982925 |

| Gene ID | rMATS name | *p*-value | dPSI |
|---|---|---|---|
| ENSG00000173933 | SE_RBM4_chr11_66652276_66652452_66639699_66640123_66665855_66668374_+ | 3,44169E-15 | -0,227 |
| ENSG00000161016 | SE_RPL8_chr8_144791427_144791872_144791276_144791310_144791991_144792057_- | 3,66374E-15 | 0,12 |
| ENSG00000166435 | MXE_XRRA1_chr11_74859050_74859283_74862980_74863021_74851988_74852082_74906238_74906456_- | 8,77076E-15 | -0,184 |
| ENSG00000284554 | SE_AL022318.4_chr22_39025069_39025349_39022821_39023014_39029362_39029519_+ | 1,31672E-13 | 0,176 |
| ENSG00000095794 | A3SS_CREM_chr10_35211258_35211691_35211655_35211691_35206894_35207051_+ | 1,33227E-13 | -0,105 |
| ENSG00000167840 | RI_ZNF232_chr17_5108925_5109868_5108925_5109052_5109393_5109868_- | 1,59095E-13 | 0,118 |
| ENSG00000130714 | RI_POMT1_chr9_131511336_131512136_131511336_131511472_131512040_131512136_+ | 1,75748E-13 | 0,15 |
| ENSG00000166435 | SE_XRRA1_chr11_74859157_74859283_74848114_74848462_74862980_74863021_- | 4,88165E-13 | 0,14 |
| ENSG00000150527 | SE_MIA2_chr14_39320929_39321056_39319208_39319291_39326863_39327022_+ | 9,20597E-13 | 0,109 |
| ENSG00000172890 | A3SS_NADSYN1_chr11_71476032_71477438_71477344_71477438_71474394_71474526_+ | 3,84393E-12 | 0,103 |
| ENSG00000140521 | RI_POLG_chr15_89317938_89318749_89317938_89318037_89318540_89318749_- | 4,26925E-12 | 0,12 |
| ENSG00000114779 | SE_ABHD14B_chr3_51971459_51971698_51969083_51970184_51973964_51974021_- | 2,98253E-11 | 0,113 |
| ENSG00000223959 | SE_AFG3L1P_chr16_89978472_89978878_89977693_89977802_89980241_89980336_+ | 1,76439E-10 | 0,138 |
| ENSG00000214106 | RI_PAXIP1-AS2_chr7_154947730_154948090_154947730_154947773_154947897_154948090_+ | 3,35795E-10 | 0,14 |
| ENSG00000135486 | SE_HNRNPA1_chr12_54282843_54283234_54281794_54281896_54283826_54283967_+ | 4,36207E-10 | 0,202 |
| ENSG00000135486 | SE_HNRNPA1_chr12_54282843_54283228_54281794_54281896_54283826_54283967_+ | 4,83566E-10 | 0,201 |
| ENSG00000135486 | SE_HNRNPA1_chr12_54282843_54283225_54281794_54281896_54283826_54283967_+ | 4,85339E-10 | 0,201 |
| ENSG00000111271 | MXE_ACAD10_chr12_111749010_111749103_111749152_111749345_111748316_111748475_111755667_111755745_+ | 6,42419E-10 | -0,142 |
| ENSG00000170633 | SE_RNF34_chr12_121404848_121405113_121402741_121402807_121416158_121416377_+ | 1,63253E-09 | -0,165 |
| ENSG00000112511 | RI_PHF1_chr6_33415214_33415670_33415214_33415329_33415592_33415670_+ | 2,22947E-09 | -0,163 |
| ENSG00000126953 | A3SS_TIMM8A_chrX_101345660_101348401_101345660_101346660_101348532_101348742_- | 2,24542E-09 | 0,16 |
| ENSG00000266714 | A3SS_MYO15B_chr17_75594480_75594972_75594839_75594972_75592678_75592840_+ | 3,66611E-09 | -0,135 |
| ENSG00000171574 | SE_ZNF584_chr19_58409940_58410091_58408648_58409165_58415523_58415646_+ | 6,42303E-09 | -0,161 |
| ENSG00000114745 | RI_GORASP1_chr3_39100304_39100877_39100304_39100503_39100787_39100877_- | 1,14662E-08 | 0,12 |
| ENSG00000111271 | MXE_ACAD10_chr12_111749010_111749103_111749172_111749345_111748316_111748475_111755667_111755745_+ | 1,92417E-08 | -0,102 |
| ENSG00000170421 | SE_KRT8_chr12_52948722_52949218_52904675_52905027_52949455_52949842_- | 2,38563E-08 | 0,166 |
| ENSG00000121897 | SE_LIAS_chr4_39468118_39468427_39467517_39467646_39470018_39470164_+ | 4,37133E-08 | 0,143 |
| ENSG00000153774 | SE_CFDP1_chr16_75414577_75414695_75412534_75412754_75433288_75433461_- | 5,30556E-08 | -0,112 |
| ENSG00000170421 | SE_KRT8_chr12_52948722_52948828_52904675_52905027_52949455_52949517_- | 5,82229E-08 | 0,191 |
| ENSG00000163694 | SE_RBM47_chr4_40436440_40436644_40432650_40432862_40437770_40438924_- | 1,05533E-07 | -0,131 |
| ENSG00000185379 | SE_RAD51D_chr17_35116858_35117037_35107365_35107447_35118500_35118619_- | 1,1039E-07 | 0,101 |
| ENSG00000153774 | SE_CFDP1_chr16_75414577_75414694_75412534_75412754_75433288_75433503_- | 1,55471E-07 | -0,148 |
| ENSG00000019144 | SE_PHLDB1_chr11_118613293_118613450_118610426_118610526_118613815_118613896_+ | 1,60417E-07 | 0,144 |
| ENSG00000120451 | SE_SNX19_chr11_130880621_130880796_130879623_130879711_130903254_130903384_- | 1,76147E-07 | 0,195 |
| ENSG00000099330 | A3SS_OCEL1_chr19_17228802_17229219_17229008_17229219_17228255_17228309_+ | 2,04343E-07 | 0,105 |
| ENSG00000143774 | SE_GUK1_chr1_228148370_228148539_228147620_228147699_228148664_228148808_+ | 2,15272E-07 | -0,109 |
| ENSG00000121310 | MXE_ECHDC2_chr1_52907013_52907221_52907623_52907762_52906518_52906611_52907867_52907954_- | 4,21576E-07 | 0,158 |
| ENSG00000121310 | MXE_ECHDC2_chr1_52906981_52907221_52907623_52907762_52906518_52906611_52907867_52907954_- | 4,33192E-07 | 0,161 |
| ENSG00000095059 | SE_DHPS_chr19_12677143_12677211_12676016_12676142_12677290_12677396_- | 5,3824E-07 | 0,114 |
| ENSG00000149657 | MXE_LSM14B_chr20_62127584_62127701_62129784_62129952_62126303_62126439_62130218_62130296_+ | 6,10501E-07 | -0,177 |
| ENSG00000121897 | A5SS_LIAS_chr4_39467517_39468427_39467517_39467646_39470018_39470164_+ | 7,13616E-07 | 0,148 |
| ENSG00000075539 | SE_FRYL_chr4_48710518_48710698_48684672_48684795_48780077_48780248_- | 8,02388E-07 | 0,109 |
| ENSG00000123159 | SE_GIPC1_chr19_14482688_14482819_14480592_14480778_14491655_14491743_- | 8,17672E-07 | 0,141 |
| ENSG00000120451 | SE_SNX19_chr11_130880621_130880805_130879623_130879711_130903254_130903384_- | 9,9946E-07 | 0,146 |
| ENSG00000140474 | A5SS_ULK3_chr15_74837750_74837798_74837756_74837798_74837368_74837435_- | 1,06257E-06 | -0,258 |
| ENSG00000120451 | SE_SNX19_chr11_130880621_130880806_130879623_130879711_130903254_130903384_- | 1,21542E-06 | 0,128 |
| ENSG00000162650 | RI_ATXN7L2_chr1_109488846_109490128_109488846_109489100_109489929_109490128_+ | 1,23455E-06 | 0,114 |
| ENSG00000099330 | A3SS_OCEL1_chr19_17228778_17229218_17229008_17229218_17228255_17228309_+ | 1,59007E-06 | 0,124 |
| ENSG00000141084 | MXE_RANBP10_chr16_67738012_67738035_67744287_67744455_67734857_67735042_67772033_67772086_- | 1,84207E-06 | 0,104 |
| ENSG00000159069 | A5SS_FBXW5_chr9_136941754_136942466_136942099_136942466_136941536_136941684_- | 2,68903E-06 | -0,134 |
| ENSG00000159788 | MXE_RGS12_chr4_3342936_3343053_3386415_3386437_3316067_3318051_3414071_3414241_+ | 5,40501E-06 | -0,13 |
| ENSG00000033867 | SE_SLC4A7_chr3_27379005_27379356_27376763_27376845_27383152_27383250_- | 5,58308E-06 | 0,113 |
| ENSG00000143774 | SE_GUK1_chr1_228148357_228148455_228147620_228147699_228148664_228148984_+ | 6,98768E-06 | -0,251 |
| ENSG00000100401 | SE_RANGAP1_chr22_41284762_41285153_41280932_41281082_41285543_41285965_- | 1,01301E-05 | -0,145 |
| ENSG00000174483 | A5SS_BBS1_chr11_66511021_66511239_66511021_66511089_66514405_66514678_+ | 1,26065E-05 | 0,102 |
| ENSG00000143774 | SE_GUK1_chr1_228148357_228148539_228147620_228147699_228148664_228148808_+ | 1,31766E-05 | -0,206 |
| ENSG00000184465 | RI_WDR27_chr6_169663840_169664286_169663840_169663878_169664165_169664286_- | 2,56468E-05 | 0,121 |
| ENSG00000169231 | RI_THBS3_chr1_155197188_155197659_155197188_155197213_155197462_155197659_- | 3,309E-05 | 0,231 |
| ENSG00000143774 | SE_GUK1_chr1_228148233_228148455_228147614_228147699_228148664_228148984_+ | 3,95245E-05 | -0,204 |
| ENSG00000135486 | A5SS_HNRNPA1_chr12_54282398_54282820_54282398_54282641_54283826_54283967_+ | 4,2615E-05 | -0,107 |
| ENSG00000135486 | A5SS_HNRNPA1_chr12_54282398_54282823_54282398_54282641_54283826_54283967_+ | 4,56795E-05 | -0,108 |
| ENSG00000130635 | A3SS_COL5A1_chr9_134829975_134830176_134830107_134830176_134825791_134825904_+ | 4,60787E-05 | 0,157 |
| ENSG00000135486 | A5SS_HNRNPA1_chr12_54282398_54282841_54282398_54282641_54283826_54283967_+ | 5,23273E-05 | -0,11 |
| ENSG00000017260 | SE_ATP2C1_chr3_130894589_130894775_130894014_130894337_130930415_130930526_+ | 5,50584E-05 | -0,104 |
| ENSG00000141084 | MXE_RANBP10_chr16_67738012_67738035_67744264_67744455_67734857_67735042_67772033_67772086_- | 6,05959E-05 | 0,103 |
| ENSG00000157193 | RI_LRP8_chr1_53271226_53275753_53271226_53271346_53275630_53275753_- | 6,10973E-05 | 0,164 |
| ENSG00000109466 | SE_KLHL2_chr4_165228806_165228913_165219933_165220059_165238777_165238831_+ | 6,22297E-05 | -0,109 |
| ENSG00000114745 | A5SS_GORASP1_chr3_39100746_39100877_39100787_39100877_39100304_39100503_- | 6,54811E-05 | 0,15 |
| ENSG00000022277 | SE_RTF2_chr20_56516943_56516989_56513390_56513428_56517105_56517201_+ | 9,8009E-05 | -0,165 |
| ENSG00000142046 | A3SS_TMEM91_chr19_41383560_41384083_41383862_41384083_41382771_41382921_+ | 0,000112723 | 0,106 |
| ENSG00000170919 | A5SS_TPT1-AS1_chr13_45383090_45383235_45383090_45383169_45383759_45383838_+ | 0,000113588 | -0,185 |
| ENSG00000022277 | A3SS_RTF2_chr20_56516934_56517200_56517105_56517200_56513314_56513428_+ | 0,000123459 | -0,165 |
| ENSG00000185522 | RI_LMNTD2_chr11_555733_556114_555733_555930_555995_556114_- | 0,000139527 | 0,246 |
| ENSG00000163694 | A3SS_RBM47_chr4_40436440_40436647_40436440_40436644_40437770_40438924_- | 0,000151145 | 0,116 |
| ENSG00000223959 | SE_AFG3L1P_chr16_89985445_89985570_89984530_89984599_89988851_89988951_+ | 0,000153873 | 0,156 |
| ENSG00000105339 | MXE_DENND3_chr8_141160631_141160787_141165185_141165289_141155848_141155970_141166189_141166389_+ | 0,000157391 | 0,176 |
| ENSG00000183648 | SE_NDUFB1_chr14_92121195_92121413_92117497_92117642_92121641_92121917_- | 0,000157629 | 0,117 |
| ENSG00000022277 | SE_RTF2_chr20_56516934_56516987_56513314_56513428_56517105_56517201_+ | 0,000177984 | -0,148 |
| ENSG00000137817 | A5SS_PARP6_chr15_72272392_72272554_72272502_72272554_72271022_72271287_- | 0,000202847 | 0,117 |
| ENSG00000186577 | MXE_SMIM29_chr6_34247043_34247149_34247466_34247492_34246394_34246868_34247680_34247864_- | 0,000218413 | -0,109 |
| ENSG00000107404 | RI_DVL1_chr1_1339582_1339812_1339582_1339649_1339735_1339812_- | 0,000218645 | 0,102 |
| ENSG00000122085 | SE_MTERF4_chr2_241091894_241092114_241089979_241090044_241094276_241094432_- | 0,000223833 | 0,146 |
| ENSG00000108039 | SE_XPNPEP1_chr10_109893011_109893073_109891578_109891826_109907690_109907819_- | 0,000254986 | 0,124 |
| ENSG00000141524 | SE_TMC6_chr17_78120652_78120832_78119296_78119392_78121012_78121164_- | 0,000301718 | 0,164 |
| ENSG00000160226 | A5SS_CFAP410_chr21_44334060_44335804_44335757_44335804_44333032_44333262_- | 0,000318762 | 0,11 |
| ENSG00000196756 | A5SS_SNHG17_chr20_38422095_38422984_38422290_38422984_38421005_38421098_- | 0,000345814 | 0,156 |
| ENSG00000134285 | MXE_FKBP11_chr12_48924560_48924648_48924717_48925166_48924222_48924256_48925299_48925485_- | 0,000369023 | 0,142 |
| ENSG00000152223 | A3SS_EPG5_chr18_45882273_45882487_45882273_45882431_45884616_45884811_- | 0,000418612 | -0,109 |
| ENSG00000100401 | SE_RANGAP1_chr22_41284900_41285153_41280932_41281082_41285543_41285965_- | 0,000466072 | -0,128 |
| ENSG00000108592 | RI_FTSJ3_chr17_63820838_63821115_63820838_63820938_63821029_63821115_- | 0,000466551 | -0,106 |

| | | | |
|---|---|---|---|
| ENSG00000133943 | A5SS_DGLUCY_chr14_91215404_91215793_91215404_91215556_91224683_91225355_+ | 0,000492415 | 0,193 |
| ENSG00000166526 | SE_ZNF3_chr7_100079539_100079656_100077302_100077433_100081634_100081703_- | 0,000507905 | -0,119 |
| ENSG00000116754 | SE_SRSF11_chr1_70231094_70231257_70228421_70228555_70231858_70231975_+ | 0,000542496 | 0,148 |
| ENSG00000133943 | SE_DGLUCY_chr14_91223675_91223795_91215404_91215556_91224683_91224822_+ | 0,000567783 | 0,223 |
| ENSG00000155506 | A5SS_LARP1_chr5_154793594_154793901_154793594_154793723_154794099_154794262_+ | 0,00058998 | -0,168 |
| ENSG00000102897 | SE_LYRM1_chr16_20915555_20916047_20902485_20902717_20920121_20920674_+ | 0,000614065 | 0,179 |
| ENSG00000006534 | RI_ALDH3B1_chr11_68015296_68018637_68015296_68015459_68018526_68018637_+ | 0,00062218 | -0,141 |
| ENSG00000162434 | MXE_JAK1_chr1_64886258_64886341_65066627_65066871_64879024_64879148_65067603_65067754_- | 0,00062499 | 0,205 |
| ENSG00000176641 | SE_RNF152_chr18_61890364_61890516_61816121_61816598_61892794_61892917_- | 0,00063292 | 0,216 |
| ENSG00000002933 | SE_TMEM176A_chr7_150804372_150804485_150803619_150803832_150804826_150805113_+ | 0,000638658 | -0,167 |
| ENSG00000115446 | A3SS_UNC50_chr2_98609642_98610039_98609755_98610039_98608588_98608726_+ | 0,000712594 | 0,159 |
| ENSG00000072501 | RI_SMC1A_chrX_53405244_53405671_53405244_53405391_53405492_53405671_- | 0,000741387 | -0,101 |
| ENSG00000163001 | RI_CFAP36_chr2_55543940_55544294_55543940_55544074_55544219_55544294_+ | 0,000746556 | -0,173 |
| ENSG00000180354 | SE_MTURN_chr7_30146176_30146299_30134985_30135298_30157437_30162760_+ | 0,000788305 | -0,11 |
| ENSG00000116754 | SE_SRSF11_chr1_70231094_70231203_70228421_70228555_70231858_70231975_+ | 0,000788424 | 0,132 |
| ENSG00000144455 | MXE_SUMF1_chr3_4362038_4362254_4376329_4376388_4068568_4068745_4410864_4410978_- | 0,000833621 | -0,178 |
| ENSG00000100949 | A5SS_RABGGTA_chr14_24269269_24269692_24269490_24269692_24269079_24269163_- | 0,000867184 | -0,111 |
| ENSG00000235703 | RI_LINC00894_chrX_149948048_149948394_149948048_149948175_149948305_149948394_+ | 0,000872699 | -0,104 |
| ENSG00000006704 | A3SS_GTF2IRD1_chr7_74601024_74601106_74601043_74601106_74595013_74595051_+ | 0,000885989 | 0,117 |
| ENSG00000198276 | RI_UCKL1_chr20_63940816_63941043_63940816_63940856_63940949_63941043_- | 0,000908852 | 0,156 |
| ENSG00000143776 | MXE_CDC42BPA_chr1_227193785_227193934_227199556_227199652_227160542_227160636_227213135_227213219_- | 0,000949874 | 0,126 |
| ENSG00000168268 | RI_NT5DC2_chr3_52527296_52527684_52527296_52527375_52527616_52527684_- | 0,000974704 | -0,114 |
| ENSG00000005810 | SE_MYCBP2_chr13_77118339_77118519_77098067_77099013_77121372_77121495_- | 0,001008806 | 0,192 |
| ENSG00000108963 | RI_DPH1_chr17_2041767_2043898_2041767_2042231_2042604_2043898_+ | 0,001020712 | 0,137 |
| ENSG00000149269 | MXE_PAK1_chr11_77343818_77343931_77355667_77355842_77340645_77340763_77358897_77359017_- | 0,001043847 | 0,11 |
| ENSG00000140632 | A3SS_GLYR1_chr16_4821331_4821593_4821331_4821453_4822874_4822931_- | 0,001086437 | 0,124 |
| ENSG00000198836 | SE_OPA1_chr3_193617177_193617285_193615673_193615770_193618868_193618936_+ | 0,001161562 | 0,112 |
| ENSG00000114391 | A5SS_RPL24_chr3_101682440_101682896_101682770_101682896_101681095_101681215_- | 0,001197597 | -0,123 |
| ENSG00000154380 | MXE_ENAH_chr1_225514595_225514900_225517195_225517306_225512870_225513016_225519197_225519565_- | 0,001283435 | 0,163 |
| ENSG00000170144 | RI_HNRNPA3_chr2_177215977_177216940_177215977_177216771_177216859_177216940_+ | 0,001407183 | -0,121 |
| ENSG00000268750 | SE_AC010522.1_chr19_57840837_57840908_57839022_57839149_57856103_57856233_+ | 0,001445287 | -0,204 |
| ENSG00000221978 | MXE_CCNL2_chr1_1388491_1388510_1388625_1388767_1387953_1388065_1390229_1390371_- | 0,001575858 | -0,202 |
| ENSG00000149657 | MXE_LSM14B_chr20_62127584_62127701_62130218_62130296_62126303_62126439_62130529_62130691_+ | 0,001615768 | -0,155 |
| ENSG00000116455 | SE_WDR77_chr1_111441498_111441604_111441297_111441389_111442024_111442093_- | 0,001632024 | -0,13 |
| ENSG00000060339 | A5SS_CCAR1_chr10_68721226_68721670_68721226_68721282_68722454_68722577_+ | 0,001819757 | 0,114 |
| ENSG00000268750 | SE_AC010522.1_chr19_57840837_57840927_57839022_57839149_57856103_57856233_+ | 0,001900338 | -0,204 |
| ENSG00000140632 | A3SS_GLYR1_chr16_4821331_4821579_4821331_4821453_4822874_4822931_- | 0,001973288 | 0,12 |
| ENSG00000118495 | A5SS_PLAGL1_chr6_143965849_143966198_143966157_143966198_143964786_143964818_- | 0,0020115 | 0,107 |
| ENSG00000122359 | MXE_ANXA11_chr10_80166889_80166984_80167225_80167313_80166083_80166197_80168968_80169358_- | 0,002055676 | 0,103 |
| ENSG00000105552 | SE_BCAT2_chr19_48807755_48808236_48806999_48807074_48810983_48811027_- | 0,002119924 | -0,143 |
| ENSG00000134285 | RI_FKBP11_chr12_48924223_48924615_48924223_48924256_48924560_48924615_- | 0,002129933 | 0,12 |
| ENSG00000185345 | MXE_PRKN_chr6_162201130_162201252_162262524_162262765_162054090_162054174_162443309_162443473_- | 0,002130345 | 0,166 |
| ENSG00000110619 | MXE_CARS1_chr11_3001969_3002174_3002540_3002600_3000929_3001248_3005365_3005433_- | 0,00215229 | -0,225 |
| ENSG00000165282 | RI_PIGO_chr9_35091239_35092767_35091239_35091291_35092542_35092767_- | 0,002349949 | 0,12 |
| ENSG00000125347 | MXE_IRF1_chr5_132484361_132484495_132484548_132484639_132483978_132484075_132485666_132485716_- | 0,002403458 | 0,154 |
| ENSG00000196756 | RI_SNHG17_chr20_38421005_38422368_38421005_38421098_38422290_38422368_- | 0,002433868 | 0,122 |
| ENSG00000186088 | RI_GSAP_chr7_77329332_77330367_77329332_77329391_77330238_77330367_- | 0,002542228 | -0,105 |
| ENSG00000276234 | MXE_TADA2A_chr17_37442563_37442652_37444695_37444768_37440504_37440662_37458523_37458587_+ | 0,002574884 | 0,217 |
| ENSG00000100897 | SE_DCAF11_chr14_24115203_24115293_24114811_24114924_24115542_24115749_+ | 0,002711488 | -0,111 |
| ENSG00000110931 | SE_CAMKK2_chr12_121248605_121248861_121245139_121245240_121249786_121249874_- | 0,00279562 | -0,147 |
| ENSG00000185379 | MXE_RAD51D_chr17_35107365_35107447_35116858_35117037_35106987_35107122_35118500_35118619_- | 0,002807934 | 0,134 |
| ENSG00000105552 | SE_BCAT2_chr19_48807768_48808236_48807012_48807074_48810983_48811027_- | 0,002812574 | -0,144 |
| ENSG00000163516 | RI_ANKZF1_chr2_219234132_219235312_219234132_219234288_219234825_219235312_+ | 0,002833507 | -0,11 |
| ENSG00000196535 | SE_MYO18A_chr17_29085603_29085648_29082315_29082438_29086437_29086577_- | 0,002856164 | 0,11 |
| ENSG00000145012 | SE_LPP_chr3_188341655_188341719_188225404_188225527_188406111_188406313_+ | 0,002876502 | 0,132 |
| ENSG00000108465 | RI_CDK5RAP3_chr17_47975513_47976013_47975513_47975653_47975868_47976013_+ | 0,002884029 | -0,109 |
| ENSG00000117643 | MXE_MAN1C1_chr1_25686439_25686536_25746667_25746782_25616790_25618337_25749254_25749335_+ | 0,002947181 | -0,107 |
| ENSG00000111364 | MXE_DDX55_chr12_123605930_123606159_123607431_123607523_123602098_123602256_123607599_123607662_+ | 0,002956279 | -0,244 |
| ENSG00000174197 | SE_MGA_chr15_41742545_41742724_41739905_41740203_41748636_41748927_+ | 0,00300923 | -0,128 |
| ENSG00000100664 | RI_EIF5_chr14_103334236_103334597_103334236_103334260_103334388_103334597_+ | 0,003015665 | 0,164 |
| ENSG00000088727 | MXE_KIF9_chr3_47277281_47277379_47282494_47283119_47275324_47275490_47283400_47283451_- | 0,003065097 | 0,168 |
| ENSG00000170291 | RI_ELP5_chr17_7258587_7258925_7258587_7258683_7258825_7258925_+ | 0,003209741 | 0,195 |
| ENSG00000124574 | RI_ABCC10_chr6_43437933_43438795_43437933_43438013_43438623_43438795_+ | 0,003267883 | -0,182 |
| ENSG00000145012 | MXE_LPP_chr3_188341655_188341719_188406111_188406157_188225404_188225527_188484591_188484704_+ | 0,003327587 | 0,146 |
| ENSG00000177479 | SE_ARIH2_chr3_48922327_48922811_48918894_48918998_48924717_48924839_+ | 0,003344936 | -0,137 |
| ENSG00000196792 | MXE_STRN3_chr14_30913523_30913657_30929200_30929311_30912006_30912182_30935162_30935304_- | 0,003457187 | 0,139 |
| ENSG00000158552 | RI_ZFAND2B_chr2_219208983_219209418_219208983_219209049_219209261_219209418_+ | 0,003518485 | -0,177 |
| ENSG00000129187 | SE_DCTD_chr4_182916418_182916539_182915460_182915575_182917287_182917389_- | 0,003608804 | -0,149 |
| ENSG00000108848 | MXE_LUC7L3_chr17_50737286_50737479_50738140_50738199_50736959_50737026_50740305_50740345_+ | 0,003648469 | 0,102 |
| ENSG00000160789 | RI_LMNA_chr1_156137653_156138757_156137653_156137743_156138487_156138757_+ | 0,003653588 | 0,122 |
| ENSG00000163755 | A5SS_HPS3_chr3_149129637_149130453_149129637_149129940_149140003_149140498_+ | 0,003674112 | -0,168 |
| ENSG00000272896 | A5SS_AL355987.3_chr9_136791378_136791777_136791378_136791424_136791946_136792030_+ | 0,003692287 | 0,146 |
| ENSG00000116350 | SE_SRSF4_chr1_29168502_29168638_29160374_29160517_29181645_29181890_- | 0,003709281 | 0,114 |
| ENSG00000214176 | RI_PLEKHM1P1_chr17_64821766_64822393_64821766_64822052_64822253_64822393_- | 0,00372294 | -0,129 |
| ENSG00000156858 | SE_PRR14_chr16_30650992_30651127_30650777_30650806_30651595_30651668_+ | 0,003774016 | 0,223 |
| ENSG00000122970 | A5SS_IFT81_chr12_110124334_110124861_110124334_110124443_110127359_110127524_+ | 0,003794355 | 0,126 |
| ENSG00000221978 | MXE_CCNL2_chr1_1388491_1388510_1388625_1388743_1387953_1388065_1390229_1390371_- | 0,003822379 | -0,182 |
| ENSG00000204574 | A3SS_ABCF1_chr6_30578072_30578202_30578075_30578202_30577817_30577913_+ | 0,003996534 | -0,181 |
| ENSG00000203879 | RI_GDI1_chrX_154439940_154440424_154439940_154440139_154440374_154440424_+ | 0,004032063 | -0,127 |
| ENSG00000111481 | MXE_COPZ1_chr12_54345459_54345515_54347766_54347844_54343224_54343316_54347999_54348051_+ | 0,004190879 | -0,108 |
| ENSG00000038532 | A5SS_CLEC16A_chr16_11003073_11003305_11003073_11003257_11020192_11020325_+ | 0,0042618 | 0,18 |
| ENSG00000114735 | RI_HEMK1_chr3_50571715_50572208_50571715_50571801_50572114_50572208_+ | 0,004348948 | 0,169 |
| ENSG00000103145 | RI_HCFC1R1_chr16_3022840_3023298_3022840_3022998_3023232_3023298_- | 0,004350485 | 0,13 |
| ENSG00000115875 | MXE_SRSF7_chr2_38748897_38749173_38749239_38749346_38748578_38748653_38749528_38749705_- | 0,004408492 | 0,104 |
| ENSG00000177575 | RI_CD163_chr12_7479859_7481256_7479859_7479996_7481160_7481256_- | 0,004457377 | 0,15 |
| ENSG00000166436 | SE_TRIM66_chr11_8654433_8654578_8651799_8651903_8671896_8672098_- | 0,004482139 | 0,146 |
| ENSG00000099331 | MXE_MYO9B_chr19_17156908_17157038_17159394_17159484_17154314_17154415_17162349_17162466_+ | 0,004761329 | 0,11 |
| ENSG00000153406 | SE_NMRAL1_chr16_4471313_4471449_4469226_4469465_4474092_4474166_- | 0,004801807 | -0,115 |
| ENSG00000113108 | RI_APBB3_chr5_140560727_140561101_140560727_140560754_140561017_140561101_- | 0,0051348 | 0,128 |
| ENSG00000017260 | SE_ATP2C1_chr3_130894730_130894775_130894181_130894337_130930415_130930526_+ | 0,005307509 | -0,113 |
| ENSG00000109654 | RI_TRIM2_chr4_153322647_153324148_153322647_153322816_153324077_153324148_+ | 0,005386966 | 0,156 |

| | | | |
|---|---|---|---|
| ENSG00000203485 | SE_INF2_chr14_104715832_104716017_104715283_104715340_104718794_104719603_+ | 0,005435821 | 0,126 |
| ENSG00000123684 | SE_LPGAT1_chr1_211830182_211830326_211829058_211829323_211830572_211830763_- | 0,005506777 | 0,177 |
| ENSG00000166689 | RI_PLEKHA7_chr11_16802972_16803295_16802972_16803052_16803226_16803295_- | 0,005524434 | -0,107 |
| ENSG00000221978 | MXE_CCNL2_chr1_1388491_1388767_1388949_1389047_1387953_1388065_1390229_1390371_- | 0,005712807 | -0,141 |
| ENSG00000164654 | MXE_MIOS_chr7_7585614_7585805_7589404_7589563_7583117_7583372_7594979_7595132_+ | 0,005716475 | -0,134 |
| ENSG00000156113 | MXE_KCNMA1_chr10_76974486_76974573_77001406_77001580_76969973_76970067_77008184_77008193_- | 0,005968731 | -0,151 |
| ENSG00000116001 | MXE_TIA1_chr2_70215124_70215471_70216207_70216292_70214348_70214494_70216403_70216499_- | 0,006006677 | -0,148 |
| ENSG00000175575 | MXE_PAAF1_chr11_73900310_73900420_73909398_73909593_73899145_73899244_73914412_73914504_+ | 0,006148995 | -0,108 |
| ENSG00000114841 | RI_DNAH1_chr3_52388417_52388937_52388417_52388609_52388805_52388937_+ | 0,006188369 | 0,15 |
| ENSG00000172785 | MXE_CBWD1_chr9_147900_148040_151304_151427_146101_146158_152033_152078_- | 0,006295607 | 0,115 |
| ENSG00000115875 | SE_SRSF7_chr2_38749239_38749346_38748578_38748653_38749528_38749705_- | 0,006330819 | 0,173 |
| ENSG00000059378 | MXE_PARP12_chr7_140041643_140041839_140046883_140047007_140034234_140034331_140054661_140054763_- | 0,006505701 | -0,103 |
| ENSG00000157654 | SE_PALM2AKAP2_chr9_109942685_109943381_109931926_109932028_110015953_110016039_+ | 0,006640454 | -0,157 |
| ENSG00000079308 | SE_TNS1_chr2_217830366_217830390_217821739_217821938_217831454_217831547_- | 0,00675772 | 0,146 |
| ENSG00000060138 | RI_YBX3_chr12_10701959_10704148_10701959_10702134_10704059_10704148_- | 0,00698067 | -0,106 |
| ENSG00000095203 | RI_EPB41L4B_chr9_109256135_109256473_109256135_109256224_109256392_109256473_- | 0,007048626 | 0,137 |
| ENSG00000100462 | MXE_PRMT5_chr14_22927525_22927614_22928125_22928211_22926701_22926814_22928496_22928615_- | 0,007105506 | 0,122 |
| ENSG00000196365 | RI_LONP1_chr19_5693297_5693768_5693297_5693462_5693551_5693768_- | 0,007394067 | 0,136 |
| ENSG00000127328 | MXE_RAB3IP_chr12_69755570_69755659_69756404_69756663_69755381_69755478_69784719_69784815_+ | 0,007473934 | -0,102 |
| ENSG00000257341 | RI_AL928654.3_chr14_105488332_105488794_105488332_105488516_105488662_105488794_+ | 0,007525332 | 0,102 |
| ENSG00000196756 | SE_SNHG17_chr20_38422091_38422241_38421005_38421098_38426418_38426503_- | 0,007547432 | 0,152 |
| ENSG00000011275 | SE_RNF216_chr7_5741747_5741815_5741578_5741644_5752845_5752979_- | 0,007607738 | 0,147 |
| ENSG00000145012 | MXE_LPP_chr3_188225407_188225527_188341655_188341719_188212703_188212831_188406111_188406313_+ | 0,007806983 | -0,14 |
| ENSG00000145012 | SE_LPP_chr3_188341662_188341719_188225404_188225527_188406111_188406313_+ | 0,00797609 | 0,115 |
| ENSG00000103037 | MXE_SETD6_chr16_58516477_58516672_58516807_58516928_58516201_58516343_58518180_58518231_+ | 0,008202482 | 0,139 |
| ENSG00000288520 | SE_AL357075.4_chr6_154441291_154441347_154433465_154433507_154442087_154442290_- | 0,00837693 | -0,115 |
| ENSG00000180891 | RI_CUEDC1_chr17_57866473_57867415_57866473_57866544_57867356_57867415_- | 0,00850945 | -0,101 |
| ENSG00000064999 | MXE_ANKS1A_chr6_34981689_34981986_34982751_34982827_34970009_34970166_34983112_34983214_+ | 0,008510707 | 0,139 |
| ENSG00000112367 | RI_FIG4_chr6_109735149_109738453_109735149_109735298_109738324_109738453_+ | 0,008533983 | 0,135 |
| ENSG00000001497 | RI_LAS1L_chrX_65524055_65524614_65524055_65524262_65524563_65524614_- | 0,008684819 | -0,144 |
| ENSG00000164715 | MXE_LMTK2_chr7_98203573_98203706_98203943_98204185_98191613_98194572_98205463_98209638_+ | 0,008710505 | 0,148 |
| ENSG00000139436 | A3SS_GIT2_chr12_109947255_109948846_109947255_109947504_109951166_109951316_- | 0,008804472 | -0,14 |
| ENSG00000006704 | A3SS_GTF2IRD1_chr7_74515344_74515596_74515440_74515596_74512829_74512971_+ | 0,008873879 | 0,122 |
| ENSG00000099899 | MXE_TRMT2A_chr22_20115265_20115447_20115671_20115780_20114964_20115079_20116037_20116612_- | 0,009042657 | 0,112 |
| ENSG00000129158 | SE_SERGEF_chr11_17959469_17959634_17878207_17878244_17988596_17988757_- | 0,009250779 | -0,112 |
| ENSG00000158863 | RI_FAM160B2_chr8_22101707_22102061_22101707_22101851_22101975_22102061_+ | 0,009272908 | 0,113 |
| ENSG00000136811 | MXE_ODF2_chr9_128469182_128469353_128471307_128471473_128460941_128461010_128472912_128473042_+ | 0,009330457 | -0,103 |
| ENSG00000278259 | SE_MYO19_chr17_36501851_36501961_36501068_36501235_36503096_36503200_- | 0,009540108 | -0,125 |
| ENSG00000120158 | MXE_RCL1_chr9_4826857_4827033_4833153_4833228_4823547_4823619_4834140_4834265_+ | 0,009628015 | -0,166 |
| ENSG00000102081 | A3SS_FMR1_chrX_147944868_147945051_147944943_147945051_147943130_147943326_+ | 0,009635745 | 0,108 |
| ENSG00000146733 | MXE_PSPH_chr7_56017233_56017379_56019599_56019734_56015022_56015171_56021072_56021231_- | 0,009724932 | 0,155 |
| ENSG00000134285 | MXE_FKBP11_chr12_48924560_48924648_48924717_48925122_48924222_48924256_48925299_48925492_- | 0,009733839 | 0,113 |
| ENSG00000138760 | SE_SCARB2_chr4_76180210_76180354_76179516_76179705_76180953_76181101_- | 0,00989492 | 0,107 |
| ENSG00000196756 | SE_SNHG17_chr20_38422086_38422241_38421005_38421098_38426418_38426503_- | 0,010095718 | 0,143 |
| ENSG00000108509 | A3SS_CAMTA2_chr17_4969148_4969549_4969148_4969337_4969629_4969701_- | 0,010386155 | 0,157 |
| ENSG00000161265 | RI_U2AF1L4_chr19_35744069_35744421_35744069_35744145_35744322_35744421_- | 0,010549846 | -0,154 |
| ENSG00000102901 | RI_CENPT_chr16_67832011_67832545_67832011_67832108_67832454_67832545_- | 0,01066987 | 0,116 |
| ENSG00000169919 | MXE_GUSB_chr7_65976014_65976202_65979833_65979911_65974918_65975071_65980223_65980409_- | 0,010893594 | -0,129 |
| ENSG00000164494 | MXE_PDSS2_chr6_107210438_107210570_107212108_107212282_107193821_107193854_107245547_107245619_- | 0,011025862 | -0,11 |
| ENSG00000115760 | SE_BIRC6_chr2_32435495_32435585_32433643_32433804_32436052_32436184_+ | 0,011071393 | 0,111 |
| ENSG00000010803 | SE_SCMH1_chr1_41152598_41152741_41142877_41143112_41159724_41159933_- | 0,011211021 | 0,164 |
| ENSG00000168610 | RI_STAT3_chr17_42316788_42317224_42316788_42316851_42317181_42317224_- | 0,011319045 | 0,11 |
| ENSG00000048740 | MXE_CELF2_chr10_11321206_11321386_11325835_11325979_11314138_11314270_11328925_11329097_+ | 0,011401286 | 0,154 |
| ENSG00000002933 | SE_TMEM176A_chr7_150804372_150804719_150803619_150803832_150804826_150805118_+ | 0,011498554 | -0,187 |
| ENSG00000155508 | A5SS_CNOT8_chr5_154858644_154858833_154858644_154858768_154863206_154863395_+ | 0,011533146 | 0,147 |
| ENSG00000090565 | MXE_RAB11FIP3_chr16_461403_461497_471294_471389_425648_426720_482524_482736_+ | 0,011550358 | -0,111 |
| ENSG00000176986 | SE_SEC24C_chr10_73762097_73762166_73760712_73760849_73763489_73763555_+ | 0,011741687 | 0,143 |
| ENSG00000127527 | SE_EPS15L1_chr19_16377121_16377254_16361578_16361984_16385128_16385211_- | 0,01182308 | -0,136 |
| ENSG00000165699 | A3SS_TSC1_chr9_132909806_132910692_132909806_132910689_132911001_132911113_- | 0,011845046 | -0,146 |
| ENSG00000165915 | RI_SLC39A13_chr11_47415038_47415433_47415038_47415159_47415287_47415433_+ | 0,011881774 | 0,111 |
| ENSG00000108825 | MXE_PTGES3L-AARSD1_chr17_42951799_42951894_42954875_42954967_42950525_42950728_42955157_42955224_- | 0,011922141 | -0,116 |
| ENSG00000100124 | MXE_ANKRD54_chr22_37832957_37833109_37833158_37833206_37832636_37832744_37833683_37833755_- | 0,011930513 | 0,134 |
| ENSG00000053254 | SE_FOXN3_chr14_89190386_89190449_89180700_89180806_89280949_89281014_- | 0,011934387 | 0,111 |
| ENSG00000089159 | SE_PXN_chr12_120216272_120216581_120215559_120215661_120221622_120221758_- | 0,011973143 | -0,102 |
| ENSG00000204394 | RI_VARS1_chr6_31780873_31781123_31780873_31780938_31781018_31781123_- | 0,012084254 | 0,101 |
| ENSG00000198258 | RI_UBL5_chr19_9827891_9828376_9827891_9828064_9828326_9828376_+ | 0,012084951 | 0,122 |
| ENSG00000126226 | A5SS_PCID2_chr13_113190567_113190971_113190871_113190971_113185484_113185560_- | 0,012113092 | -0,148 |
| ENSG00000090006 | SE_LTBP4_chr19_40622384_40622667_40619346_40619493_40623603_40623732_+ | 0,012146518 | -0,101 |
| ENSG00000105755 | MXE_ETHE1_chr19_43511436_43511566_43526514_43526659_43508774_43508864_43527096_43527201_- | 0,012295257 | 0,111 |
| ENSG00000164654 | MXE_MIOS_chr7_7585635_7585805_7589404_7589563_7582953_7583372_7594979_7595132_+ | 0,012441697 | -0,13 |
| ENSG00000198561 | SE_CTNND1_chr11_57791491_57791701_57761800_57762046_57794009_57794081_+ | 0,012536852 | 0,109 |
| ENSG00000198399 | MXE_ITSN2_chr2_24313459_24313523_24315131_24315224_24312211_24312375_24328051_24328100_- | 0,012581798 | 0,106 |
| ENSG00000111877 | SE_MCM9_chr6_118932606_118932740_118931419_118931738_118934890_118935024_- | 0,012609746 | -0,217 |
| ENSG00000126934 | RI_MAP2K2_chr19_4099573_4100072_4099573_4099851_4099989_4100072_- | 0,012798847 | -0,111 |
| ENSG00000090006 | SE_LTBP4_chr19_40622367_40622667_40619346_40619493_40623603_40623732_+ | 0,012874847 | -0,103 |
| ENSG00000108509 | A3SS_CAMTA2_chr17_4969148_4969520_4969148_4969337_4969629_4969701_- | 0,012953135 | 0,128 |
| ENSG00000163785 | A3SS_RYK_chr3_134191848_134191986_134191848_134191974_134195081_134195339_- | 0,013050472 | -0,14 |
| ENSG00000133392 | RI_MYH11_chr16_15724646_15724909_15724646_15724799_15724887_15724909_- | 0,013125414 | -0,141 |
| ENSG00000140386 | MXE_SCAPER_chr15_76404523_76404679_76434077_76434310_76379708_76381615_76471211_76471335_- | 0,01333231 | 0,124 |
| ENSG00000204842 | MXE_ATXN2_chr12_111554157_111554217_111597748_111597993_111552905_111552977_111598783_111599676_- | 0,013364968 | 0,116 |
| ENSG00000198561 | SE_CTNND1_chr11_57791493_57791701_57761796_57762046_57794009_57794081_+ | 0,013511628 | 0,108 |
| ENSG00000213066 | SE_CEP43_chr6_167027866_167027993_167026546_167026615_167032602_167032642_+ | 0,013546684 | 0,155 |
| ENSG00000142871 | RI_CCN1_chr1_85581364_85582284_85581364_85581578_85581927_85582284_+ | 0,013634473 | -0,127 |
| ENSG00000108239 | MXE_TBC1D12_chr10_94497054_94497172_94500220_94500327_94493364_94493447_94507266_94507347_+ | 0,013697357 | 0,165 |
| ENSG00000118873 | MXE_RAB3GAP2_chr1_220210387_220210489_220210800_220210876_220205906_220206006_220210954_220211002_- | 0,013754792 | -0,111 |
| ENSG00000128298 | RI_BAIAP2L2_chr22_38087123_38088964_38087123_38088747_38088964_- | 0,013771021 | -0,204 |
| ENSG00000198089 | A5SS_SFI1_chr22_31604286_31604671_31604286_31604404_31604868_31604945_+ | 0,013774501 | 0,123 |
| ENSG00000025708 | RI_TYMP_chr22_50529526_50529990_50529526_50529719_50529903_50529990_- | 0,013826652 | 0,112 |
| ENSG00000084072 | A5SS_PPIE_chr1_39745374_39748767_39745374_39745498_39748902_39749088_+ | 0,014054133 | -0,173 |
| ENSG00000165795 | RI_NDRG2_chr14_21022466_21022905_21022466_21022497_21022863_21022905_- | 0,01423047 | -0,114 |
| ENSG00000095637 | SE_SORBS1_chr10_95414493_95414655_95410632_95410777_95421960_95422073_- | 0,014462894 | 0,119 |

| | | | |
|---|---|---|---|
| ENSG00000090006 | SE_LTBP4_chr19_40622400_40622613_40619346_40619493_40623603_40623932_+ | 0,014516668 | -0,129 |
| ENSG00000134262 | MXE_AP4B1_chr1_113896257_113896465_113897839_113897943_113895756_113896038_113898717_113898801_- | 0,014735179 | -0,131 |
| ENSG00000130733 | MXE_YIPF2_chr19_10925685_10925773_10927629_10927716_10924075_10924192_10927798_10927959_- | 0,014789865 | 0,126 |
| ENSG00000141027 | SE_NCOR1_chr17_16198600_16198719_16194461_16194639_16215361_16215530_- | 0,01507282 | -0,121 |
| ENSG00000166762 | MXE_CATSPER2_chr15_43647916_43648066_43649726_43649879_43647419_43647467_43664417_43664543_- | 0,015085143 | -0,15 |
| ENSG00000164715 | MXE_LMTK2_chr7_98203573_98203706_98203943_98204186_98191613_98194572_98205463_98209638_+ | 0,015278211 | 0,132 |
| ENSG00000103061 | A3SS_SLC7A6OS_chr16_68302380_68302509_68302380_68302501_68304025_68304195_- | 0,015653074 | -0,165 |
| ENSG00000135679 | MXE_MDM2_chr12_68813553_68813628_68828770_68828931_68809211_68809292_68839273_68839849_+ | 0,015746669 | -0,11 |
| ENSG00000082805 | MXE_ERC1_chr12_1115865_1116033_1141619_1141766_1110191_1110347_1180539_1180677_+ | 0,015750969 | -0,11 |
| ENSG00000112146 | RI_FBXO9_chr6_53092733_53093561_53092733_53092824_53093465_53093561_+ | 0,0157592 | -0,133 |
| ENSG00000077549 | SE_CAPZB_chr1_19448833_19449330_19419660_19419750_19485435_19485513_- | 0,01576073 | 0,11 |
| ENSG00000135679 | MXE_MDM2_chr12_68813553_68813628_68828770_68828954_68809207_68809292_68835828_68835984_+ | 0,015835647 | -0,113 |
| ENSG00000075407 | SE_ZNF37A_chr10_38094930_38095224_38094333_38094490_38095537_38095559_+ | 0,015841326 | -0,123 |
| ENSG00000105339 | MXE_DENND3_chr8_141155848_141155970_141160631_141160787_141151618_141151837_141163332_141163429_+ | 0,016163989 | -0,195 |
| ENSG00000136933 | MXE_RABEPK_chr9_125220538_125220700_125227909_125228059_125213369_125213522_125232595_125232745_+ | 0,016246778 | 0,138 |
| ENSG00000005810 | SE_MYCBP2_chr13_77118339_77118495_77098067_77099013_77121372_77121495_- | 0,016369957 | 0,157 |
| ENSG00000165792 | RI_METTL17_chr14_20994793_20995233_20994793_20994901_20995164_20995233_+ | 0,016562173 | -0,192 |
| ENSG00000149269 | MXE_PAK1_chr11_77349238_77349287_77358897_77359017_77343818_77343931_77374327_77374365_- | 0,016634786 | 0,101 |
| ENSG00000108963 | RI_DPH1_chr17_2041767_2043898_2041767_2041875_2042604_2043898_+ | 0,016729469 | 0,12 |
| ENSG00000170502 | A5SS_NUDT9_chr4_87422588_87423004_87422588_87422617_87434980_87435220_+ | 0,016878345 | 0,141 |
| ENSG00000138592 | SE_USP8_chr15_50484274_50484361_50481480_50482065_50489800_50489881_+ | 0,016929306 | 0,139 |
| ENSG00000124570 | SE_SERPINB6_chr6_2966399_2966590_2959167_2959342_2968778_2970745_- | 0,016960789 | 0,103 |
| ENSG00000214078 | RI_CPNE1_chr20_35626043_35626803_35626043_35626381_35626566_35626803_- | 0,017027964 | -0,104 |
| ENSG00000197948 | SE_FCHSD1_chr5_141647967_141648426_141647397_141647520_141648956_141649020_- | 0,017078562 | 0,15 |
| ENSG00000095637 | SE_SORBS1_chr10_95414498_95414862_95410632_95410777_95421960_95422073_- | 0,017383954 | 0,128 |
| ENSG00000116001 | MXE_TIA1_chr2_70215124_70215471_70216207_70216292_70214363_70214491_70216403_70216499_- | 0,017573374 | -0,14 |
| ENSG00000140526 | SE_ABHD2_chr15_89103902_89104299_89088421_89088563_89113724_89113824_+ | 0,017785959 | 0,108 |
| ENSG00000227372 | A5SS_TP73-AS1_chr1_3739835_3740387_3740123_3740387_3737538_3739551_- | 0,017854513 | -0,104 |
| ENSG00000102897 | SE_LYRM1_chr16_20915555_20915713_20902485_20902717_20920121_20920674_+ | 0,01787196 | 0,119 |
| ENSG00000225470 | SE_JPX_chrX_73946998_73947374_73944326_73944662_73994839_73994961_+ | 0,017941706 | -0,143 |
| ENSG00000111859 | MXE_NEDD9_chr6_11192344_11192446_11193590_11193692_11189963_11191205_11213280_11213727_- | 0,018100007 | 0,122 |
| ENSG00000196756 | SE_SNHG17_chr20_38422095_38422241_38421005_38421098_38426418_38426503_- | 0,01812703 | 0,137 |
| ENSG00000119912 | A3SS_IDE_chr10_92479276_92480916_92479276_92479421_92483254_92483337_- | 0,018151772 | 0,16 |
| ENSG00000102796 | SE_DHRS12_chr13_51797825_51797908_51791164_51791257_51799533_51799667_- | 0,018243052 | 0,128 |
| ENSG00000145020 | RI_AMT_chr3_49418970_49421572_49418970_49420342_49421491_49421572_- | 0,018252695 | -0,105 |
| ENSG00000233429 | SE_HOTAIRM1_chr7_27099073_27099366_27096232_27096388_27099778_27099966_+ | 0,018287206 | 0,142 |
| ENSG00000182872 | RI_RBM10_chrX_47179879_47180283_47179879_47180040_47180211_47180283_+ | 0,018326059 | -0,177 |
| ENSG00000134531 | SE_EMP1_chr12_13199177_13199533_13196725_13196872_13211468_13211520_+ | 0,018453687 | 0,15 |
| ENSG00000169228 | RI_RAB24_chr5_177302127_177302496_177302127_177302178_177302396_177302496_- | 0,018527984 | -0,12 |
| ENSG00000185513 | A3SS_L3MBTL1_chr20_43515918_43516177_43516092_43516177_43515291_43515415_+ | 0,01857865 | -0,128 |
| ENSG00000197249 | SE_SERPINA1_chr14_94388566_94389010_94383170_94383241_94390456_94390635_- | 0,018612489 | 0,157 |
| ENSG00000144824 | MXE_PHLDB2_chr3_111948931_111949075_111952571_111952712_111945267_111945357_111953929_111954029_+ | 0,018839973 | -0,107 |
| ENSG00000115648 | SE_MLPH_chr2_237534563_237534647_237527376_237527516_237540347_237540446_+ | 0,018878715 | -0,126 |
| ENSG00000047365 | MXE_ARAP2_chr4_36212395_36212487_36213242_36213319_36210389_36210743_36214421_36214480_- | 0,019093992 | -0,119 |
| ENSG00000122882 | MXE_ECD_chr10_73152292_73152421_73154255_73154448_73148275_73148404_73156274_73156453_- | 0,019413328 | 0,107 |
| ENSG00000174483 | SE_BBS1_chr11_66523171_66523365_66521269_66521376_66523455_66523531_+ | 0,019458499 | -0,194 |
| ENSG00000270149 | MXE_AL591806.2_chr1_161021011_161021487_161038550_161038634_161001284_161001353_161038879_161038962_- | 0,019479422 | 0,12 |
| ENSG00000077150 | RI_NFKB2_chr10_102396448_102396823_102396448_102396489_102396724_102396823_+ | 0,01952227 | 0,103 |
| ENSG00000143727 | SE_ACP1_chr2_272036_272065_271865_271939_272191_272305_+ | 0,019615573 | -0,119 |
| ENSG00000270149 | MXE_AL591806.2_chr1_161021014_161021487_161038550_161038634_161001284_161001353_161038879_161038962_- | 0,019778393 | 0,119 |
| ENSG00000099330 | RI_OCEL1_chr19_17226692_17227199_17226692_17226869_17226993_17227199_+ | 0,019801819 | 0,128 |
| ENSG00000162129 | MXE_CLPB_chr11_72358879_72359008_72380280_72380384_72329706_72329804_72402965_72403038_- | 0,01986124 | -0,152 |
| ENSG00000198089 | MXE_SFI1_chr22_31556941_31557059_31561289_31561392_31550253_31550348_31573057_31573214_+ | 0,020171914 | -0,115 |
| ENSG00000269699 | A3SS_ZIM2_chr19_56824261_56824741_56824261_56824427_56826387_56826463_- | 0,020370162 | -0,21 |
| ENSG00000108510 | MXE_MED13_chr17_61968034_61968258_61972726_61972888_61966461_61966651_61982197_61983114_- | 0,020372064 | 0,12 |
| ENSG00000068354 | MXE_TBC1D25_chrX_48544868_48545023_48558896_48559026_48541332_48541442_48559157_48559346_+ | 0,020496428 | 0,105 |
| ENSG00000117758 | MXE_STX12_chr1_27789561_27789631_27793532_27793632_27773218_27773425_27801677_27801815_+ | 0,020523251 | 0,135 |
| ENSG00000164306 | MXE_PRIMPOL_chr4_184682247_184682336_184685411_184685498_184678231_184678394_184685575_184685684_+ | 0,021073491 | -0,142 |
| ENSG00000131899 | RI_LLGL1_chr17_18235469_18236760_18235469_18235537_18236606_18236760_+ | 0,0215793 | 0,236 |
| ENSG00000149923 | A5SS_PPP4C_chr16_30082745_30082963_30082745_30082847_30083393_30083567_+ | 0,021604602 | 0,106 |
| ENSG00000132024 | A5SS_CC2D1A_chr19_13919798_13919951_13919798_13919939_13920556_13920668_+ | 0,021732455 | -0,133 |
| ENSG00000132199 | RI_ENOSF1_chr18_691067_691304_691067_691106_691203_691304_- | 0,021855612 | -0,109 |
| ENSG00000010295 | RI_IFFO1_chr12_6548064_6548545_6548064_6548157_6548424_6548545_- | 0,021855655 | -0,101 |
| ENSG00000119185 | SE_ITGB1BP1_chr2_9419989_9420100_9418625_9418732_9422478_9422566_- | 0,021868694 | -0,138 |
| ENSG00000175198 | SE_PCCA_chr13_100305750_100305834_100302923_100302998_100307191_100307260_+ | 0,021976462 | -0,149 |
| ENSG00000126456 | A3SS_IRF3_chr19_49660712_49660844_49660712_49660828_49661947_49662328_- | 0,022101234 | 0,182 |
| ENSG00000255717 | RI_SNHG1_chr11_62853800_62854064_62853800_62853833_62854017_62854064_- | 0,022362407 | 0,117 |
| ENSG00000130164 | MXE_LDLR_chr19_11123173_11123344_11128007_11128085_11120369_11120522_11129512_11129670_+ | 0,022507707 | 0,149 |
| ENSG00000168385 | MXE_SEPTIN2_chr2_241316421_241316540_241317498_241317599_241316169_241316315_241324215_241324241_+ | 0,022590215 | 0,108 |
| ENSG00000108176 | MXE_DNAJC12_chr10_67811523_67811663_67823313_67823392_67805582_67805787_67837933_67838059_- | 0,022676539 | 0,119 |
| ENSG00000186866 | RI_POFUT2_chr21_45277022_45277410_45277022_45277142_45277232_45277410_- | 0,022718638 | 0,146 |
| ENSG00000064687 | MXE_ABCA7_chr19_1056906_1057429_1058145_1058269_1056329_1056499_1058617_1058747_+ | 0,022739755 | -0,131 |
| ENSG00000151461 | MXE_UPF2_chr10_11999905_12000009_12001675_12001825_11997671_11997757_12004529_12004727_- | 0,022982465 | -0,108 |
| ENSG00000099968 | MXE_BCL2L13_chr22_17655661_17655832_17683213_17683321_17638583_17638886_17688985_17689142_+ | 0,023003675 | -0,105 |
| ENSG00000233429 | SE_HOTAIRM1_chr7_27099097_27099366_27096123_27096388_27099778_27099966_+ | 0,023020149 | 0,134 |
| ENSG00000099992 | SE_TBC1D10A_chr22_30293158_30293453_30292009_30292851_30293650_30294110_- | 0,023133524 | -0,122 |
| ENSG00000144455 | SE_SUMF1_chr3_4362038_4362254_4068568_4068745_4376329_4376389_- | 0,023152097 | 0,127 |
| ENSG00000103994 | A3SS_ZNF106_chr15_42439032_42439813_42439032_42439552_42442072_42442414_- | 0,023287457 | 0,114 |
| ENSG00000128908 | MXE_INO80_chr15_41059866_41059926_41069569_41069665_41058638_41058781_41070466_41070547_- | 0,023432365 | 0,156 |
| ENSG00000227372 | A5SS_TP73-AS1_chr1_3739623_3740387_3740123_3740387_3737538_3739551_- | 0,023503591 | -0,103 |
| ENSG00000120451 | MXE_SNX19_chr11_130907850_130908083_130910017_130910137_130906624_130906721_130910269_130910370_- | 0,023548713 | -0,107 |
| ENSG00000136960 | RI_ENPP2_chr8_119570704_119572250_119570704_119570841_119572175_119572250_- | 0,023566342 | 0,127 |
| ENSG00000151474 | SE_FRMD4A_chr10_13759164_13759506_13747735_13747819_13761646_13761669_- | 0,023571654 | 0,122 |
| ENSG00000127415 | RI_IDUA_chr4_1003026_1003470_1003026_1003157_1003344_1003470_+ | 0,023592248 | 0,109 |
| ENSG00000089094 | RI_KDM2B_chr12_121444449_121445418_121444449_121444536_121445274_121445418_- | 0,023676455 | 0,171 |
| ENSG00000156858 | MXE_PRR14_chr16_30650992_30651127_30651595_30651668_30650777_30650806_30651795_30651964_+ | 0,024175091 | 0,197 |
| ENSG00000148634 | MXE_HERC4_chr10_67936152_67936235_67939587_67939654_67932596_67932780_67940938_67941105_- | 0,024428791 | -0,104 |
| ENSG00000129566 | MXE_TEP1_chr14_20373677_20373810_20374428_20374545_20373506_20373583_20375754_20375868_- | 0,02462122 | 0,102 |
| ENSG00000266714 | RI_MYO15B_chr17_75594480_75594972_75594480_75594759_75594839_75594972_+ | 0,024716125 | -0,105 |
| ENSG00000140365 | RI_COMMD4_chr15_75338061_75338420_75338061_75338133_75338354_75338420_+ | 0,024868968 | 0,143 |
| ENSG00000127663 | MXE_KDM4B_chr19_5071009_5071059_5077366_5077470_5047475_5047669_5082366_5082437_+ | 0,025511509 | 0,125 |

| | | | |
|---|---|---|---|
| ENSG00000103811 | A5SS_CTSH_chr15_78927367_78927766_78927712_78927766_78925333_78925440_- | 0,025555169 | 0,145 |
| ENSG00000103365 | RI_GGA2_chr16_23493359_23494378_23493359_23493458_23494302_23494378_- | 0,026142301 | -0,104 |
| ENSG00000114315 | A5SS_HES1_chr3_194136960_194137407_194136960_194137048_194137682_194138732_+ | 0,026209611 | -0,152 |
| ENSG00000091262 | MXE_ABCC6_chr16_16177451_16177626_16178797_16178965_16175910_16175986_16182411_16182588_- | 0,026755603 | -0,132 |
| ENSG00000104381 | A5SS_GDAP1_chr8_74351273_74351418_74351273_74351321_74360136_74360310_+ | 0,026827636 | -0,144 |
| ENSG00000100207 | SE_TCF20_chr22_42168608_42168736_42160012_42161358_42169846_42169896_- | 0,026873219 | -0,117 |
| ENSG00000141569 | SE_TRIM65_chr17_75891812_75891878_75888962_75891347_75892010_75892185_- | 0,027135566 | -0,121 |
| ENSG00000185485 | SE_SDHAP1_chr3_195984491_195984737_195984111_195984276_195985626_195985789_- | 0,027780827 | 0,178 |
| ENSG00000059378 | MXE_PARP12_chr7_140037714_140037856_140046883_140047007_140034234_140034331_140054661_140054763_- | 0,028256027 | -0,104 |
| ENSG00000110025 | RI_SNX15_chr11_65035519_65038829_65035519_65035663_65038571_65038829_+ | 0,028393067 | -0,136 |
| ENSG00000141027 | SE_NCOR1_chr17_16198507_16198719_16194461_16194639_16215361_16215401_- | 0,028401131 | -0,104 |
| ENSG00000134285 | MXE_FKBP11_chr12_48924560_48924648_48924717_48925111_48924222_48924256_48925299_48925492_- | 0,028604287 | 0,113 |
| ENSG00000072042 | SE_RDH11_chr14_67686083_67686257_67685014_67685204_67690211_67692851_- | 0,028668133 | -0,102 |
| ENSG00000105778 | MXE_AVL9_chr7_32543140_32543261_32544693_32544779_32495488_32495802_32548846_32548918_+ | 0,029429085 | 0,116 |
| ENSG00000162736 | RI_NCSTN_chr1_160356259_160356728_160356259_160356347_160356599_160356728_+ | 0,029589525 | 0,125 |
| ENSG00000269893 | SE_SNHG8_chr4_118278965_118279137_118278754_118278792_118279388_118279431_+ | 0,029775338 | -0,12 |
| ENSG00000089356 | RI_FXYD3_chr19_35123308_35122919_35123308_35122954_35123270_35123308_+ | 0,029979128 | -0,115 |
| ENSG00000135424 | RI_ITGA7_chr12_55696898_55697275_55696898_55697050_55697215_55697275_- | 0,030481826 | 0,15 |
| ENSG00000171824 | RI_EXOSC10_chr1_11080498_11080911_11080498_11080549_11080763_11080911_- | 0,030672438 | -0,109 |
| ENSG00000026036 | RI_RTEL1-TNFRSF6B_chr20_63689749_63690210_63689749_63689865_63690086_63690210_+ | 0,03074532 | 0,146 |
| ENSG00000106948 | RI_AKNA_chr9_114343707_114346009_114343707_114343803_114345862_114346009_- | 0,030907587 | 0,117 |
| ENSG00000114993 | SE_RTKN_chr2_74439550_74439977_74432058_74432666_74440352_74440583_- | 0,03095704 | 0,128 |
| ENSG00000174501 | MXE_ANKRD36C_chr2_95897269_95897342_95902885_95902958_95889965_95889994_95903052_95903081_- | 0,03099729 | 0,112 |
| ENSG00000110090 | MXE_CPT1A_chr11_68775119_68775432_68780639_68780770_68773264_68773429_68781770_68781959_- | 0,031070977 | -0,107 |
| ENSG00000130830 | A3SS_MPP1_chrX_154790982_154791344_154790982_154791068_154791768_154791847_- | 0,031165818 | 0,154 |
| ENSG00000267106 | A5SS_ZNF561-AS1_chr19_9626783_9627806_9626783_9626860_9632227_9632417_+ | 0,031183332 | -0,101 |
| ENSG00000143499 | RI_SMYD2_chr1_214332017_214334308_214332017_214332192_214334199_214334308_+ | 0,031550221 | 0,168 |
| ENSG00000184634 | RI_MED12_chrX_71125353_71125713_71125353_71125495_71125662_71125713_+ | 0,031799537 | -0,117 |
| ENSG00000184216 | MXE_IRAK1_chrX_154014041_154014278_154016031_154016097_154013042_154013433_154016436_154016644_- | 0,031964031 | 0,116 |
| ENSG00000127419 | RI_TMEM175_chr4_947708_948154_947708_947892_948115_948154_+ | 0,032220141 | -0,115 |
| ENSG00000113240 | RI_CLK4_chr5_178617343_178618778_178617343_178617434_178618555_178618778_- | 0,032426579 | -0,131 |
| ENSG00000171634 | SE_BPTF_chr17_67875556_67875744_67874919_67875020_67891843_67892034_+ | 0,032439997 | -0,116 |
| ENSG00000069956 | SE_MAPK6_chr15_52042685_52042833_52019237_52019376_52045829_52046109_+ | 0,032528405 | -0,117 |
| ENSG00000260565 | SE_ERVK13-1_chr16_2672706_2672976_2671344_2672570_2673282_2673427_- | 0,032811721 | -0,162 |
| ENSG00000185015 | MXE_CA13_chr8_85250739_85250937_85259420_85259539_85245457_85245865_85266607_85266703_+ | 0,032967005 | 0,118 |
| ENSG00000140416 | SE_TPM1_chr15_63043705_63043831_63042746_63042943_63044026_63044152_+ | 0,033263237 | 0,136 |
| ENSG00000168016 | SE_TRANK1_chr3_36850153_36851417_36847199_36847346_36851718_36851856_- | 0,033307552 | 0,102 |
| ENSG00000164828 | RI_SUN1_chr7_852608_852952_852608_852667_852809_852952_+ | 0,033495964 | 0,112 |
| ENSG00000120318 | MXE_ARAP3_chr5_141671264_141671387_141671569_141671939_141670511_141670628_141672101_141672301_- | 0,033657462 | -0,116 |
| ENSG00000116171 | MXE_SCP2_chr1_52941795_52941853_52948008_52948080_52927285_52927465_52950754_52950886_+ | 0,033827856 | 0,121 |
| ENSG00000131370 | MXE_SH3BP5_chr3_15304102_15304231_15330503_15330566_15269712_15269877_15332270_15332586_- | 0,034110233 | 0,119 |
| ENSG00000197586 | MXE_ENTPD6_chr20_25218549_25218614_25221231_25221333_25217501_25217581_25222837_25222978_+ | 0,034157897 | -0,151 |
| ENSG00000203485 | SE_INF2_chr14_104695989_104696490_104689620_104689739_104701356_104701756_+ | 0,035256299 | 0,109 |
| ENSG00000136699 | SE_SMPD4_chr2_130159512_130159634_130157250_130157396_130161185_130161272_- | 0,035576962 | -0,139 |
| ENSG00000064932 | MXE_SBNO2_chr19_1109134_1109211_1109291_1109423_1108778_1108969_1109505_1109598_- | 0,03599622 | 0,155 |
| ENSG00000100412 | SE_ACO2_chr22_41477912_41478127_41469127_41469182_41499725_41499862_+ | 0,036162866 | 0,107 |
| ENSG00000054523 | SE_KIF1B_chr1_10325899_10326359_10324757_10324895_10334519_10334638_+ | 0,036258208 | 0,112 |
| ENSG00000103249 | RI_CLCN7_chr16_1449275_1450666_1449275_1449327_1450496_1450666_- | 0,036323208 | 0,104 |
| ENSG00000107164 | RI_FUBP3_chr9_130631160_130631628_130631160_130631293_130631556_130631628_+ | 0,036735879 | -0,135 |
| ENSG00000132970 | SE_WASF3_chr13_26680034_26680201_26676548_26676724_26681053_26681320_+ | 0,036846393 | -0,129 |
| ENSG00000009413 | MXE_REV3L_chr6_111313351_111313489_111315266_111315381_111311068_111311259_111322568_111322678_- | 0,036987818 | 0,101 |
| ENSG00000105127 | SE_AKAP8_chr19_15369066_15369224_15368234_15368322_15370145_15370179_- | 0,037212266 | 0,147 |
| ENSG00000119185 | SE_ITGB1BP1_chr2_9419989_9420100_9418625_9418732_9423372_9423480_- | 0,037281294 | -0,105 |
| ENSG00000104381 | A5SS_GDAP1_chr8_74351273_74351466_74351273_74351321_74360136_74360310_+ | 0,037451314 | -0,121 |
| ENSG00000184216 | MXE_IRAK1_chrX_154014039_154014278_154016031_154016097_154013042_154013343_154016436_154016644_- | 0,037667266 | 0,12 |
| ENSG00000135372 | MXE_NAT10_chr11_34115822_34115884_34118179_34118294_34113715_34113838_34118395_34118503_+ | 0,03776064 | -0,103 |
| ENSG00000164062 | RI_APEH_chr3_49674195_49674621_49674195_49674413_49674488_49674621_+ | 0,037802274 | 0,125 |
| ENSG00000148187 | SE_MRRF_chr9_122286038_122286166_122285168_122285287_122291748_122291798_+ | 0,037961881 | -0,13 |
| ENSG00000143742 | SE_SRP9_chr1_225786861_225786985_225783299_225783368_225789239_225790466_+ | 0,038095868 | 0,152 |
| ENSG00000170004 | A5SS_CHD3_chr17_7907352_7907701_7907352_7907488_7907893_7908019_+ | 0,038398041 | 0,133 |
| ENSG00000077238 | SE_IL4R_chr16_27318634_27318767_27313991_27314020_27340185_27340273_+ | 0,038405166 | 0,126 |
| ENSG00000104774 | MXE_MAN2B1_chr19_12661259_12661376_12663316_12663462_12658427_12658510_12663702_12663835_- | 0,0384198 | 0,113 |
| ENSG00000157483 | MXE_MYO1E_chr15_59178392_59178537_59188117_59188216_59174125_59174240_59195460_59195567_- | 0,038727957 | -0,118 |
| ENSG00000224078 | SE_SNHG14_chr15_25067408_25067531_25064714_25064855_25070119_25070259_+ | 0,038829887 | -0,111 |
| ENSG00000106686 | MXE_SPATA6L_chr9_4656040_4656089_4661898_4662036_4646298_4646528_4666211_4666307_- | 0,038857372 | 0,102 |
| ENSG00000224078 | RI_SNHG14_chr15_25087466_25089866_25087466_25087599_25089613_25089866_+ | 0,039386581 | -0,109 |
| ENSG00000115875 | RI_SRSF7_chr2_38749239_38749705_38749239_38749346_38749528_38749705_- | 0,040332174 | -0,109 |
| ENSG00000081307 | SE_UBA5_chr3_132660898_132661048_132660600_132660698_132665822_132665868_+ | 0,04047385 | -0,111 |
| ENSG00000135423 | A3SS_GLS2_chr12_56475043_56475183_56475043_56475110_56475623_56475682_- | 0,040671245 | 0,122 |
| ENSG00000271723 | MXE_MROH7-TTC4_chr1_54679262_54679439_54682655_54682794_54678741_54678854_54686257_54686448_+ | 0,040756928 | 0,142 |
| ENSG00000100030 | MXE_MAPK1_chr22_21788256_21788388_21788693_21788808_21772872_21772982_21799011_21799128_- | 0,040884551 | -0,119 |
| ENSG00000215788 | MXE_TNFRSF25_chr1_6462628_6462663_6464551_6464719_6461993_6462174_6465439_6465560_- | 0,041083227 | -0,162 |
| ENSG00000286001 | MXE_AC021087.5_chr5_304176_304221_311292_311642_272710_272772_314416_314955_+ | 0,041382438 | -0,145 |
| ENSG00000171469 | MXE_ZNF561_chr19_9617081_9617171_9617620_9617758_9614058_9614103_9618090_9618179_- | 0,041591195 | 0,125 |
| ENSG00000184922 | RI_FMNL1_chr17_45245252_45245731_45245252_45245416_45245631_45245731_+ | 0,042024048 | -0,103 |
| ENSG00000157353 | RI_FCSK_chr16_70478550_70479403_70478550_70478650_70479179_70479403_+ | 0,042234268 | -0,15 |
| ENSG00000077147 | MXE_TM9SF3_chr10_96552932_96553059_96559658_96559736_96551310_96551411_96561977_96562138_- | 0,042268902 | -0,101 |
| ENSG00000244187 | SE_TMEM141_chr9_136791629_136791777_136791403_136791424_136791946_136792030_+ | 0,042478917 | 0,129 |
| ENSG00000132669 | A3SS_RIN2_chr20_19870843_19871111_19871081_19871111_19799621_19799747_+ | 0,042511814 | 0,165 |
| ENSG00000196187 | RI_TMEM63A_chr1_225862462_225862851_225862462_225862578_225862770_225862851_- | 0,042831207 | -0,11 |
| ENSG00000119185 | SE_ITGB1BP1_chr2_9419989_9420418_9418625_9418732_9422478_9423250_- | 0,042938322 | -0,135 |
| ENSG00000175634 | SE_RPS6KB2_chr11_67432355_67432492_67431367_67431515_67432599_67432657_+ | 0,043527744 | -0,132 |
| ENSG00000145020 | MXE_AMT_chr3_49420210_49420342_49421491_49421572_49419709_49419788_49422103_49422271_- | 0,043564653 | 0,123 |
| ENSG00000072135 | A3SS_PTPN18_chr2_130370421_130370782_130370556_130370782_130370127_130370190_+ | 0,043565657 | -0,157 |
| ENSG00000119688 | MXE_ABCD4_chr14_74289482_74289972_74290026_74290118_74288715_74288765_74290290_74290499_- | 0,043673976 | -0,112 |
| ENSG00000224078 | SE_SNHG14_chr15_25067411_25067531_25064714_25064855_25070119_25070259_+ | 0,044154342 | -0,121 |
| ENSG00000108387 | RI_SEPTIN4_chr17_58526219_58526978_58526219_58526313_58526681_58526978_- | 0,044166299 | 0,121 |
| ENSG00000088682 | MXE_COQ9_chr16_57456930_57457015_57458245_57458350_57456503_57456646_57459564_57459720_+ | 0,044168863 | 0,116 |
| ENSG00000172638 | RI_EFEMP2_chr11_65866440_65867079_65866440_65866712_65866844_65867079_- | 0,044340519 | 0,123 |
| ENSG00000173992 | RI_CCS_chr11_66599115_66599636_66599115_66599253_66599458_66599636_+ | 0,044367508 | -0,106 |
| ENSG00000005471 | MXE_ABCB4_chr7_87443662_87443773_87444861_87444975_87443318_87443444_87447033_87447205_- | 0,044368743 | -0,202 |

| | | | |
|---|---|---|---|
| ENSG00000115419 | A5SS_GLS_chr2_190895603_190895871_190895603_190895725_190900563_190900693_+ | 0,044939549 | 0,143 |
| ENSG00000213782 | RI_DDX47_chr12_12823202_12824016_12823202_12823319_12823869_12824016_+ | 0,045144418 | 0,109 |
| ENSG00000145916 | A5SS_RMND5B_chr5_178138107_178138396_178138107_178138258_178142582_178142728_+ | 0,045363883 | 0,115 |
| ENSG00000137266 | SE_SLC22A23_chr6_3276680_3276976_3273355_3273412_3283851_3284120_- | 0,045462839 | 0,147 |
| ENSG00000115761 | MXE_NOL10_chr2_10675793_10675871_10681970_10682069_10673519_10673557_10684566_10684612_- | 0,045665367 | 0,15 |
| ENSG00000140521 | RI_POLG_chr15_89317375_89318749_89317375_89317536_89317938_89318749_- | 0,045666335 | 0,116 |
| ENSG00000127616 | SE_SMARCA4_chr19_11047594_11047795_11041438_11041560_11058257_11058363_+ | 0,045723204 | 0,186 |
| ENSG00000171720 | RI_HDAC3_chr5_141628113_141628639_141628113_141628187_141628558_141628639_- | 0,046148311 | 0,128 |
| ENSG00000156504 | A3SS_FAM122B_chrX_134769566_134772283_134769566_134771354_134781820_134781917_- | 0,046210734 | 0,112 |
| ENSG00000111679 | RI_PTPN6_chr12_6951608_6952177_6951608_6951731_6951982_6952177_+ | 0,046332565 | -0,188 |
| ENSG00000141504 | RI_SAT2_chr17_7627362_7627802_7627362_7627414_7627552_7627802_- | 0,046641645 | 0,132 |
| ENSG00000089820 | MXE_ARHGAP4_chrX_153910921_153910999_153911128_153911189_153910699_153910834_153912699_153912802_- | 0,04670542 | -0,102 |
| ENSG00000137074 | RI_APTX_chr9_32973375_32974561_32973375_32973652_32974457_32974561_- | 0,047304618 | 0,125 |
| ENSG00000247774 | MXE_PCED1B-AS1_chr12_47209280_47209458_47216164_47216264_47205899_47206585_47216383_47216452_- | 0,047506845 | 0,12 |
| ENSG00000108963 | A5SS_DPH1_chr17_2041767_2042231_2041767_2041875_2042604_2043898_+ | 0,048233441 | 0,11 |
| ENSG00000171466 | MXE_ZNF562_chr19_9658008_9658135_9659378_9659467_9656546_9656653_9660719_9660874_- | 0,048329489 | -0,102 |
| ENSG00000139436 | MXE_GIT2_chr12_109945259_109945349_109947255_109947504_109939164_109939247_109951166_109951316_- | 0,048900781 | 0,103 |
| ENSG00000175634 | MXE_RPS6KB2_chr11_67431367_67431515_67432599_67432657_67429526_67429595_67432736_67432837_+ | 0,049010477 | 0,142 |
| ENSG00000123136 | RI_DDX39A_chr19_14409741_14410334_14409741_14409873_14410215_14410334_- | 0,049025443 | 0,141 |
| ENSG00000064687 | MXE_ABCA7_chr19_1057313_1057429_1058145_1058269_1056906_1057084_1058617_1058747_+ | 0,04907229 | -0,176 |
| ENSG00000166411 | SE_IDH3A_chr15_78150803_78150960_78149376_78149430_78155212_78155275_+ | 0,049079678 | -0,117 |
| ENSG00000196924 | MXE_FLNA_chrX_154357433_154357446_154357447_154357622_154354824_154355072_154358198_154358355_- | 0,049318003 | -0,107 |
| ENSG00000197548 | MXE_ATG7_chr3_11364658_11364734_11379971_11380052_11362812_11362928_11426803_11426926_+ | 0,049792858 | 0,126 |
| ENSG00000227345 | SE_PARG_chr10_49922546_49922665_49922335_49922419_49932099_49932283_- | 0,04981713 | -0,109 |

| Event | P value | Hazard Ratio |
|---|---|---|
| A3SS_ZIM2_56824261_56824741 | 1,96E-06 | 0,066229179 |
| A3SS_L3MBTL1_43515918_43516177 | 0,026829029 | 0,317628872 |
| A3SS_ZNF106_42439032_42439813 | 0,010567186 | 6,252055445 |
| A3SS_GLS2_56475043_56475183 | 0,265796112 | 0,481241937 |
| A3SS_ABCF1_30578072_30578202 | 0,003566595 | 0,217506043 |
| A3SS_IRF3_49660712_49660844 | 0,157508314 | 2,123226943 |
| A3SS_GTF2IRD1_74515344_74515596 | 0,347701467 | 0,593012896 |
| A3SS_OCEL1_17228802_17229219 | 0,040609149 | 4,736362416 |
| A3SS_UNC50_98609642_98610039 | 0,035218703 | 295615758 |
| A3SS_EPG5_45882273_45882487 | 0,04769686 | 0,276815694 |
| A3SS_IDE_92479276_92480916 | 0,110465827 | 3,281789681 |
| A3SS_FMR1_147944868_147945051 | 0,115336552 | 2,430190835 |
| A3SS_NADSYN1_71476032_71477438 | 0,099804754 | 2,944813432 |
| A3SS_COL5A1_134829975_134830176 | 0,012747733 | 4,192654763 |
| A3SS_TIMM8A_101345660_101348401 | 0,125513177 | 2,667575731 |
| A3SS_FAM122B_134769566_134772283 | 0,263953546 | 0,418062181 |
| A3SS_GIT2_109947255_109948846 | 0,048030502 | 0,293953127 |
| A3SS_TMEM91_41383560_41384083 | 0,241468583 | 0,314627924 |
| A3SS_TSC1_132909806_132910692 | 0,03957597 | 0,326882497 |
| A3SS_RTF2_56516934_56517200 | 0,099826541 | 0,300600876 |
| A3SS_SLC7A6OS_68302380_68302509 | 0,104742507 | 5,015760288 |
| A3SS_RBM47_40436440_40436647 | 0,227015095 | 2,11018565 |
| A3SS_CAMTA2_4969148_4969520 | 0,244193755 | 2,161694709 |
| A3SS_CAMTA2_4969148_4969549 | 0,244193755 | 2,161694709 |
| A3SS_GTF2IRD1_74601024_74601106 | 0,129266917 | 0,402446416 |
| A3SS_OCEL1_17228778_17229218 | 0,040609149 | 4,736362416 |
| A3SS_PTPN18_130370421_130370782 | 0,260520746 | 0,326064499 |
| A3SS_RYK_134191848_134191986 | 0,066484308 | 0,339562688 |
| A3SS_GLYR1_4821331_4821579 | 0,050324438 | 3,788837786 |
| A3SS_GLYR1_4821331_4821593 | 0,073640284 | 3,91106319 |
| A3SS_RIN2_19870843_19871111 | 0,327287971 | 0,373779946 |
| A3SS_MYO15B_75594480_75594972 | 0,295404834 | 1,884234293 |
| A3SS_MPP1_154790982_154791344 | 0,136667387 | 4,11E-09 |
| A3SS_CREM_35211258_35211691 | 0,208748929 | 0,477520992 |
| A5SS_ZNF561_AS1_9626783_9627806 | 0,38417849 | 0,598633628 |
| A5SS_SNHG17_38422095_38422984 | 0,00257163 | 5,229969009 |
| A5SS_DPH1_2041767_2042231 | 0,051280906 | 274209595,7 |
| A5SS_PPP4C_30082745_30082963 | 0,267105611 | 0,552398594 |
| A5SS_CLEC16A_11003073_11003305 | 0,088056175 | 0,370050585 |
| A5SS_ULK3_74837750_74837798 | 0,1293062 | 0,382274454 |
| A5SS_HPS3_149129637_149130453 | 0,056146813 | 0,250316822 |
| A5SS_IFT81_110124334_110124861 | 0,068419457 | 3,510229948 |
| A5SS_GDAP1_74351273_74351418 | 0,062527469 | 3,60E-09 |
| A5SS_GDAP1_74351273_74351466 | 0,074808651 | 3,72E-09 |
| A5SS_RMND5B_178138107_178138396 | 0,05795611 | 5,730316118 |
| A5SS_PPIE_39745374_39748767 | 0,009480592 | 0,203418895 |
| A5SS_TP73_AS1_3739623_3740387 | 0,008948714 | 0,181031375 |
| A5SS_TP73_AS1_3739835_3740387 | 0,007044673 | 0,148318489 |
| A5SS_LIAS_39467517_39468427 | 0,107830434 | 2,741642658 |
| A5SS_PLAGL1_143965849_143966198 | 0,240844356 | 2,092975497 |
| A5SS_GORASP1_39100746_39100877 | 0,105965741 | 3,399721654 |
| A5SS_TPT1_AS1_45383090_45383235 | 0,343125505 | 1,741075843 |

| | | |
|---|---|---|
| A5SS_FBXW5_136941754_136942466 | 0,007403331 | 0,238908649 |
| A5SS_DGLUCY_91215404_91215793 | 0,162437862 | 2,103541857 |
| A5SS_CFAP410_44334060_44335804 | 0,454374127 | 1,551190373 |
| A5SS_SFI1_31604286_31604671 | 0,187620589 | 2,160522775 |
| A5SS_CC2D1A_13919798_13919951 | 0,061806402 | 0,351590816 |
| A5SS_CHD3_7907352_7907701 | 5,89E-05 | 9,629930827 |
| A5SS_CTSH_78927367_78927766 | 0,210060875 | 2,16924229 |
| A5SS_RABGGTA_24269269_24269692 | 0,145401922 | 0,389616828 |
| A5SS_NUDT9_87422588_87423004 | 0,42288515 | 1,31E-08 |
| A5SS_CNOT8_154858644_154858833 | 0,327824528 | 1,705499483 |
| A5SS_HES1_194136960_194137407 | 0,002264129 | 0,180958413 |
| A5SS_RPL24_101682440_101682896 | 0,191694172 | 0,435412647 |
| A5SS_PCID2_113190567_113190971 | 0,042369999 | 0,345466254 |
| A5SS_CCAR1_68721226_68721670 | 0,344638058 | 2,038086132 |
| A5SS_HNRNPA1_54282398_54282820 | 0,357028646 | 0,478827706 |
| A5SS_HNRNPA1_54282398_54282823 | 0,357028646 | 0,478827706 |
| A5SS_HNRNPA1_54282398_54282841 | 0,357028646 | 0,478827706 |
| A5SS_LARP1_154793594_154793901 | 0,048599939 | 0,155839232 |
| A5SS_PARP6_72272392_72272554 | 0,002329126 | 5,620242634 |
| A5SS_GLS_190895603_190895871 | 0,112977512 | 2,501880142 |
| A5SS_AL355987_3_136791378_13679177 | 0,000507203 | 6,064904892 |
| A5SS_BBS1_66511021_66511239 | 0,236593379 | 2,134685086 |
| MXE_ANKRD54_37833158_37833206 | 0,226129272 | 1,975435445 |
| MXE_SFI1_31556941_31557059 | 0,003516853 | 0,154875766 |
| MXE_MAPK1_21788693_21788808 | 0,226201338 | 3,288406757 |
| MXE_MYO9B_17156908_17157038 | 0,12960785 | 0,420797054 |
| MXE_MAN2B1_12663316_12663462 | 0,160122921 | 2,625574075 |
| MXE_LDLR_11123173_11123344 | 7,61E-05 | 13,12679158 |
| MXE_SBNO2_1109291_1109423 | 0,132459464 | 2,222509854 |
| MXE_ENTPD6_25218549_25218614 | 0,009214703 | 0,239932002 |
| MXE_MED13_61972726_61972888 | 0,039632223 | 3,590116271 |
| MXE_LUC7L3_50737286_50737479 | 0,030258327 | 3,721763859 |
| MXE_TADA2A_37442563_37442652 | 0,197941441 | 2,376756738 |
| MXE_YIPF2_10927629_10927716 | 0,000546242 | 5,614403374 |
| MXE_ABCC6_16178797_16178965 | 0,015449201 | 0,19045575 |
| MXE_PRR14_30650992_30651127 | 2,46E-05 | 10,15037077 |
| MXE_RAB11FIP3_461403_461497 | 0,212286525 | 0,489219216 |
| MXE_INO80_41069569_41069665 | 0,022336864 | 3,434587161 |
| MXE_TRMT2A_20115671_20115780 | 0,019597507 | 4,351924548 |
| MXE_KDM4B_5071009_5071059 | 0,178275179 | 0,463381243 |
| MXE_SPATA6L_4661898_4662036 | 0,056735307 | 3,294122247 |
| MXE_LSM14B_62127584_62127701 | 6,65E-06 | 0,087000706 |
| MXE_LSM14B_62127584_62127701_1 | 0,000262323 | 0,132499308 |
| MXE_NEDD9_11193590_11193692 | 0,000392723 | 7,668025483 |
| MXE_PCED1B_AS1_47216164_47216264 | 0,059633388 | 5,726930561 |
| MXE_CDC42BPA_227199556_227199652 | 0,007667044 | 5,433453116 |
| MXE_RANBP10_67744264_67744455 | 0,010177746 | 3,894885165 |
| MXE_ANKRD54_37833158_37833206_1 | 0,010177746 | 3,894885165 |
| MXE_ANKRD54_37833158_37833206_2 | 0,003975766 | 0,162648799 |
| MXE_ANKRD54_37833158_37833206_3 | 0,421843811 | 1,606439053 |
| MXE_ANKRD54_37833158_37833206_4 | 0,01199426 | 4,134502778 |
| MXE_ANKRD54_37833158_37833206_5 | 0,05523319 | 0,357850686 |
| MXE_ANKRD54_37833158_37833206_6 | 0,001983239 | 6,059232572 |

| | | |
|---|---|---|
| MXE_ANKRD54_37833158_37833206_7 | 0,001162748 | 5,716336946 |
| MXE_ANKRD54_37833158_37833206_8 | 0,333404342 | 1,677482262 |
| MXE_ANKRD54_37833158_37833206_9 | 0,661230442 | 0,750741436 |
| MXE_ANKRD54_37833158_37833206_10 | 0,002451931 | 5,537907874 |
| MXE_ANKRD54_37833158_37833206_11 | 0,223625189 | 0,47466008 |
| MXE_ANKRD54_37833158_37833206_12 | 0,056714136 | 3,745416353 |
| MXE_ANKRD54_37833158_37833206_13 | 0,263580795 | 3,036352121 |
| MXE_ANKRD54_37833158_37833206_14 | 0,214674913 | 0,472607012 |
| MXE_ANKRD54_37833158_37833206_15 | 0,016373668 | 5,171986253 |
| MXE_ANKRD54_37833158_37833206_16 | 0,053414357 | 3,254876917 |
| MXE_ANKRD54_37833158_37833206_17 | 0,399125949 | 0,603186564 |
| MXE_ANKRD54_37833158_37833206_18 | 0,162945836 | 0,390289666 |
| MXE_ANKRD54_37833158_37833206_19 | 0,140644961 | 2,196358227 |
| MXE_ANKRD54_37833158_37833206_20 | 0,018764527 | 8,864786825 |
| MXE_ANKRD54_37833158_37833206_21 | 0,025686502 | 4,809266684 |
| MXE_ANKRD54_37833158_37833206_22 | 0,020793748 | 0,20243422 |
| MXE_ANKRD54_37833158_37833206_23 | 0,010466043 | 0,203232926 |
| MXE_ANKRD54_37833158_37833206_24 | 0,053658708 | 0,303945874 |
| MXE_ANKRD54_37833158_37833206_25 | 0,218343954 | 2,565417825 |
| MXE_ANKRD54_37833158_37833206_26 | 0,093142997 | 0,400985487 |
| MXE_ANKRD54_37833158_37833206_27 | 0,16321156 | 2,175845478 |
| MXE_ANKRD54_37833158_37833206_28 | 0,201407004 | 2,327234183 |
| MXE_ANKRD54_37833158_37833206_29 | 0,000327833 | 9,477386045 |
| MXE_ANKRD54_37833158_37833206_30 | 0,195846617 | 0,379252721 |
| MXE_ANKRD54_37833158_37833206_31 | 0,029495682 | 3,12E-09 |
| MXE_ANKRD54_37833158_37833206_32 | 0,094521629 | 0,375839412 |
| MXE_ANKRD54_37833158_37833206_33 | 0,007714686 | 0,221112834 |
| MXE_ANKRD54_37833158_37833206_34 | 0,121370156 | 245659229,7 |
| MXE_ANKRD54_37833158_37833206_35 | 0,278667208 | 1,945833414 |
| MXE_ANKRD54_37833158_37833206_36 | 0,021301778 | 0,294822409 |
| MXE_ANKRD54_37833158_37833206_37 | 0,001178967 | 6,342834366 |
| MXE_ANKRD54_37833158_37833206_38 | 0,001297816 | 6,492737542 |
| MXE_ANKRD54_37833158_37833206_39 | 0,003749647 | 0,210908759 |
| MXE_ANKRD54_37833158_37833206_40 | 0,0467959 | 0,353860763 |
| MXE_ANKRD54_37833158_37833206_41 | 0,037113585 | 3,290539206 |
| MXE_ANKRD54_37833158_37833206_42 | 0,070522338 | 0,353051143 |
| MXE_ANKRD54_37833158_37833206_43 | 0,045125547 | 0,323078753 |
| MXE_ANKRD54_37833158_37833206_44 | 0,125137379 | 0,227777616 |
| MXE_ANKRD54_37833158_37833206_45 | 0,047016797 | 0,163424641 |
| MXE_ANKRD54_37833158_37833206_46 | 0,353733276 | 1,649263895 |
| MXE_ANKRD54_37833158_37833206_47 | 0,074811494 | 2,577310169 |
| MXE_ANKRD54_37833158_37833206_48 | 0,026006726 | 4,126680822 |
| MXE_ANKRD54_37833158_37833206_49 | 0,042373494 | 3,230283038 |
| MXE_ANKRD54_37833158_37833206_50 | 0,121383975 | 4,37801911 |
| MXE_ANKRD54_37833158_37833206_51 | 0,091859416 | 2,711923298 |
| MXE_ANKRD54_37833158_37833206_52 | 0,003978613 | 0,174883756 |
| MXE_ANKRD54_37833158_37833206_53 | 0,074776874 | 2,850631193 |
| MXE_ANKRD54_37833158_37833206_54 | 0,017222979 | 5,95842055 |
| MXE_ANKRD54_37833158_37833206_55 | 8,16E-05 | 0,115526408 |
| MXE_ANKRD54_37833158_37833206_56 | 0,002641404 | 6,202782721 |
| MXE_ANKRD54_37833158_37833206_57 | 0,02375227 | 7,713548835 |
| MXE_ANKRD54_37833158_37833206_58 | 0,052944272 | 3,625026393 |
| MXE_ANKRD54_37833158_37833206_59 | 0,20336595 | 1,968179012 |

| | | |
|---|---|---|
| MXE_ANKRD54_37833158_37833206_60 | 2,23E-05 | 0,039341415 |
| MXE_ANKRD54_37833158_37833206_61 | 0,002961797 | 0,125028604 |
| MXE_ANKRD54_37833158_37833206_62 | 0,073420929 | 0,347990792 |
| MXE_ANKRD54_37833158_37833206_63 | 0,027548927 | 4,922918605 |
| MXE_ANKRD54_37833158_37833206_64 | 6,77E-05 | 0,107892809 |
| MXE_ANKRD54_37833158_37833206_65 | 0,091278329 | 0,309441122 |
| MXE_ANKRD54_37833158_37833206_66 | 0,006821137 | 5,425317847 |
| MXE_ANKRD54_37833158_37833206_67 | 0,089304699 | 3,065383293 |
| MXE_ANKRD54_37833158_37833206_68 | 0,003781422 | 5,898349101 |
| MXE_ANKRD54_37833158_37833206_69 | 0,429110515 | 0,597713193 |
| MXE_ANKRD54_37833158_37833206_70 | 0,214242077 | 2,247424097 |
| MXE_ANKRD54_37833158_37833206_71 | 0,02744393 | 5,379206539 |
| MXE_ANKRD54_37833158_37833206_72 | 0,184206763 | 2,146961485 |
| MXE_ANKRD54_37833158_37833206_73 | 0,039660211 | 3,097323555 |
| MXE_ANKRD54_37833158_37833206_74 | 0,07455986 | 0,3796514 |
| MXE_ANKRD54_37833158_37833206_75 | 0,063110973 | 0,374834264 |
| MXE_ANKRD54_37833158_37833206_76 | 0,004673191 | 0,217269916 |
| MXE_ANKRD54_37833158_37833206_77 | 0,02441608 | 3,673014065 |
| MXE_ANKRD54_37833158_37833206_78 | 0,00436947 | 0,142545571 |
| MXE_ANKRD54_37833158_37833206_79 | 0,000280592 | 7,543059773 |
| MXE_ANKRD54_37833158_37833206_80 | 0,337123135 | 0,378702607 |
| MXE_ANKRD54_37833158_37833206_81 | 0,167239956 | 0,451293443 |
| MXE_ANKRD54_37833158_37833206_82 | 0,049768086 | 0,350113256 |
| MXE_ANKRD54_37833158_37833206_83 | 0,098738682 | 2,606982851 |
| MXE_ANKRD54_37833158_37833206_84 | 0,081414027 | 0,39876517 |
| MXE_ANKRD54_37833158_37833206_85 | 0,512897283 | 1,678669251 |
| MXE_ANKRD54_37833158_37833206_86 | 0,024269046 | 0,304234478 |
| MXE_ANKRD54_37833158_37833206_87 | 0,01594443 | 4,98122719 |
| MXE_ANKRD54_37833158_37833206_88 | 0,052388376 | 0,246483469 |
| MXE_ANKRD54_37833158_37833206_89 | 0,006196694 | 5,427775077 |
| MXE_ANKRD54_37833158_37833206_90 | 0,129573927 | 0,445061474 |
| MXE_ANKRD54_37833158_37833206_91 | 0,031669996 | 3,797353508 |
| MXE_ANKRD54_37833158_37833206_92 | 0,031669996 | 3,797353508 |
| MXE_ANKRD54_37833158_37833206_93 | 0,03620724 | 2,988962021 |
| MXE_ANKRD54_37833158_37833206_94 | 0,007974295 | 4,88552846 |
| MXE_ANKRD54_37833158_37833206_95 | 0,076212856 | 2,759608536 |
| MXE_ANKRD54_37833158_37833206_96 | 0,076212856 | 2,759608536 |
| MXE_ANKRD54_37833158_37833206_97 | 0,075005604 | 0,341661517 |
| MXE_ANKRD54_37833158_37833206_98 | 0,068624033 | 2,922524167 |
| MXE_ANKRD54_37833158_37833206_99 | 0,043815138 | 0,315427664 |
| MXE_ANKRD54_37833158_37833206_100 | 0,36338217 | 0,59893935 |
| MXE_ANKRD54_37833158_37833206_101 | 0,057437939 | 3,53E-09 |
| MXE_ANKRD54_37833158_37833206_102 | 0,195421596 | 2,279907263 |
| MXE_ANKRD54_37833158_37833206_103 | 0,020893668 | 5,281461125 |
| MXE_ANKRD54_37833158_37833206_104 | 0,023323658 | 8,513377402 |
| MXE_ANKRD54_37833158_37833206_105 | 0,05346549 | 0,362792896 |
| MXE_ANKRD54_37833158_37833206_106 | 0,002961797 | 0,125028604 |
| MXE_ANKRD54_37833158_37833206_107 | 0,165023555 | 0,4331125 |
| MXE_ANKRD54_37833158_37833206_108 | 0,025868574 | 0,230636628 |
| MXE_ANKRD54_37833158_37833206_109 | 0,000104781 | 8,347111468 |
| MXE_ANKRD54_37833158_37833206_110 | 0,179791075 | 0,488975927 |
| MXE_ANKRD54_37833158_37833206_111 | 0,000481048 | 7,14072698 |
| MXE_ANKRD54_37833158_37833206_112 | 0,006383363 | 0,227386016 |

| | | |
|---|---|---|
| MXE_ANKRD54_37833158_37833206_113 | 0,001226964 | 6,485496429 |
| MXE_ANKRD54_37833158_37833206_114 | 0,001226964 | 6,485496429 |
| RI_BAIAP2L2_38087264_38088747 | 0,02732494 | 0,321823771 |
| RI_U2AF1L4_35744145_35744322 | 0,001382909 | 0,07545218 |
| RI_RTEL1_TNFRSF6B_63689865_636900 | 0,030982893 | 3,018165036 |
| RI_SNHG17_38421098_38422290 | 0,00107017 | 5,855261988 |
| RI_PLEKHM1P1_64822052_64822253 | 0,004995635 | 5,013866751 |
| RI_FTSJ3_63820938_63821029 | 0,169882159 | 0,356481593 |
| RI_CUEDC1_57866544_57867356 | 0,001503416 | 0,191288347 |
| RI_LLGL1_18235537_18236606 | 0,018695512 | 4,619326497 |
| RI_ZNF232_5109052_5109393 | 0,000396695 | 7,564960867 |
| RI_DPH1_2041875_2042604 | 0,051280906 | 274209595,7 |
| RI_DPH1_2042231_2042604 | 0,025045117 | 7,689046627 |
| RI_FCSK_70478650_70479179 | 0,01938389 | 0,236316011 |
| RI_CLCN7_1449327_1450496 | 0,071342122 | 2,860356186 |
| RI_CPNE1_35626381_35626566 | 0,232827601 | 2,51744741 |
| RI_POLG_89317536_89317938 | 0,078333521 | 2,658883929 |
| RI_POLG_89318037_89318540 | 0,033554406 | 3,510551864 |
| RI_COMMD4_75338133_75338354 | 0,008664564 | 4,952155106 |
| RI_CENPT_67832108_67832454 | 0,025225928 | 5,347783923 |
| RI_EIF5_103334260_103334388 | 0,086789942 | 2,972401166 |
| RI_ATXN7L2_109489100_109489929 | 0,234455061 | 1,895391282 |
| RI_CLK4_178617434_178618555 | 0,084253305 | 0,39176953 |
| RI_HNRNPA3_177216771_177216859 | 0,084062117 | 0,37068076 |
| RI_ENPP2_119570841_119572175 | 0,048929298 | 4,480183809 |
| RI_TRIM2_153322816_153324077 | 0,080872633 | 3,965822742 |
| RI_TMEM175_947892_948115 | 0,011034729 | 0,175116699 |
| RI_IFFO1_6548157_6548424 | 0,008962656 | 0,096910455 |
| RI_OCEL1_17226869_17226993 | 0,321789052 | 1,740995493 |
| RI_SUN1_852667_852809 | 0,002164381 | 14,3362308 |
| RI_FBXO9_53092824_53093465 | 0,062546574 | 0,347948765 |
| RI_DDX39A_14409873_14410215 | 0,005492333 | 5,144302739 |
| RI_LMNA_156137743_156138487 | 0,005378702 | 5,233767221 |
| RI_DNAH1_52388609_52388805 | 0,021413269 | 0,199968869 |
| RI_CCN1_85581578_85581927 | 0,026559772 | 0,298339818 |
| RI_APTX_32973652_32974457 | 0,000572758 | 6,290615062 |
| RI_METTL17_20994901_20995164 | 0,002200396 | 0,170623859 |
| RI_STAT3_42316851_42317181 | 0,325831778 | 1,28E-08 |
| RI_LAS1L_65524262_65524563 | 0,0896071 | 3,86E-09 |
| RI_AMT_49420342_49421491 | 0,063623812 | 0,260413175 |
| RI_LRP8_53271346_53275630 | 0,019229746 | 6,145591282 |
| RI_ABCC10_43438013_43438623 | 0,020592177 | 0,24698941 |
| RI_RAB24_177302178_177302396 | 0,303818492 | 1,82850521 |
| RI_GGA2_23493458_23494302 | 0,001227952 | 0,115051591 |
| RI_YBX3_10702134_10704059 | 0,16201361 | 0,47550512 |
| RI_SAT2_7627414_7627552 | 0,005604786 | 5,089037203 |
| RI_NFKB2_102396489_102396724 | 0,010587383 | 4,966244015 |
| RI_FIG4_109735298_109738324 | 0,186198383 | 1,13E-08 |
| RI_HDAC3_141628187_141628558 | 0,10280463 | 2,451757414 |
| RI_SEPTIN4_58526313_58526681 | 0,126320107 | 0,32372603 |
| RI_SNHG14_25087599_25089613 | 0,010042931 | 2,32E-09 |
| RI_SMYD2_214332192_214334199 | 0,05820102 | 4,14761776 |
| RI_ANKZF1_219234288_219234825 | 0,091427084 | 0,369150882 |

| | | |
|---|---|---|
| RI_CDK5RAP3_47975653_47975868 | 0,267142071 | 1,830106682 |
| RI_PTPN6_6951731_6951982 | 0,04782054 | 0,311846052 |
| RI_GORASP1_39100503_39100787 | 0,138444132 | 2,682604083 |
| RI_EFEMP2_65866712_65866844 | 9,58E-05 | 9,006316627 |
| RI_GSAP_77329391_77330238 | 0,009965435 | 0,210839268 |
| RI_ALDH3B1_68015459_68018526 | 0,15531895 | 2,325424896 |
| RI_DDX47_12823319_12823869 | 0,221383898 | 0,509849297 |
| RI_PIGO_35091291_35092542 | 0,03366413 | 3,061456483 |
| RI_AKNA_114343803_114345862 | 0,075110341 | 3,41963799 |
| RI_POMT1_131511472_131512040 | 0,07215811 | 2,920061149 |
| RI_SNHG1_62853833_62854017 | 0,066237376 | 4,132341996 |
| RI_SNX15_65035663_65038571 | 0,108943837 | 0,41771502 |
| RI_CCS_66599253_66599458 | 0,130196147 | 0,410647101 |
| RI_CD163_7479996_7481160 | 0,041328552 | 3,24766811 |
| RI_KDM2B_121444536_121445274 | 0,002637381 | 5,35360629 |
| RI_POFUT2_45277142_45277232 | 0,165813811 | 2,198377551 |
| RI_FXYD3_35122954_35123270 | 0,004739724 | 0,216758593 |
| RI_LONP1_5693462_5693551 | 0,31636412 | 0,517532966 |
| RI_UCKL1_63940856_63940949 | 0,098809667 | 2,899102182 |
| RI_AL928654_3_105488516_105488662 | 0,024522517 | 3,622729185 |
| RI_NDRG2_21022497_21022863 | 0,069669254 | 0,170854435 |
| RI_CFAP36_55544074_55544219 | 0,000110395 | 0,097734897 |
| RI_PHF1_33415329_33415592 | 0,077652147 | 0,36295508 |
| RI_NT5DC2_52527375_52527616 | 0,065517603 | 0,342498797 |
| RI_NCSTN_160356347_160356599 | 0,004262306 | 13,06966406 |
| RI_DVL1_1339649_1339735 | 0,022296666 | 5,561636901 |
| RI_EPB41L4B_109256224_109256392 | 0,019371718 | 3,867686468 |
| RI_PAXIP1_AS2_154947773_154947897 | 0,064421523 | 0,174234093 |
| RI_TMEM63A_225862578_225862770 | 0,025799016 | 2,46E-09 |
| RI_SMC1A_53405391_53405492 | 0,000284158 | 0,119957126 |
| RI_THBS3_155197213_155197462 | 0,137929525 | 4,10E-09 |
| RI_ZFAND2B_219209049_219209261 | 2,02E-05 | 0,081023785 |
| RI_TYMP_50529719_50529903 | 0,003781422 | 5,898349101 |
| RI_APBB3_140560754_140561017 | 0,137045169 | 2,315772389 |
| RI_MYH11_15724799_15724887 | 0,257001372 | 0,542630681 |
| RI_FAM160B2_22101851_22101975 | 0,159738026 | 2,335968943 |
| RI_EXOSC10_11080549_11080763 | 0,022979001 | 0,244624972 |
| RI_SRSF7_38749346_38749528 | 0,177945551 | 0,487475556 |
| RI_GDI1_154440139_154440374 | 0,002460115 | 0,144435267 |
| RI_HEMK1_50571801_50572114 | 0,013529385 | 0,225385589 |
| RI_FMNL1_45245416_45245631 | 0,075161001 | 3,020249078 |
| RI_LINC00894_149948175_149948305 | 0,025640191 | 0,280276891 |
| RI_HCFC1R1_3022998_3023232 | 0,038882447 | 6,586987044 |
| RI_MED12_71125495_71125662 | 0,013275752 | 0,253061885 |
| RI_APEH_49674413_49674488 | 0,461092314 | 0,669438018 |
| RI_PLEKHA7_16803052_16803226 | 0,029935236 | 0,306818929 |
| RI_RBM10_47180040_47180211 | 0,030605164 | 0,314451218 |
| RI_IDUA_1003157_1003344 | 0,190125178 | 2,706613512 |
| RI_MYO15B_75594759_75594839 | 0,110539697 | 0,203282634 |
| RI_VARS1_31780938_31781018 | 0,351813282 | 1,926993553 |
| RI_WDR27_169663878_169664165 | 0,358251019 | 1,756206019 |
| RI_FUBP3_130631293_130631556 | 0,129930656 | 0,406101599 |
| RI_LMNTD2_555930_555995 | 0,055772514 | 3,924863132 |

| | | |
|---|---|---|
| RI_SLC39A13_47415159_47415287 | 0,344385324 | 0,598413267 |
| RI_ELP5_7258683_7258825 | 0,032638082 | 3,193330468 |
| RI_MAP2K2_4099851_4099989 | 0,165957194 | 2,314090776 |
| RI_ENOSF1_691106_691203 | 0,016549352 | 0,207743018 |
| RI_ITGA7_55697050_55697215 | 0,00120585 | 7,414218429 |
| RI_UBL5_9828064_9828326 | 0,224388544 | 0,498471338 |
| RI_FKBP11_48924256_48924560 | 0,004822476 | 4,273401508 |
| SE_TCF20_42168608_42168736 | 0,31058681 | 1,76942754 |
| SE_RANGAP1_41284900_41285153 | 0,124410082 | 2,692043104 |
| SE_AL022318_4_39025069_39025349 | 0,10008506 | 3,67E-09 |
| SE_BCAT2_48807755_48808236 | 0,080515757 | 0,381452906 |
| SE_EPS15L1_16377121_16377254 | 9,80E-06 | 0,037268727 |
| SE_AKAP8_15369066_15369224 | 0,138839225 | 4,14394322 |
| SE_SMARCA4_11047594_11047795 | 0,259858647 | 1,862554675 |
| SE_SNHG17_38422086_38422241 | 0,07247848 | 2,878269746 |
| SE_SNHG17_38422091_38422241 | 0,089162304 | 2,737552611 |
| SE_SNHG17_38422095_38422241 | 0,074873043 | 2,978411832 |
| SE_MYO18A_29085603_29085648 | 0,002457144 | 5,713520812 |
| SE_CFDP1_75414577_75414695 | 0,006048537 | 0,231502533 |
| SE_PRR14_30650992_30651127 | 0,001307676 | 6,582979848 |
| SE_ERVK13_1_2672706_2672976 | 0,007410761 | 0,234607339 |
| SE_IL4R_27318634_27318767 | 3,04E-05 | 10,87192703 |
| SE_IDH3A_78150803_78150960 | 0,417732088 | 0,645926513 |
| SE_USP8_50484274_50484361 | 0,016710614 | 5,237738767 |
| SE_INF2_104695989_104696490 | 0,189121787 | 3,621940299 |
| SE_INF2_104715832_104716017 | 0,151558928 | 2,591743796 |
| SE_RDH11_67686083_67686257 | 0,163420011 | 2,305528646 |
| SE_SLC22A23_3276680_3276976 | 0,064445182 | 4,180478156 |
| SE_GUK1_228148370_228148539 | 0,001678977 | 0,144895509 |
| SE_ZNF37A_38094930_38095224 | 0,146779144 | 0,343844564 |
| SE_DCTD_182916418_182916539 | 0,012304529 | 0,245941556 |
| SE_SCARB2_76180210_76180354 | 0,023521835 | 4,168406194 |
| SE_ATP2C1_130894589_130894775 | 0,062641878 | 0,357984738 |
| SE_ATP2C1_130894730_130894775 | 0,013906306 | 4,824335457 |
| SE_ACO2_41477912_41478127 | 0,163452106 | 2,426475139 |
| SE_MLPH_237534563_237534647 | 0,120619355 | 0,399258503 |
| SE_DHRS12_51797825_51797908 | 0,003089549 | 4,539237482 |
| SE_LPP_188341662_188341719 | 0,022614126 | 3,428518699 |
| SE_UBA5_132660898_132661048 | 0,244503988 | 0,316697769 |
| SE_MYO19_36501851_36501961 | 0,074777946 | 0,377847407 |
| SE_SERPINB6_2966399_2966590 | 0,289098342 | 0,56916903 |
| SE_BIRC6_32435495_32435585 | 0,075187418 | 0,32994991 |
| SE_SRP9_225786861_225786985 | 0,001382746 | 7,966559379 |
| SE_WDR77_111441498_111441604 | 0,165942173 | 2,230312654 |
| SE_RTKN_74439550_74439977 | 0,532475967 | 1,399716635 |
| SE_FOXN3_89190386_89190449 | 0,105808398 | 4,614843155 |
| SE_NCOR1_16198600_16198719 | 0,07238608 | 3,69E-09 |
| SE_ZNF584_58409940_58410091 | 0,095862549 | 0,400400209 |
| SE_ITGB1BP1_9419989_9420100 | 0,287957353 | 1,87374473 |
| SE_ITGB1BP1_9419989_9420100_1 | 0,130927611 | 239062214,1 |
| SE_JPX_73946998_73947374 | 0,000839967 | 0,162217215 |
| SE_LIAS_39468118_39468427 | 0,12586758 | 2,631314738 |
| SE_RNF152_61890364_61890516 | 0,185166923 | 0,429857505 |

| | | |
|---|---|---|
| SE_SRSF4_29168502_29168638 | 0,020960241 | 3,753238585 |
| SE_SCMH1_41152598_41152741 | 0,012951784 | 4,80332361 |
| SE_MRRF_122286038_122286166 | 0,228056165 | 2,206793446 |
| SE_TPM1_63043705_63043831 | 0,0429981 | 3,139655485 |
| SE_CAPZB_19448833_19449330 | 0,018267342 | 5,718378688 |
| SE_OPA1_193617177_193617285 | 0,372715484 | 0,608073496 |
| SE_ACP1_272036_272065 | 0,000164976 | 0,106514934 |
| SE_RPS6KB2_67432355_67432492 | 0,076452622 | 0,317105986 |
| SE_TMC6_78120652_78120832 | 0,039143848 | 2,903502437 |
| SE_RAD51D_35116858_35117037 | 0,015454964 | 3,727510825 |
| SE_MTURN_30146176_30146299 | 0,030018981 | 0,143506778 |
| SE_SNHG14_25067408_25067531 | 0,006615367 | 0,100960771 |
| SE_SNHG14_25067411_25067531 | 0,011923923 | 0,239589876 |
| SE_KLHL2_165228806_165228913 | 0,096420593 | 0,336697975 |
| SE_SNX19_130880621_130880806 | 0,003245152 | 6,139215667 |
| SE_SMPD4_130159512_130159634 | 0,006740635 | 0,233965017 |
| SE_XRRA1_74859157_74859283 | 0,186880507 | 2,187993118 |
| SE_TRANK1_36850153_36851417 | 0,048461672 | 2,900326734 |
| SE_TNS1_217830366_217830390 | 0,088872136 | 2,537397337 |
| SE_MTERF4_241091894_241092114 | 0,053769429 | 3,655741537 |
| SE_ABHD14B_51971459_51971698 | 1,79E-06 | 17,69477782 |
| SE_CEP43_167027866_167027993 | 0,349351671 | 0,540308501 |
| SE_NMRAL1_4471313_4471449 | 3,33E-05 | 0,097944203 |
| SE_HOTAIRM1_27099073_27099366 | 0,017407995 | 3,724726676 |
| SE_HOTAIRM1_27099097_27099366 | 0,036005712 | 3,218702933 |
| SE_PHLDB1_118613293_118613450 | 0,266990086 | 0,541434085 |
| SE_SRSF11_70231094_70231203 | 0,000928049 | 9,436877908 |
| SE_FRYL_48710518_48710698 | 0,269497502 | 1,796867051 |
| SE_SDHAP1_195984491_195984737 | 0,102406171 | 0,411852229 |
| SE_AFG3L1P_89985445_89985570 | 0,001132883 | 7,603481238 |
| SE_RNF34_121404848_121405113 | 0,108015344 | 0,215589274 |
| SE_TRIM65_75891812_75891878 | 0,124880945 | 2,734039681 |
| SE_EMP1_13199177_13199533 | 0,009016411 | 6,579845196 |
| SE_PALM2AKAP2_109942685_109943381 | 0,008004469 | 0,188900501 |
| SE_PXN_120216272_120216581 | 0,269170646 | 0,518906257 |
| SE_BBS1_66523171_66523365 | 0,007260815 | 4,29272611 |
| SE_SEC24C_73762097_73762166 | 8,68E-06 | 14,0546776 |
| SE_SORBS1_95414493_95414655 | 0,00110698 | 8,918874644 |
| SE_FRMD4A_13759164_13759506 | 0,012418339 | 4,878407888 |
| SE_WASF3_26680034_26680201 | 0,048155039 | 0,328414226 |
| SE_MYCBP2_77118339_77118519 | 0,089446856 | 2,485310074 |
| SE_MCM9_118932606_118932740 | 0,025126672 | 0,123578762 |
| SE_RANGAP1_41284762_41285153 | 0,124410082 | 2,692043104 |
| SE_TBC1D10A_30293158_30293453 | 0,296622384 | 2,836607844 |
| SE_BCAT2_48807768_48808236 | 0,054876255 | 0,362959262 |
| SE_RTF2_56516934_56516987 | 0,14285167 | 0,241718805 |
| SE_RTF2_56516943_56516989 | 0,024587865 | 0,207454895 |
| SE_GIPC1_14482688_14482819 | 1,81E-05 | 11,41872643 |
| SE_CFDP1_75414577_75414694 | 0,006048537 | 0,231502533 |
| SE_LYRM1_20915555_20915713 | 0,183872325 | 2,611851915 |
| SE_LYRM1_20915555_20916047 | 0,183872325 | 2,611851915 |
| SE_ABHD2_89103902_89104299 | 0,082370455 | 3,281334571 |
| SE_SERPINA1_94388566_94389010 | 0,036399412 | 3,384224292 |

| | | |
|---|---|---|
| SE_MIA2_39320929_39321056 | 0,001364009 | 7,420725658 |
| SE_DCAF11_24115203_24115293 | 0,144808783 | 4,138098657 |
| SE_GUK1_228148233_228148455 | 0,001678977 | 0,144895509 |
| SE_GUK1_228148357_228148455 | 0,001678977 | 0,144895509 |
| SE_GUK1_228148357_228148539 | 0,001678977 | 0,144895509 |
| SE_XPNPEP1_109893011_109893073 | 0,119521768 | 2,742397105 |
| SE_PARG_49922546_49922665 | 0,165241147 | 0,461807602 |
| SE_RBM47_40436440_40436644 | 0,471232365 | 1,623314386 |
| SE_LPP_188341655_188341719 | 0,049768086 | 2,85621862 |
| SE_ZNF3_100079539_100079656 | 0,461220906 | 0,570924241 |
| SE_SNHG8_118278965_118279137 | 0,17951058 | 0,4539815 |
| SE_FCHSD1_141647967_141648426 | 0,054676447 | 4,381508145 |
| SE_NCOR1_16198507_16198719 | 0,07238608 | 3,69E-09 |
| SE_SUMF1_4362038_4362254 | 0,030463393 | 3,934742737 |
| SE_ITGB1BP1_9419989_9420418 | 0,157586874 | 2,499426399 |
| SE_SRSF7_38749239_38749346 | 0,007396682 | 4,896414089 |
| SE_HNRNPA1_54282843_54283225 | 0,011830445 | 4,500370049 |
| SE_HNRNPA1_54282843_54283228 | 0,011830445 | 4,500370049 |
| SE_HNRNPA1_54282843_54283234 | 0,011830445 | 4,500370049 |
| SE_SLC4A7_27379005_27379356 | 0,544889385 | 1,408119105 |
| SE_CAMKK2_121248605_121248861 | 0,130475119 | 0,447187328 |
| SE_PCCA_100305750_100305834 | 0,000468982 | 0,120162952 |
| SE_ARIH2_48922327_48922811 | 0,18858554 | 0,360552714 |
| SE_LPGAT1_211830182_211830326 | 0,093861547 | 2,567964697 |
| SE_SNX19_130880621_130880796 | 0,001466192 | 6,430523408 |
| SE_SNX19_130880621_130880805 | 0,001466192 | 6,430523408 |
| SE_TRIM66_8654433_8654578 | 0,033398404 | 4,322065158 |
| SE_SERGEF_17959469_17959634 | 0,510454785 | 1,473746357 |
| SE_DHPS_12677143_12677211 | 0,115739915 | 2,487101374 |
| SE_AL357075_4_154441291_154441347 | 0,252396911 | 0,531337614 |
| SE_RNF216_5741747_5741815 | 0,027969185 | 296220050,2 |
| SE_AC010522_1_57840837_57840908 | 0,497466626 | 1,494488444 |
| SE_AC010522_1_57840837_57840927 | 0,497466626 | 1,494488444 |
| SE_KIF1B_10325899_10326359 | 0,267598051 | 1,925450701 |
| SE_TMEM176A_150804372_150804485 | 0,099488838 | 0,292386577 |
| SE_TMEM176A_150804372_150804719 | 0,056915173 | 0,256823012 |
| SE_SRSF11_70231094_70231257 | 0,000167285 | 8,456066695 |
| SE_AFG3L1P_89978472_89978878 | 0,268747288 | 2,061783419 |
| SE_MGA_41742545_41742724 | 0,135529757 | 0,402897955 |
| SE_RPL8_144791427_144791872 | 0,309389072 | 1,24E-08 |
| SE_KRT8_52948722_52948828 | 0,095160365 | 2,523318105 |
| SE_KRT8_52948722_52949218 | 0,127264145 | 2,329192396 |
| SE_TMEM141_136791629_136791777 | 0,001531495 | 5,741714425 |
| SE_DGLUCY_91223675_91223795 | 0,091455256 | 3,726860096 |
| SE_BPTF_67875556_67875744 | 0,020469609 | 0,190554167 |
| SE_CTNND1_57791491_57791701 | 0,060686922 | 2,979659426 |
| SE_CTNND1_57791493_57791701 | 0,060686922 | 2,979659426 |
| SE_RBM4_66652276_66652452 | 0,074263305 | 0,177589794 |
| SE_NDUFB1_92121195_92121413 | 0,046145728 | 4,59008161 |
| SE_MAPK6_52042685_52042833 | 0,192235461 | 0,475040644 |
| SE_SORBS1_95414498_95414862 | 0,007143223 | 4,703724142 |
| SE_LTBP4_40622367_40622667 | 0,004331827 | 0,180907868 |
| SE_LTBP4_40622384_40622667 | 0,004331827 | 0,180907868 |

| | | |
|---|---|---|
| SE_LTBP4_40622400_40622613 | 0,031235295 | 0,307915031 |
| SE_MYCBP2_77118339_77118495 | 0,171219605 | 2,115928066 |

**Table S5. Differentially spliced genes in common in both cohorts.**

| Inclusion events in common | Exclusion events in common | |
|---|---|---|
| *MYCBP2* | *LAS1L* | *GTF2IRD1* |
| *GTF2IRD1* | *CLEC16A* | *YBX3* |
| *SBNO2* | *SBNO2* | *PTPN18* |
| *NFKB2* | *KDM2B* | *SORBS1* |
| *TCF20* | *MYO9B* | *MAPK1* |
| *FUBP3* | *FMR1* | *ZNF106* |
| *DNAH1* | *CAMTA2* | *MED13* |
| *BIRC6* | *CAMKK2* | *DNAH1* |
| *SRSF4* | *GLS* | *BIRC6* |
| *ABCC10* | *PCID2* | *SMARCA4* |
| *PCID2* | *KDM4B* | *BAIAP2L2* |
| *MAP2K2* | *INO80* | *COL5A1* |
| *KDM4B* | *LLGL1* | *CC2D1A* |
| *LLGL1* | *WASF3* | *ITGA7* |
| *USP8* | *POLG* | *NCOR1* |
| *POLG* | *RANBP10* | *CDC42BPA* |
| *CDC42BPA* | *LPP* | *FRMD4A* |
| *BPTF* | *EPG5* | *PABIR2* |
| *WDR27* | *ATXN7L2* | *NCSTN* |
| *ABCF1* | *HPS3* | *TSC1* |
| *LINC00894* | *SLC39A13* | *TRIM66* |
| | *STAT3* | *CHD3* |
| | *BPTF* | *EXOSC10* |
| | *PCCA* | *SEC24C* |
| | *RBM10* | *WDR27* |
| | *POFUT2* | *OPA1* |
| | *INF2* | *ABCF1* |
| | *MYO15B* | |
